# Benchmarking atlas-level data integration in single-cell genomics

**DOI:** 10.1101/2020.05.22.111161

**Authors:** MD Luecken, M Büttner, K Chaichoompu, A Danese, M Interlandi, MF Mueller, DC Strobl, L Zappia, M Dugas, M Colomé-Tatché, FJ Theis

## Abstract

Cell atlases often include samples that span locations, labs, and conditions, leading to complex, nested batch effects in data. Thus, joint analysis of atlas datasets requires reliable data integration.

Choosing a data integration method is a challenge due to the difficulty of defining integration success. Here, we benchmark 38 method and preprocessing combinations on 77 batches of gene expression, chromatin accessibility, and simulation data from 23 publications, altogether representing >1.2 million cells distributed in nine atlas-level integration tasks. Our integration tasks span several common sources of variation such as individuals, species, and experimental labs. We evaluate methods according to scalability, usability, and their ability to remove batch effects while retaining biological variation.

Using 14 evaluation metrics, we find that highly variable gene selection improves the performance of data integration methods, whereas scaling pushes methods to prioritize batch removal over conservation of biological variation. Overall, BBKNN, Scanorama, and scVI perform well, particularly on complex integration tasks; Seurat v3 performs well on simpler tasks with distinct biological signals; and methods that prioritize batch removal perform best for ATAC-seq data integration. Our freely available reproducible python module can be used to identify optimal data integration methods for new data, benchmark new methods, and improve method development.

## Introduction

The complexity of single-cell omics datasets is increasing. Current datasets often include many samples^1^, generated across multiple conditions^2^, with the involvement of multiple labs^3^. Such complexity, which is common in maps of specific tissues and organs or whole reference atlas initiatives such as the Human Cell Atlas^4^, creates inevitable batch effects. Therefore, the development of data integration methods that overcome the complex, nonlinear, nested batch effects in these data has become a priority. Indeed, data integration has been described as one of the grand challenges of scRNA-seq data analysis^5,6^.

Batch effects represent unwanted technical variation in the data that affects groups (or batches) of cells. Batch effects can arise from variations in sequencing depth, sequencing lanes, read length, plates or flow cells, protocol, experimental labs, sample acquisition and handling, sample composition, reagents or media, and/or sampling time. Furthermore, biological factors such as tissues, spatial locations, species, time points, or inter-individual variation can also be regarded as a batch effect under certain circumstances.

Appropriate data integration methods are required to deal with these batch effects. Here, we define single-cell data integration as the process of combining datasets or samples of high-throughput sequencing data to produce a self-consistent version of the data for downstream analysis^7^. The output of these methods is either an integrated graph, a joint embedding, or a corrected feature space. Importantly, we distinguish data integration from batch correction according to method complexity, i.e., the complexity of the batch effect that can be removed. Whereas batch removal is typically used to integrate samples from the same lab and/or experiment, data integration should be applied to tasks involving nested batch effects from, for example, multiple labs and/or protocols.

Currently, 31 integration methods for scRNA-seq data are available^8^ (as of February 2020; **Supplementary Table 1**). Consequently, when confronted with a new data integration problem, analysts face the difficult decision of choosing a particular method. Moreover, it is difficult to envisage how an integrated dataset should look; thus, integration method choice can be biased by the subjective opinion of the analyst. Benchmarking integration methods can help solve this problem and provide an unbiased guide to method choice.

Previous studies on benchmarking methods for data integration have focused on the simpler problem of batch effect removal in scRNA-seq^9,10^. These studies benchmarked methods on simple integration tasks with low batch complexity and found that ComBat^9^ or the linear, principal component analysis (PCA)-based, Harmony method^10^ outperformed more complex, nonlinear, methods.

Here, we present the first benchmarking study in which the performance of data integration methods in complex integration tasks (such as those now commonly required in the analysis of tissue and organ atlases) is investigated. Specifically, we benchmark 10 popular data integration tools on nine data integration tasks consisting of up to 23 batches and 1 million cells, for both scRNA- and scATAC-seq data. We selected eight single-cell data integration tools [matching mutual nearest neighbors (MNN)^11^, Seurat v3^12^, scVI^13^, Scanorama^14^, batch-balanced k-nearest neighbors (BBKNN)^15^, LIGER^16^, clustering on network of samples (Conos)^17^, and Harmony^18^], a bulk data integration tool (ComBat^19^), and a perturbation modeling tool [transformer variational autoencoder (trVAE)^20^]. Moreover, we use 14 metrics to evaluate the integration methods on their ability to remove batch effects while conserving biological variation. We focus in particular on assessing the conservation of biological variation beyond cell identity labels, e.g., we assess the conservation of trajectories or cell cycle effects via novel integration metrics. Our methodology allows us to adequately assess the strengths and limitations of nonlinear methods, which have become necessary in the atlas-level integration tasks increasingly faced by the data analysis community. We find that BBKNN, Scanorama, and scVI perform well, particularly on complex integration tasks. In addition, Seurat v3 performs well on simpler tasks with distinct biological signals, and Harmony and scVI are partially effective for scATAC-seq data integration.

## Results

### Single-cell integration benchmarking (scIB)

We benchmarked 10 popular data integration methods on nine preprocessed integration tasks: two simulation tasks, five RNA-seq tasks, and two ATAC-seq tasks (**Fig. 1**). Each task posed a unique challenge (e.g., nested batch effects caused by protocols and donors, batch effects in a different data modality, and scalability up to 1 million cells) that revolved around integrating data on a particular tissue from multiple labs (**Table 1**). These real data represent complex, nested batch-effect scenarios; therefore, careful assessment of the “ground truth” is required. Our simulation tasks allowed us to assess the integration methods in a setting where the nature of the batch effect could be determined and the ground truth is known. We predetermined this ground truth by preprocessing and annotating real data from 23 publications separately for each batch (see **Methods**).

**Table 1:** Integration tasks for benchmarking. Overview of the tasks used to benchmark data integration methods. The tested feature describes the unique challenge presented by the integration task. *Donor* refers to human individuals, *sample* is used when mice are involved, and *batches* is the general term that includes dataset and sample batches.

| Integration task | Cell number | Batches | Tested features |
| --- | --- | --- | --- |
| Pancreas | 16,382 | 9 batches | Widely used test data, protocols |
| Lung | 32,472 | 16 donors | Human variation, protocols, spatial locations, high resolution subtypes, labs |
| Immune (human) | 33,506 | 10 donors | Tissues, labs, similar cell types |
| Immune (human & mouse) | 97,952 | 23 samples | Tissues, labs, similar cell types, species |
| Mouse brain (RNA) | 978,734 | 4 datasets | Large dataset, spatial locations, nucleus vs cell |
| Mouse brain small (ATAC) | 25,960 | 3 datasets | Different modality |
| Mouse brain large (ATAC) | 67,612 | 3 datasets | Different modality, unbalanced batches |
| Simulation 1 | 12,097 | 6 batches | Variation in cellular compositions |
| Simulation 2 | 19,318 | 16 batches | Nested batch effects, composition variation |

**Figure 1:**
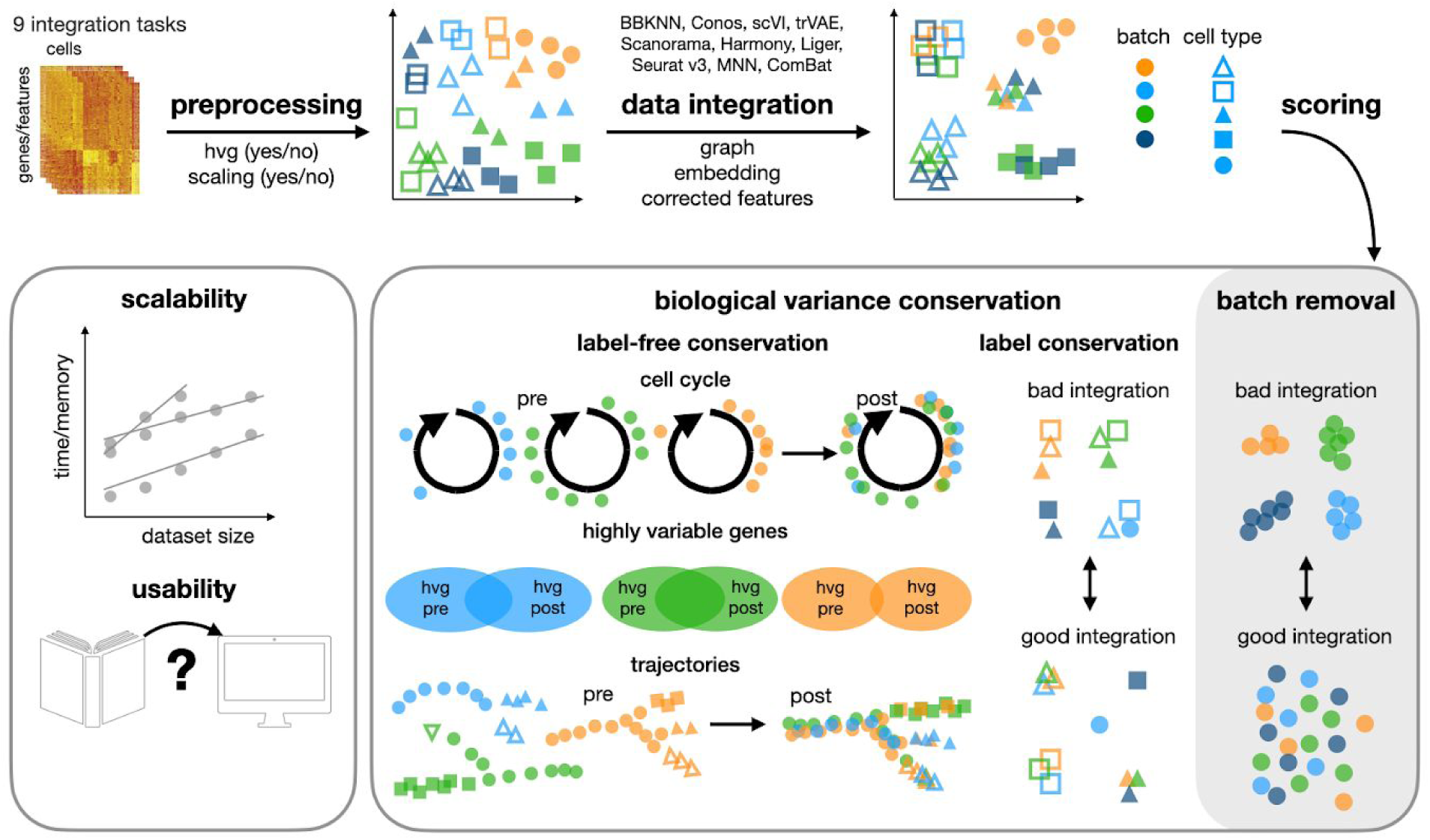
Design of single-cell integration benchmarking (scIB). Schematic diagram of the benchmarking workflow. Here, 10 data integration methods with four preprocessing decisions are tested on nine integration tasks. Integration results are evaluated using 14 metrics that assess batch removal, conservation of biological variance from cell identity labels (label conservation), and conservation of biological variance beyond labels (label-free conservation). The scalability and usability of the methods are also evaluated.

Each integration method was evaluated with regards to accuracy, usability, and scalability (see **Methods**). Integration accuracy was evaluated using 14 performance metrics divided into two categories that typically oppose each other: removal of batch effects and conservation of biological variance (**Fig. 1**). Batch effect removal per cell identity label was measured via the k-nearest neighbor batch effect test (kBET)^21^, kNN graph connectivity, and the Average Silhouette Width (ASW)^21^ across batches. Independently of cell identity labels, we further measured batch removal using the graph integration Local Inverse Simpson’s Index (graph iLISI, extended from iLISI^18^) and PCA regression^21^. Conservation of biological variation in single-cell data can be captured at the scale of cell identity labels (label conservation) and beyond this level of annotation (i.e., label-free conservation). Therefore, we used both classical label conservation metrics [assessed using local neighborhoods (graph cLISI, extended from cLISI^18^), global cluster matching (Adjusted Rand Index^22^, Normalized Mutual Information^23^), relative distances (cell type ASW), and two novel metrics evaluating rare cell identity annotations (isolated label scores)] and three novel label-free conservation metrics: (1) cell cycle variance conservation, (2) overlaps of highly variable genes (HVGs) per batch before and after integration, and (3) conservation of trajectories (see **Methods**).

The diversity in output formats from data integration methods poses a challenge to fair benchmarking^24^. Although input data are consistently preprocessed, requirements on scaling and HVG selection also differ between methods. We addressed these challenges in three ways. Firstly, all integration outputs were treated as separate integration runs. For example, Scanorama outputs both corrected expression matrices and embeddings; these are evaluated as two separate outputs (Scanorama gene and Scanorama embedding). Secondly, we developed novel extensions to kBET and LISI scores that worked on graph-based outputs, joint embeddings, and corrected data matrices in a consistent manner (**Supplementary Notes 1** and **2**). For instance, we sped up graph LISI scoring via a fast, parallel C++ implementation that scales to millions of cells. Thus, multiple metrics can be computed for each category of batch effect removal, label conservation, and label-free conservation (**Supplementary Table 2**). Overall accuracy scores were computed by taking the weighted mean of all metrics computed for an integration run, with a 40:60 weighting of batch effect removal to biological variance conservation (bio-conservation) irrespective of the number of metrics computed. Thirdly, while we ran each method according to defaults provided by the authors (see **Methods**) and contacted them if errors were encountered, we also included preprocessing decisions in our benchmark to assess whether scaling or HVG selection improves output. We considered that some methods cannot accept scaled input data (i.e., LIGER, trVAE, and scVI). Thus, we tested 38 data integration setups per integration task, resulting in 342 attempted integration runs. All performance metrics, integration methods with parameterizations, and preprocessing functions have been made available in our *scIB* python module. Furthermore, our workflow is provided as a reproducible Snakemake^25^ pipeline to allow users to test and evaluate data integration methods in their own setting.

### Data integration benchmarking exemplified with human immune cells

To demonstrate our evaluation of data integration methods, we first focus on the human immune cell integration task (**Fig. 2a** and **Supplementary Note 3.1.1**). This task comprises 10 batches representing donors from five datasets with cells from peripheral blood and bone marrow. All integration methods successfully completed this task without exceeding time and memory limitations. With particular preprocessing, Scanorama (using joint embeddings), Conos, Harmony, and BBKNN performed well. By considering the embedded data plots of the integration results (**Fig. 2b,c**), it is possible to understand how method performance rankings were obtained.

**Figure 2:**
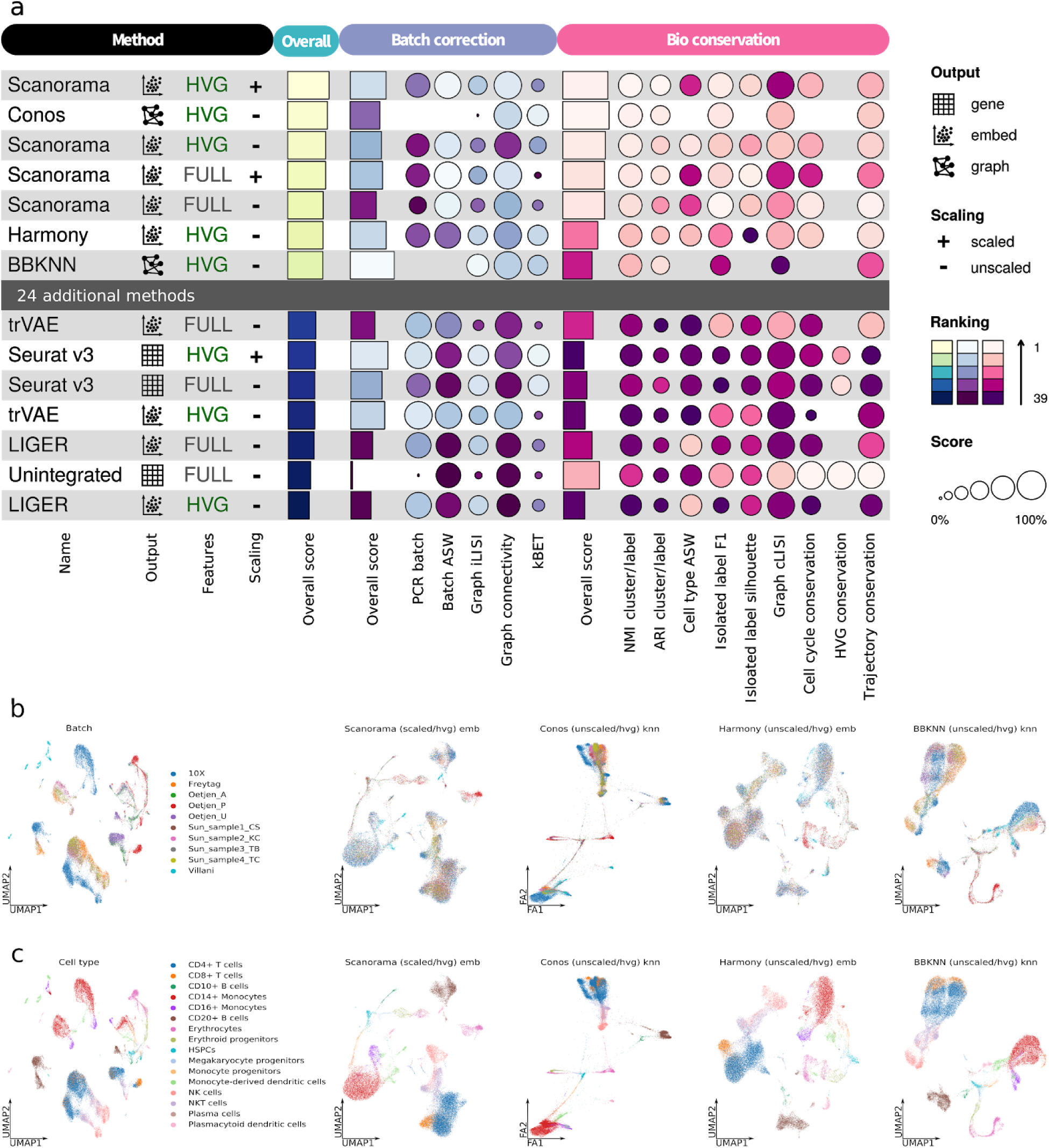
Benchmarking results for the human immune cell task. (a) Overview of top and bottom ranked methods by overall score for the human immune cell task. Metrics are divided into batch correction and bio-conservation (pink) categories. Overall scores are computed using a 40:60 weighted mean of these category scores (see **Methods** for further visualization details and **Supplementary Fig. 4** for the full plot). (b and c) Visualization of the best performers on the human immune cell integration task colored by batch (b) and cell identity annotation (c). The plots show Force Atlas 2 (Conos) and UMAP (all other methods) layouts for the unintegrated data (left), and the top four performers (right).

All high-performing methods succeeded in removing batch effects between individuals and platforms while conserving biological variation at the cell type and subtype levels; this is reflected in their relatively high batch removal and bio conservation scores. In comparison to other top-performing methods, Conos had a lower batch removal score principally due to a low graph iLISI score. Consequently, batch structure was found within the CD4+ T cell cluster, and there was a closer proximity between the Smart-seq2 clusters from Villani *et al*.^*26*^ in the Conos output. In contrast, BBKNN exhibited a lower bio-conservation compared with its batch removal score due to a lower isolated label F1 score. The isolated labels in this task were *CD10*+ *B cells, erythroid progenitors (EPs), erythrocytes*, and *megakaryocyte progenitors (MPs)*, which are exclusive to Oetjen *et al*.^*27*^ batches. BBKNN separated the *MPs* into two populations independently of their batch, leading to a low F1 score. In contrast, Harmony kept each isolated cell label together, but showed an overlap between these populations (specifically *EPs, MPs*, erythrocytes, and monocyte-derived dendritic cells), leading to a comparatively low isolated label ASW score but a high isolated label F1 score.

We also focused on the conservation of trajectories. In this integration task, we assessed erythrocyte development from hematopoietic stem and progenitor cells (HSPCs) via MPs and EPs to erythrocytes (**Supplementary Fig. 1-3**). All of the top performing methods exhibited high trajectory conservation scores, whereas LIGER and Seurat v3, produced poor conservation of this trajectory: LIGER lost most of the trajectory structure beyond HSPCs and MPs and Seurat v3 appeared to place the cell types in broadly the correct order in a UMAP, but did not reflect this order in diffusion map space, in which a branching structure was produced (**Supplementary Fig. 2**).

### The trade-off between batch removal and conserving biological variation

Considering the results of the five RNA-seq and two simulation tasks (**Supplementary Note 3** and **Supplementary Fig. 4,6-18**), we found that the varying complexity of tasks affects the ranking of integration methods. For example, Seurat v3 and Harmony perform well on simulations, whereas BBKNN, Scanorama, and scVI tend to perform better on more complex real data. In general, the simulations contain less nuanced biological variation but exhibit clearly defined, often strong, batch effects. Specifically, simulation task 1 posed little difficulty to most methods independent of preprocessing decisions (**Supplementary Note 3.2**). Similar to the simulation scenarios, the widely used pancreas integration task contains distinct cell type variation and batch effects; thus even methods which perform poorly overall, performed well on this task (**Supplementary Fig. 9** and **16**, and **Supplementary Note 3.4**).

Particularly in more complex integration tasks we observed a trade-off between batch effect removal and bio-conservation (**Fig. 3a** and **Supplementary Fig. 19**). While methods such as BBKNN and Seurat v3 tend to favor the removal of batch effects over conservation of biological variation, Scanorama and Conos make the opposite choice. This trade-off is particularly noticeable where biological and batch effects overlap, such as in the lung atlas task. In this task, three datasets sample two distinct spatial locations (the airways and parenchyma). Particular cell types such as endothelial cells perform different functions in these locations (e.g., gas exchange in the parenchyma). While Seurat v3 integrates across the locations to merge these cells, thereby providing a broad cell type overview, Scanorama preserves the spatial variation in endothelial cells and other cell types that have functional differences across locations (**Supplementary Note 3.5**).

**Figure 3:**
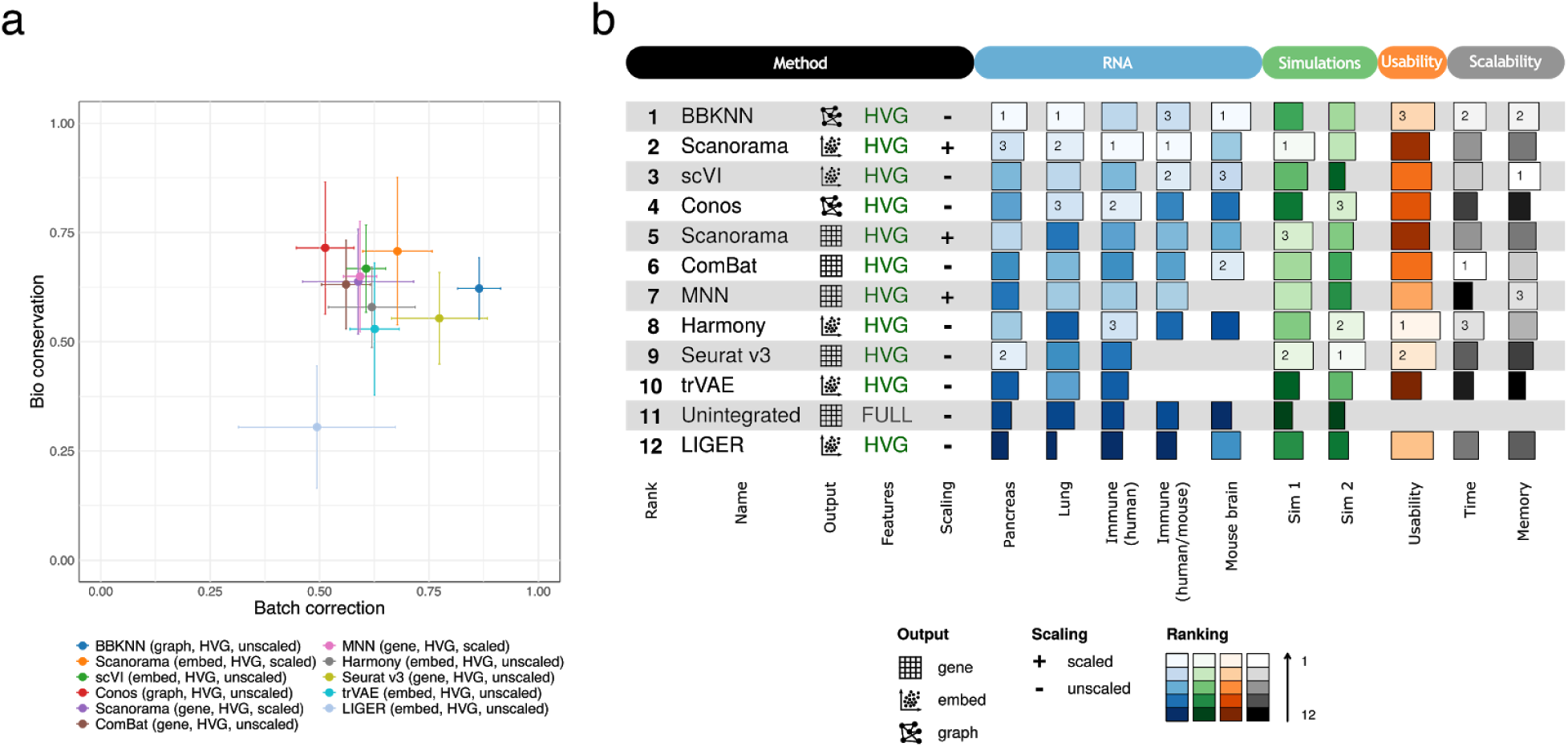
Overview of benchmarking results on all RNA integration tasks and simulations, including usability and scalability results. (a) Scatter plot of the mean overall batch correction score against mean overall bio-conservation score for the selected methods on RNA tasks. Error bars indicate one standard deviation. (b) The overall scores for the best performing methods on each task as well as their usability and scalability. Methods that failed to run for a particular task were assigned the unintegrated ranking for that task.

Where methods were focused on the removal of strong batch effects, we found that they often lost nuanced biological variation in cell subtypes or states. The most challenging batch effects across the integration tasks were due to species, sampling locations, single-nucleus vs single-cell data, and integration of microwell-seq data from the mouse cell atlas (MCA; **Supplementary Note 3**). Interestingly, the strongest batch effect contributors tended to also be interpretable as biological signals rather than technical noise. While the top performing methods across the integration tasks were largely unable to integrate across these effects (**Supplementary Fig. 13,17,18**), LIGER and Seurat v3 were successful. These integration results, however, are often generated by the bottom four performers because biological variation is also removed with the batch effect. This effect was particularly noticeable for the immune cell human/mouse and mouse brain tasks. For immune cells, only LIGER and Conos integrated across species (Seurat v3 failed to run on this task). While Conos removed the majority of variation in the integration process, LIGER integrated across species while retaining broad cell type variation. Nevertheless, LIGER also merged smaller cell labels (e.g., neutrophils and monocytes), created heterogeneous larger clusters, and removed the trajectory structure (**Supplementary Fig. 5,13** and **Supplementary Note 3.1.2**). Two exceptions are Scanorama and scVI, which integrated mouse brain data from single nuclei and single cells while retaining biological variation on spatial locations and rare cell types (**Supplementary Note 3.6**).

We found that methods that favor bio-conservation tended to perform better on label-free metrics. Indeed, Scanorama, ComBat, and MNN consistently perform well at conserving cell cycle variance and trajectory structure in the integrated data, whereas scVI, LIGER, Harmony, and Seurat v3 perform poorly. This effect is particularly notable from our trajectory results (**Supplementary Fig. 3-6** and **Supplementary Data 1**). For human immune cell data, the batch effect is comparatively small as the cells that form the trajectory originate from one dataset; thus Scanorama, ComBat, and MNN placed cells in the expected order per batch. These methods, and scVI (which successfully merged MCA and Dahlin *et al*.^*28*^ bone marrow data), also performed well per batch in the human/mouse immune cell task, but their results contained individual clusters as outliers, and human and mouse erythrocyte development was not integrated into a single trajectory; thus, while local trajectory structure was well-represented, the global trajectory structure was not conserved. Even LIGER, which integrated datasets across species, poorly reflected the trajectory. Overall, performing an integrated trajectory across species is challenging due to the strong species batch effect as well as cell and cluster outliers, for which integration was performed suboptimally.

### Scaling improves batch removal but impairs bio-conservation performance

Given the lack of best-practice for preprocessing raw data for data integration, we assessed whether integration methods perform better with HVG selection or scaling. We ran every integration method with four preprocessing combinations (see **Methods**), and compared the performance between runs that only differed in one preprocessing parameter. Across RNA and simulation tasks, HVG selection generally outperformed data integration of the full gene set: for HVGs, 72% of comparisons had a higher overall score; 80% had better batch removal; and 60% had better bio-conservation scores. Notable exceptions are trajectory and cell cycle conservation scores, which tended to favor full feature integration runs.

We also found that whether or not a method performs better with prior scaling depends on the method of choice (**Fig. 3b**). Independent of the method, scaling resulted in higher batch removal scores (63% of comparisons) but lower bio-conservation (72% of comparisons). This observation is consistent with unscaled data performing better in our label-free conservation metrics. A notable exception is the trajectory conservation metric in the presence of strong batch effects (i.e., between species in the human/mouse task); such trajectories are better captured with removal of the strong batch effect (i.e., with scaling).

### BBKNN, Scanorama, and scVI perform best overall for RNA-seq integration

To evaluate overall performance of data integration methods across RNA-seq and simulation scenarios, methods can be ranked by their overall scores. We assumed that there was a single, optimal way in which to run an integration method, and therefore ranked methods by their top performing preprocessing combination. Consequently, we also obtained an optimal way in which to run each integration method (**Fig. 3b**). The optimal preprocessing combinations of BBKNN, Scanorama, trVAE, and scVI were consistent across tasks. Conos, which incorporates HVG selection and scaling within its method, performed better with HVG selection on unscaled data but, for simpler tasks, performed better with full gene sets on scaled data. The performance of MNN was similar on unscaled and scaled data, while ComBat performed similarly with HVGs and full gene sets. Interestingly, LIGER performed better with HVG sets and unscaled data overall, but it performed slightly better in 4 of 7 tasks with full gene sets rather than HVGs. In contrast, the performance of Seurat v3 and Harmony was not consistently better with a particular preprocessing combination, although preprocessing did affect their performance across tasks.

Given that the complexity of a task affects the appropriateness of a method, we ranked methods based on real data tasks that better represent the challenges typically faced by analysts. Overall, we found that the graph-based method BBKNN, and the embeddings output by Scanorama and scVI, perform best, whereas LIGER performs poorly. These results are remarkably consistent across tasks for integrating real data. However, Seurat v3 and Harmony, which usually rank outside the top third of methods for real data, are favorable for simulations. The methods with a higher level of abstraction tended to rank higher. This was particularly noticeable when comparing Scanorama embeddings and Scanorama’s corrected expression matrix output. Likewise, integrated graph methods tended to perform well; however, only a subset of metrics can be run on their outputs, so their results may be less robust. Autoencoder-based frameworks such as scVI and trVAE tended to perform better in tasks with more cells and complex batch structure. This was particularly noticeable for scVI, as trVAE did not scale to tasks of this size without GPU hardware.

### Scalability and usability

We assessed the scalability of each data integration method by monitoring the CPU time and peak memory use reported by our Snakemake pipeline (**Supplementary Fig. 20**). As expected, using the full feature matrix led to both longer runtimes and higher memory usage compared to selecting a fixed set of HVGs. In contrast, data scaling had little influence on CPU time, while peak memory use was increased in the scaled data scenario due to reduced sparsity upon scaling. In particular, Conos used considerably more memory with scaled data. For unscaled data, the memory usage of scVI was superior to other methods, while BBKNN and ComBat performed best in terms of runtime. For scaled data, the memory usage of BBKNN was superior, while the runtime of ComBat was slightly favorable. However, only BBKNN worked successfully for all datasets and all preprocessing combinations. Given the runtime and memory limitations during the benchmarking setup (see **Methods**), trVAE could not integrate datasets with >67,000 cells, while Seurat v3 failed to integrate datasets >100,000 cells. Overall, Conos had the highest memory requirements, but it succeeded in integrating 1 million cells without prior scaling. Furthermore, MNN used most CPU time, but its memory usage hardly increased with increasing cell numbers for a fixed number of HVGs.

We assessed the usability of methods according to criteria previously applied to evaluate the usability of trajectory inference methods^29^ (see **Methods** and **Supplementary Fig. 21**). Most of the methods are easy to use because of tutorials, function documentation, and open source code. However, the robustness of method performance and the accuracy quantification on real and simulated data differ between published methods. Overall, Harmony, BBKNN, and Seurat v3 have the best usability for new users. In contrast, Conos, Scanorama, and trVAE are somewhat lacking in usability as they lack function documentation or high-quality tutorials.

### scATAC-seq batch effects require strong batch correction

Several of the ten benchmarked data integration methods have been used to integrate datasets across modalities^12,16^. With the growing availability of datasets, removing batch effects within scATAC-seq data is also becoming an application of interest. As the integration challenge is similar, we asked whether method performance transfers to scATAC-seq data.

We used non-overlapping sliding windows as the canonical, unbiased unit for processing open chromatin data and as a basis for data integration. We evaluated the performance of the ten integration methods on two scATAC-seq tasks (**Table 1**). Both tasks involve integration of cells from the same three datasets. While the large ATAC task contains more samples and cells from the dominant batch (ratio of cells between datasets = 5:20:75), the small ATAC task contains a more balanced batch composition (ratio of cells between datasets = 13:57:30; **Supplementary Data 2**). To restrict the feature space, we used only the most highly variable windows that overlap between datasets (see **Methods**). This posed an ATAC-specific challenge, as integration of more batches and cells leads to a lower number of shared informative windows between datasets (**Supplementary Note 3.7**). Despite this large reduction of the feature space, scaling to >50,000 cells became a challenge; trVAE and LIGER failed to run on the large ATAC task, while MNN failed in both tasks due to its poor scalability with the number of cells and features. In contrast, MNN could be evaluated on scRNA-seq integration tasks of ≤100,000 cells. Using a higher number of shared features between datasets would increase these scalability problems.

In general, most of the methods performed poorly for batch correction in both ATAC tasks (**Fig. 4** and **Supplementary Fig. 22**). This may be attributable to the binary nature of the scATAC-seq input data; the benchmarked methods were designed for gene expression counts with a range of expression values. Furthermore, high bio-conservation scores were often mediated by high silhouette scores, which measured compact, often unintegrated, cell type clusters. We would therefore recommend to prioritize batch correction over biological conservation for ATAC integration. BBKNN, Harmony, and scVI were the top three performers for batch integration (**Fig. 4a**). BBKNN removed batch effects at the expense of a strong loss of biological conservation. With more compact (but partially unintegrated) cell identity clusters, Harmony’s bio-conservation score was higher than that of BBKNN. Finally, scVI showed a compromise between good batch correction and moderate bio-conservation. Seurat v3 and ComBat instead ranked top for biological conservation as they exhibited compact, but unintegrated clustering of cell types and thus ranked only fourth and fifth for batch correction (**Supplementary Fig. 22**).

**Figure 4:**
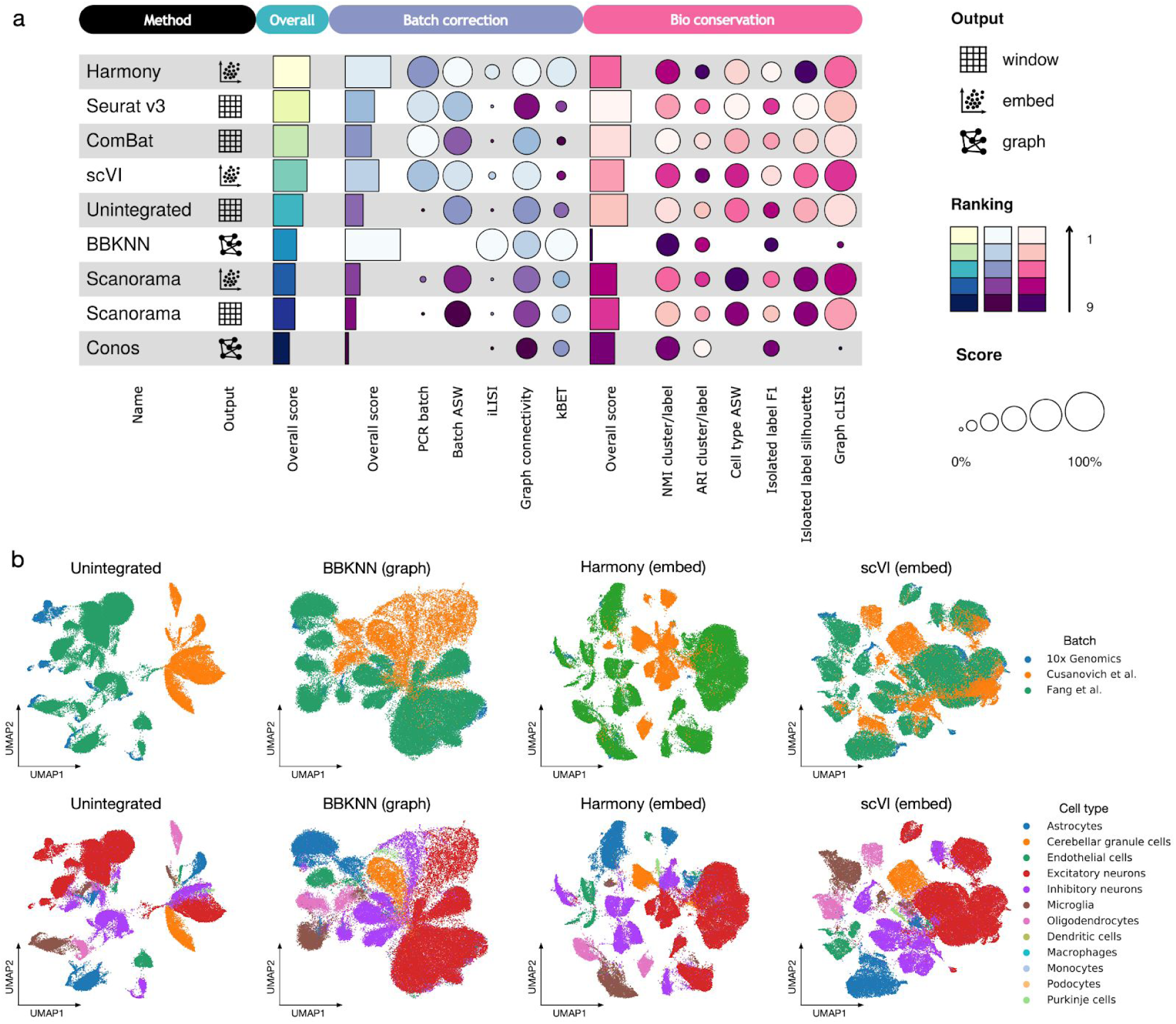
Benchmarking results for the large mouse brain ATAC task. (a) Benchmarking result for the large ATAC task. Methods that failed to run due to time or memory limitations are not shown. (b) Visualization of the best batch correction methods on the large ATAC task coloured by batch labels (top row) and cell identity annotation (bottom row). The plots show UMAP layouts for the unintegrated data, and the top three performers based on the average of batch correction scores from both ATAC tasks in descending order.

In general, stronger batch effects were found between datasets that shared fewer informative windows (**Supplementary Note 3.7**). Furthermore, batch imbalance notably affected Seurat v3, which was likely because it integrated datasets in a different order in the two tasks (**Supplementary Figs. 23** and **24**).

Overall, in the two ATAC tasks we conclude that the best batch removal methods are BBKNN, Harmony, and scVI, with different performance for bio-conservation (**Fig. 4a**). However, all methods perform inadequately: most batches remain separated in low dimensional visualizations of the integrated data (**Fig. 4b**).

## Discussion

We benchmarked ten integration methods with four pre-processing combinations on nine integration tasks consisting of scRNA-seq, scATAC-seq, and simulated data. Method evaluation was performed on the basis of usability, scalability, and integration performance via 14 metrics that measure trade-offs between batch integration and conservation of biological variance. Overall, we observed that method performance is dependent on the complexity of the integration task for RNA and simulation scenarios. For example, the use of Seurat v3 and Harmony is appropriate for simple integration tasks with distinct batch and biological structure; however, these methods typically rank outside the top three when used for complex real data scenarios, which is in agreement with recent benchmarks on simpler batch structures^10,24^. In contrast, on more complex integration tasks, BBKNN, Scanorama (embeddings), and scVI performed well.

Our overall rankings were based on metrics measuring different aspects of integration success. For example, while certain bio-conservation metrics prioritized clearly separated cell clusters, others favored continuous cellular structures such as trajectory and cell cycle conservation. Furthermore, metric usage depends on data output type. Even for the integrated graph outputs generated by Conos and BBKNN, it was possible to measure three batch removal and three bio-conservation metrics (**Supplementary Table 2**). Such metric diversity ensures that no individual method only performs well because of the optimization of a single metric, e.g., BBKNN, for which the underlying optimization function is similar to the graph iLISI metric (and therefore it also receives lower graph cLISI scores). Irrespective of the number of metrics used, we computed batch removal and bio-conservation scores from the respective metrics by taking the mean of min-max scaled metric scores, which ensured equal discriminative power for all metrics and produced robust overall rankings (a previously used z-score scaling alternative^29^ gives highly correlated overall rankings: Spearman’s R>0.94 for all tasks). Overall scores combined batch removal and bio-conservation scores with a 40:60 weighting, which reflects the relative importance of optimizing each score: in simplified terms, optimal batch removal maps all cells to a single point, whereas optimal bio-conservation reflects each cell type being detectable in a single cluster.

Across RNA and ATAC integration tasks, we observed this apparent dichotomy between bio-conservation and batch effect removal, and each method strikes its own balance between the two. For instance, while BBKNN and Seurat v3 tended to remove batch variation, Conos and Scanorama prioritized bio-conservation. Interestingly, in non-graph-based methods a stronger tendency toward batch removal was mediated in parts by a more regularized learning of the implicit latent space representation of each batch. For example, Seurat v3 removed variation within cells from a single batch that otherwise showed substructure in unintegrated data (**Supplementary Note 3.5**). A highly regularized latent space is the likely cause of biological variation removal with increased batch effect removal. We hypothesize that improved latent space learning, or even projection to the “true” underlying data manifold, will enable more methods to remove strong batch effects between species, single-nucleus and single-cell data, or spatial locations.

Additionally, we found that preprocessing decisions strongly impact downstream integration quality. Indeed, scaling the input data typically shifted results toward better batch removal but worse bio-conservation, while HVG selection improved overall performance. Notably, only metrics that measured particular functions or pathways (i.e., cell cycle and trajectory conservation metrics) performed better with full gene sets. This suggests that biological functions are better captured in integrated data if the relevant gene sets are included in the integration. For all methods except Seurat v3 and Harmony, we identified an optimal preprocessing scheme. This finding affects the ease-of-use of Seurat v3 and Harmony, and it lowers their position in our overall ranking (because we ranked by a single, optimal preprocessing scheme).

We found that batch effects between ATAC datasets were a particular challenge for data integration. Thus, BBKNN, Harmony and scVI, with the highest batch removal scores, were a particular focus. Moreover, the overall poor integration performance resulted in our silhouette-based metrics to favour bio-conservation as compact, unintegrated clusters, showing a limitation of these metrics for poor integration performance. One particular batch effect issue in scATAC-seq is substantial data sparsity, which leads to a limited overlap of informative windows between datasets. Given the large number of total windows, this effect is likely to get stronger when integrating more cells and datasets. Using more features for integration is limited by the currently available methods, which often do not scale well to the number of features. Nevertheless, integration of ATAC and RNA has previously been achieved successfully by projecting onto gene features^12^. Although this feature choice represents a biased view of the chromatin landscape, using gene features would allow ATAC input data to resemble RNA inputs more closely. Future studies of ATAC integration, including ATAC-RNA integration using different feature sets, may uncover suitable integration approaches for this modality.

The deep learning (DL) methods, scVI and trVAE, performed better with increasing cell numbers and batch complexity. scVI performed particularly well when the task contained complex batch effects (e.g., microwell-seq, single-cell and single-nuclei, or scATAC-seq data) and sufficient numbers of cells were present to fit these effects. Similar performance has been reported for another DL method, scGen^30^ (not benchmarked here as it relies also on cell type information), on heart single-nucleus and single-cell data^31^. With more tunable parameters, these methods are more complex than other benchmarked methods and are more likely to require larger input data and hyperparameter optimization for optimal performance; however this also gives them the flexibility to fit complex batch effects. For scVI, a parameter set optimized for data integration was used (extracted from the respective tutorial). In contrast, trVAE was optimized for the more general and difficult task of perturbation modeling; this circumstance contributes to its poorer scalability without GPU hardware and thus prevented us from benchmarking it on the larger, complex tasks. While parameter optimization for an individual method would have biased our benchmarking result, the scVI platform contains the hyperopt tool^32^ for this purpose. Interestingly, scVI also performed well integrating data from full-length protocols provided in units of reads per kilobase million (RPKM) or transcripts per kilobase million (TPM), as well as with binary scATAC-seq data, although these data violate a central assumption of the method (i.e., negative binomially distributed input data). Comparatively, learning latent spaces with neural networks is still at an early stage of development. However, as the availability of data and accessibility of GPU hardware increases, we expect the performance of these methods to overtake that of their counterparts, as has occurred in the field of imaging^33,34^. Future benchmarks of DL integration methods using millions of cells and GPU hardware will better showcase the potential of these approaches.

An integration method should typically be chosen according to three criteria: usability, scalability, and expected performance. All ten methods in our study can be considered usable. For scRNA-seq data, the remaining considerations can be divided into five criteria: (1) the size of the dataset and hardware/software limitations, (2) compositional shifts in the data, (3) the type of output required, (4) the strength of the expected batch effect, and (5) the resolution of the integrated dataset, *i*.*e*., does the user require a general overview of the data or nuanced transcriptional differences. For exploratory data analysis, given no limitations or expectations of the batch effect size, we recommend the top-performing integration methods BBKNN, Scanorama, and scVI. For large datasets or setups with hardware limitations, we recommend ComBat or Harmony alongside BBKNN and scVI. However, the use of ComBat should be restricted to cases where compositional shifts between datasets are limited (e.g., mouse brain RNA or simulation 1; **Supplementary Data 3**).

Differing output formats can limit the potential downstream applications of integrated data. For example, BBKNN and Conos return integrated graphs, which can be used for downstream cell-level data analysis (such as clustering) but provide neither relative distances between cells nor corrected gene expression values (e.g., Conos outputs cannot be used to generate representative UMAPs without further processing). This limits certain methods because these latter outputs may be required for certain trajectory inference methods or for scoring functional gene programs. To obtain gene-expression outputs, we recommend trying Scanorama gene (but not Scanorama embedding) and MNN for complex batch setups, ComBat for simple batch setups, and Seurat v3 where distinct biological variation is expected.

Finally, the strength of the batch effect and the level of granularity required by the user in their data output must also be considered. For example, methods that remove strong batch effects (e.g., from species and single-nucleus vs single-cell data) also tend to remove nuanced biological signals such as rare cell types. Thus, if the aim is to find rare cell types and nuanced biological variation rather than remove strong batch effects, we recommend Scanorama. However, if a broad overview of the data in the presence of strong batch effects is required, we recommend Seurat v3 for smaller datasets. Given sufficient numbers of cells, scVI has shown that it is able to remove strong batch effects while only sacrificing minimal biological variation.

Where there are strong batch effects but the user is interested in nuanced biological information, other approaches may be needed. In general, it is worth considering whether removing a strong batch effect is desirable. In the present study, we have used set definitions of batch effect and biological variation, yet the distinction between the two is not always straightforward. Effects such as spatial location, species, or tissue could be either batch or biology depending on subjective opinion. Moreover, in certain cases, retaining batch effects in a dataset to preserve nuanced biological variation may be preferable. In such cases, statistical models can be used to directly analyse raw data while also accounting for linear batch effects. This type of modeling may also be appropriate across large, aggregated datasets^35^, for which sufficiently powerful data integration methods do not yet exist.

Our benchmarking study will help analysts to navigate the space of available integration methods and integrate their datasets more efficiently, and it will guide developers toward building more efficient methods. Based on the trends we have reported, users can select suitable preprocessing and integration methods for exploratory, integrated data analysis. To enable in-depth characterization of method performance on specific tasks, we have provided a reproducible Snakemake pipeline and the *scIB* python module to users so they can easily benchmark any preprocessing and integration method. Hence, we are supporting researchers to find the optimal integration method for their particular integration scenario. In addition, we expect that this work will become a reference for method developers, who can build upon the presented scenarios and metrics to assess the performance of their newly developed atlas-level data integration tasks.

## Methods

### Datasets and preprocessing

We benchmarked data integration methods on nine integration tasks: seven real data tasks and two simulation tasks. For the real data tasks we downloaded 23 published datasets (see **Supplementary Data 2** for per-batch overview of datasets). All scRNA-seq datasets were quality controlled and normalized in the same way according to published best practices^7^. Specifically, we used scran pooling normalization^36^ (version 1.10.2 unless otherwise specified) and log+1-transformation on count data. For data solely available in TPM or RPKM units we performed log+1-transformation without any further normalization. As the datasets typically contained different cell identity annotations; we mapped these annotations by matching annotation names, overlaps of data-driven marker gene sets, and manual clustering and annotation of cell identities per batch.

For the simulation tasks, data were simulated using the Splatter package^37^ to evaluate data integration methods in a controlled setting. All of our data processing scripts are publicly available as Jupyter notebooks and R scripts at www.github.com/theislab/scib.

#### Pancreas integration task

We used six publicly available human pancreas datasets. Specifically, we used a pre-annotated collection of four datasets from the Satija lab^38–41^ (retrieved from https://satijalab.org/seurat/v3.0/integration.html on 28/08/2019) with accession codes GSE81076, GSE85241, GSE86469 (GEO), and E-MTAB-5061 (ArrayExpress). The two additional human pancreas datasets were provided in a pre-annotated format by the Hemberg lab^42,43^ (https://hemberg-lab.github.io/scRNA.seq.datasets/human/pancreas/retrievedon28/08/2019); their GEO accession codes are GSE84133 and GSE81608. We normalized all datasets that contained count data with scran pooling^36^ in a joint normalization run. This excluded the dataset from Xin *et al*.^*42*^ which was provided in normalized units of RPKM. Finally, all datasets were log+1-transformed. In total, there were 16,382 cells in the pancreas integration task. Each dataset was treated as a batch, except for the inDrop dataset^43^, in which each donor was treated as a batch.

#### Immune cell integration tasks (human and mouse)

The immune cell task contained immune cells from eight datasets comprising human and mouse cells from bone marrow and peripheral blood. Bone marrow datasets were retrieved from Oetjen et al.^27^ (three human donors), Dahlin et al.^28^ (four mouse samples), and the Mouse Cell Atlas^44^ (MCA; three mouse samples). For peripheral blood data, mouse samples were downloaded from the MCA^44^ (six samples) and human samples were obtained from 10X Genomics^45^, Freytag *et al*.^*46*^, Sun *et al*.^*47*^ and Villani *et al*.^*26*^. Details on the retrieval location of datasets, the different protocols used, and ways in which samples were chosen for analysis can be found in **Supplementary Data 4**.

Quality control was performed separately for each sample. Sample-specific thresholds were chosen for the number of genes, the fraction of mitochondrial counts, and the number of UMI counts per cell. Datasets for which count data were available were individually normalized by scran pooling^36^. This excludes the data of Villani *et al*.^*26*^, which included only TPM values. All datasets were log+1-transformed in Scanpy (version 1.4.4 commit bd5f862)^48^.

To create a consistent set of cell identity annotations across datasets, we harmonized the existing labels and annotated cells from datasets in which no labels were available. First, the label sets suggested by the MCA and Oetjen *et al*.^*27*^, were harmonized by string matching. In the second step, we collected a number of cell identity markers from the literature (**Supplementary Data 5**) and tested them, first on the pre-annotated samples, and then on the remaining samples. This procedure allowed us to refine the annotation by adding a second layer of cell labels. Where necessary, we performed sub-clustering to improve the annotations. Finally, if the annotations could not be mapped due to coarse labeling, we removed cell populations.

We created two integration tasks from the immune cell data: one containing only human samples, and one containing both human and mouse samples. The human task included cross-tissue integration of immune cells from many donors; the combined task added the complexity of cross-species integration. To integrate human and mouse data into a single data object, we mapped mouse genes (MGI symbol) to their human counterparts (HGNC symbol) using the R package biomaRt (version 2.38.0)^49^. We retained only those genes that were mapped in all batches: 8,135 genes in total. The human integration task contained 33,506 cells, whereas the combined task contained 97,952 cells. Sample IDs were used as batches for data integration.

To test the conservation of trajectories following data integration, we considered the process of erythropoiesis in the human and mouse bone marrow datasets. Specifically, we extracted HSPCs, MPs, EPs, and mature erythrocytes for each batch. We generated a trajectory for each sample using Scanpy’s diffusion maps^50^ and diffusion pseudotime^51^ functions. The root cell for pseudotime analysis was selected from the HSPCs cluster upon evaluation of the diffusion components. Specifically, we selected the cell that was assigned the maximum or minimum value of the first three diffusion components as the root cell.

#### Lung atlas integration task

Single-cell expression data for the lung integration task was retrieved from the work of Vieira Braga et al.^52^, who created a lung atlas that includes samples from three labs that were generated using Drop-seq and 10X Chromium. The Drop-seq data was available from GEO under accession code GSE130148, while the 10X data was obtained directly from the authors in a SoupX-corrected count matrix. We used three healthy datasets from Vieira Braga et al.^52^: the 10X and Drop-seq transplant datasets, along with 10X lung biopsy data. Nasal brush and lung brush samples were not included in the integration task, as suggested by the original authors, due to the cell identity populations being distinct from the other three datasets. However, we did include lung biopsy data, which comes from a distinct spatial location (the airways) relative to the location of transplant samples (the parenchyma). Following quality control filtering, the data contained 16 donors, with one sample per donor, and 32,472 cells.

Data were normalized by scran pooling^36^, which was applied to individual datasets. As the 10X datasets and the Drop-seq dataset contained different cell annotations, the annotations were harmonized using fuzzy string matching and overlaps of marker genes determined by a t-test performed in Scanpy^48^ (version 1.4.5 commit d69832a). Where annotations could not be mapped due to coarse labeling or where cell populations corresponded to filtered-out datasets, the cell populations were removed (annotations: *Mesothelium, Transformed epithelium, Ciliated (Nasal), Goblet 1 (Nasal), Goblet 2 (Nasal)*, and *Smooth Muscle Cells*). Donor IDs were used as batches for data integration.

#### Mouse brain integration task (RNA)

The mouse brain RNA task consisted of four publicly available scRNA-seq and snRNA-seq mouse brain studies^53–56^, in which additional information on cerebral regions was provided. We obtained the raw count matrix for the snRNA-seq dataset (SPLiT-seq protocol) of Rosenberg *et al*.^*53*^ (GEO accession ID: GSE110823), the annotated count matrix (10X Genomics protocol) from Zeisel *et al*.^*54*^ (http://mousebrain.org; file name L5_all.loom, downloaded on 09/09/2019), and the count matrices per cell type (Drop-seq protocol) from Saunders *et al*.^*56*^ (http://dropviz.org/; *DGE by Region* section, downloaded on 30/08/2019). FACS-sorted mouse brain tissue data (10X Genomics protocol, myeloid and non-myeloid cells, including the annotation file “annotations_FACS.csv”) from Tabula Muris^55^ were obtained from <u>figshare</u> (retrieved 14/02/2019).

We harmonized cluster labels via fuzzy string matching, attempting to preserve the original annotation wherever possible. Specifically, we annotated 10 major cell types (neurons, astrocytes, oligodendrocytes, oligodendrocyte precursor cells, endothelial cells, brain pericytes, ependymal cells, olfactory ensheathing cells, macrophages, and microglia).

From Saunders *et al*.^*56*^, we used the additional annotation data table to obtain 585 reported cell types (annotation.BrainCellAtlas_Saunders_version_2018.04.01.txt, retrieved from http://dropviz.org/ on 30/08/2019). Among these cell types, some were annotated as *endothelial tip, endothelial stalk* and *mural*, which had no correspondence in other datasets. Thus, we re-annotated these cell types as follows: Louvain clustering (default resolution parameter 1.0) was applied to cluster cells; gene expression profiling was conducted using the *rank_genes_groups* function in Scanpy (t-test); and microglia (*C1qa*), oligodendrocytes (*Plp1*), astrocytes (*Gfap* and *Clu*), and endothelial cells (*Flt1*) were assigned using marker gene expression.

In addition, we harmonized brain region information where possible. In total, we annotated 15 different brain regions (the amygdala, hippocampus, thalamus, hypothalamus, cortex, olfactory bulb, striatum, cerebellum, midbrain, medulla, substantia nigra, entopeduncular nucleus, globus pallidus and nucleus basalis, pons and spinal cord). It must be noted that Rosenberg *et al*.*53* inferred brain regions; thus, 66,648 cells in this dataset were not assigned to a brain region (marked as *Unknown* in the data).

Finally, we applied scran normalization^36^ separately to each dataset and log+1-transformed the count matrices. In total, this mouse brain integration task contained 978,734 cells. Datasets were treated as batches for data integration.

#### Mouse brain integration task (ATAC)

The mouse brain ATAC task consists of three single-cell ATAC-seq datasets. We obtained count matrices from Fang *et al*.^57^ (six samples obtained using a single nucleus ATAC-seq protocol; retrieved from http://renlab.sdsc.edu/r3fang/share/github/Mouse_Brain_MOp) and Cusanovich *et al*.^*58*^ (four samples obtained using a combinatorial indexing ATAC-seq protocol; GEO accession number GSE111586)^58^ and we retrieved BAM files from 10X Genomics (one sample retrieved from https://support.10xgenomics.com/single-cell-atac/datasets by Cell Ranger ATAC 1.2.0 on 05/12/2019). We used *pyliftover* (https://github.com/konstantint/pyliftover) and liftover chains from UCSC to convert the Cusanovich *et al*.^*58*^ data from the mm9 to the mm10 reference genome (Genome Reference Consortium Mouse Build 38, GRCm38). EpiScanpy^59^ version 0.1.10 was used to conduct preprocessing steps. For the 10X Genomics dataset, we first built binary count matrices, and then removed features covering fewer than 10 cells, before removing cells with fewer than 5000 measured features. For the Cusanovich *et al*.^*58*^ data, the features with fewer than 30 cells covered and the cells with fewer than 3000 measured features were removed. For the Fang *et al*.^*57*^ dataset, the features covered in fewer than 10 cells and the cells with fewer than 500 measured features were removed. Subsequently, we selected the 150,000 most variable features across the cells in each dataset. The available annotations of Cusanovich *et al*.^*58*^ and a list of differentially opened regions corresponding to marker genes from Danese *et al*.^59^ were used to annotate cell types.

Two ATAC integration tasks were completed: the small ATAC task consisted of selected samples from three datasets [all four samples from Cusanovich *et al*.^58^, one sample from 10X Genomics, and one sample from Fang *et al*.^*57*^ (CEMBA180305_2B)]; the large ATAC task consisted of all samples from the three datasets. After the open chromatin matrices of the two tasks were constructed, we filtered out the cells with fewer than 500 measured features. Finally, we performed a library size correction and a log+1-transformation. Ultimately, the small ATAC task contained 57,070 features and 25,960 cells from six samples; the large ATAC task consisted of 57,447 features and 67,612 cells from 11 samples.

#### Simulations

We generated synthetic datasets using an extended version of the *Splat* simulation method available in the Splatter package^37^. The standard *Splat* model produces batches with equal cell group proportions and expected library sizes. In order to modify these factors, we first generated a larger dataset in which each batch had an equal number of cells and each group was present in equal proportions. We then used a downsampling procedure to remove cells from each batch until the desired cell group proportions were obtained. The desired difference in the number of counts per cell between batches was achieved using the *downsampleMatrix* function from the DropletUtils package^60,61^; counts were downsampled in the resulting counts matrix. Basic quality control, involving the removal of cells >2 median absolute deviations below the median of counts per cell or number of expressed genes per cell within each batch, was then performed on the simulated data using the *quickPerCellQC* function in the Scater package^62^. Genes expressed in <1% of cells in the whole simulation were also removed. This resulted in an integration task consisting of 12,097 cells and six batches.

To create the nested batch effect simulation scenario, we added a step between adjusting cell group proportions and downsampling counts in order to create a sub-batch structure. For each sub-batch, we used the *Splat* model to simulate a second count matrix with the same number of cells as the sub-batch but a lower expected library size and no cell group structure. We then added this noise matrix to the counts for cells in that sub-batch. Quality control for the nested batch scenario was performed at the sub-batch level, and sub-batches were used as batch IDs for integration. The nested batch integration task consisted of 19,318 cells and 16 nested sub-batches (four sets of four sub-batches).

### Integration methods

#### ComBat

ComBat^19^ is a batch correction method developed for bulk gene expression microarray data. It uses a linear mixed effect model that fits the batch effect’s contribution both to the mean expression and the variance in expression. We ran ComBat as it is implemented in Scanpy (version 1.4.5 commit d69832a) via the *combat* function. ComBat returns a corrected gene expression or open chromatin matrix.

#### Matching mutual nearest neighbors (MNN)

MNN first detects mutual nearest neighbors in two datasets (or batches) and then infers a projection of the second dataset into the first dataset, which serves as a reference hyperplane^11^. This integrated dataset serves as a new reference to iteratively integrate more datasets. We ran MNN using the *mnn_correct* function from mnnpy (https://github.com/chriscainx/mnnpy version 0.1.9.5). The default parameters were used, including an additional cosine normalization of the input matrix. MNN returns a corrected gene expression or open chromatin matrix.

#### scVI

The scVI model combines a variational autoencoder (a neural network) with a hierarchical Bayesian model^13^. The negative binomial distribution is used to describe the gene expression of each cell, conditioned on the batch variable and unobserved factors such as differences in sensitivity between measurements. Thus, scVI takes into account fixed and random effects in the data. The output of scVI is a low-dimensional representation in latent space (an embedding). Notably, scVI expects a raw count matrix as input; this was not always available in our integration tasks. We ran scVI (version 0.5.0) using the parameterizations from the *scanpy_pbmc3k* (https://scvi.readthedocs.io/en/stable/tutorials/scanpy.html) and the *harmonization* (https://scvi.readthedocs.io/en/stable/tutorials/harmonization.html) tutorial notebooks. This parameterization includes a model with negative binomial reconstruction loss, a 30-dimensional latent space, 128 nodes in the hidden layer, and n_layers = 2. After consulting with the authors, the model was trained for 400*(20,000/*N*) epochs where *N* is the size of the dataset, while implementing a maximum of 400 epochs for small datasets. scVI returns a joint embedding of cells from all batches.

#### Scanorama

The Scanorama algorithm is based on the concept of panoramic stitching. It finds similar cells across datasets using a k-nearest neighbor search, and then reduces the connections to a set of mutual nearest neighbors^14^. Subsequently, all data points are embedded in a joint hyperplane. In the absence of a clear tutorial, we ran Scanorama (version 1.4) via the *correct_scanpy* function with the option *return_dimred=True* to obtain a joint embedding as well as a corrected expression matrix.

#### Batch-balanced k-nearest neighbors (BBKNN)

BBKNN^15^ first computes a k-nearest neighbor graph within each batch. It then computes the k-nearest neighbors of all cells to all other batches. The resulting graph contains a number of irrelevant connections across cell types; therefore, BBKNN computes a connectivity score for each pair of cells similar to the UMAP algorithm^63^. The symmetrized connectivity score represents the connection of each pair of cells. Thus, BBKNN ultimately returns a weighted neighborhood graph. Notably, BBKNN requires at least some cell types to be shared across batches. We ran BBKNN (version 1.3.5) using the *bbknn* function with mainly default parametrization. Following previous comparison runs with BBKNN from the Harmony paper^18^, we used *k* = 15 as the number of neighbors within each batch and *t* = 20 as trim parameter for data scenarios with <100,000 cells. This parametrization prevents the global network from becoming too large. For integration tasks with ≥100,000 cells, we used *k* = 30 and *t* = 30 instead.

#### Clustering on network of samples (Conos)

The Conos model constructs a joint graph of all batches in a two step process^17^. First, Conos creates pairwise connections across batches to initialize connections between identical cell types. Specifically, common principal component analysis or joint non-negative matrix factorization is used to create a joint space for cell-cell similarity computation. The cell-cell similarity scores serve as weights of the connections across datasets. Second, the number of inter-batch connections is reduced by a mutual nearest neighbor approach, and connections within a batch are down-weighted by 0.1 to account for the inherently higher cell-cell similarities of cells from the same cell type within a dataset. Ultimately, Conos returns a corrected neighborhood graph. We ran Conos (version 1.2.1) as described in the online tutorial *scanpy_integration* (https://github.com/hms-dbmi/conos/blob/master/vignettes/scanpy_integration.md). This tutorial includes HVG selection, scaling, and PCA runs per batch in Seurat. Given that Conos objects require these slots filled in order to run, we regarded the aforementioned steps as part of the Conos method. Any preprocessing combinations that we benchmark were conducted prior to the HVG selection and scaling performed within the function.

#### Seurat v3

The Seurat v3 algorithm uses canonical correlation analysis to construct a shared subspace of two batches^12^. The algorithm then identifies mutual nearest neighbors across the two datasets, which are called “anchor points”. A projection vector is then inferred from the anchor points to integrate the two datasets in a common reference hyperplane. The same projection vectors serve to integrate new cell populations without mutual neighbors. Integrating multiple datasets involves pairwise computation of anchor points followed by hierarchical clustering based on the distance between the datasets. The resulting tree defines the integration order to iteratively construct the common corrected data matrix. We ran Seurat v3 (version 3.1.1) according to Seurat’s integration tutorial (https://satijalab.org/seurat/v3.0/integration.html). After discussion with the method’s authors, modifications were made to the standard parametrization to allow the passage of HVGs from Scanpy directly to the method. Seurat v3 returns a corrected gene expression or open chromatin matrix.

#### Harmony

The Harmony^18^ algorithm initializes all datasets in PCA space along with the batch variable and alternately iterates over two complementary concepts until convergence. First, it employs maximum diversity clustering, which penalizes overcorrection and pushes clusters with the same cells apart. Second, batch effects are accounted for by a linear mixture model. Thus, Harmony returns a corrected embedding. We ran Harmony (version 1.0) according to its tutorial (http://htmlpreview.github.io/?https://github.com/immunogenomics/harmony/blob/master/docs/SeuratV3.html). As Harmony requires scaling and PCA to be run within Seurat, we regarded these steps as part of the Harmony method. Thus, any scaling or HVG selection we benchmarked occurred upstream of the scaling performed by Harmony as part of its standard workflow.

#### LIGER

LIGER performs integrative non-negative matrix factorization to integrate diverse batches. This approach consists of factorizing each batch expression matrix into a dataset-specific factor matrix and a shared factor matrix. The shared factor matrix is used as a joint embedding for cells across batches. We ran LIGER (version 0.4.2) with the default parameters (*k* = 20 and lambda = 5), as suggested in the online tutorial (https://macoskolab.github.io/liger/walkthrough_pbmc.html). This LIGER tutorial includes scaling without zero-centering and HVG selection. The custom scaling function is used as LIGER cannot accept negative input values; thus, testing our preprocessing decisions for scaling would go against the best practices for the tool. As LIGER does not give the user the flexibility to easily run alternative data scaling, we considered the LIGER scaling function to be part of the method; consequently, we only assessed the effect of HVG selection with this method.

#### Transformer Variational Autoencoder (trVAE)

The trVAE model is a conditional variational autoencoder developed for out-of-sample prediction specifically on perturbations^20^. Specifically, it uses the maximum mean discrepancy measure for distribution matching in the first decoding layer. Thus, trVAE returns both an embedding and a corrected data matrix. We ran trVAE (version 0.0.1) according to the *trVAE_Haber* example notebook (https://nbviewer.jupyter.org/github/theislab/trVAE/blob/master/examples/trVAE_Haber.ipynb). As trVAE cannot take negative values as input, we omitted scaling when testing our preprocessing decisions for this method. trVAE returns a joint embedding of cells from all batches. It can also output a corrected gene expression or open chromatin matrix, but this output was not tested here.

### Metrics

We grouped the metrics into two broad categories: (1) removal of batch effects and (2) conservation of biological variance. The latter category is further divided into conservation of variance from cell identity labels, and conservation of variance beyond cell identity labels. Scores from the first category include principal component (PC) regression (batch), ASW (batch), graph connectivity, graph iLISI, and kBET. In the second category, label conservation metrics include NMI, ARI, ASW (cell type), graph cLISI, isolated label F1 and isolated label silhouette; label-free conservation metrics include cell cycle (CC) conservation, HVG conservation, and trajectory conservation.

The metrics were run on different output types (**Supplementary Table 2**). For example, metrics that run on k-nearest neighbor (kNN) graphs can be run on all output types after preprocessing. Similarly, metrics that run on joint embeddings can also be run on corrected feature outputs. Preprocessing was performed in Scanpy (version 1.4.5 commit d69832a). kNN graphs were computed using the *neighbors* function where *k* = 15 unless otherwise specified. Where a joint embedding was available, this graph was computed using Euclidean distances on this embedding, whereas distances were computed on the top 50 PCs where a corrected feature matrix was output.

#### Normalised mutual information (NMI)

NMI compares the overlap of two clusterings. We used NMI to compare the cell type labels with Louvain clusters computed on the integrated dataset. The overlap was scaled using the mean of the entropy terms for cell type and cluster labels. Thus, NMI scores of 0 or 1 correspond to uncorrelated clustering or a perfect match, respectively. We performed optimized Louvain clustering for this metric to obtain the best match between clusters and labels. Louvain clustering was performed at a resolution range of 0.1 to 2 in steps of 0.1, and the clustering output with the highest NMI with the label set was used. We used the scikit-learn^23^ (version 0.22.1) implementation of NMI.

#### Adjusted Rand Index (ARI)

The Rand index compares the overlap of two clusterings; it considers both correct clustering overlaps while also counting correct disagreements between two clusterings^64^. Similar to NMI, we compared the cell type labels with the NMI-optimized Louvain clustering computed on the integrated dataset. The adjustment of the Rand index corrects for randomly correct labels. An ARI of 0 or 1 corresponds to random labelling or a perfect match, respectively. We also used the scikit-learn^23^ (version 0.22.1) implementation of the ARI.

#### Average silhouette width (ASW)

The silhouette width measures the relationship between the within-cluster distances of a cell and the between-cluster distances of that cell to the closest cluster^65^. Averaging over all silhouette widths yields the ASW, which ranges between −1 and 1. Originally, ASW was used to determine the separation of clusters where 1 represents dense and well-separated clusters. Furthermore, an ASW of 0 or −1 corresponds to overlapping clusters (caused by equal between- and within-cluster variability) or strong misclassification, respectively. We used the classical definition of ASW to determine the silhouette of the cell labels (cell type ASW). For this bio-conservation score, ASW was linearly scaled to a value between 0 and 1 using the equation *cell type ASW* = (*ASW* + 1)/2, where larger values indicate denser clusters. Furthermore, we also used ASW to describe the mixing of batches within cell clusters (batch ASW^9^). In this usage, an ASW of 0 indicates that batches are well-mixed, which is preferable. To obtain the batch ASW, we scaled ASW scores via the equation *batch ASW* = 1 − *abs*(*ASW*); thus, batch ASWs of 1 and 0 represent ideally mixed cases and strongly separated clusters, respectively.

We computed the ASW based on the PCA embedding of corrected expression data or on the integrated embedding output using the scikit-learn^23^ (version 0.22.1) implementation.

#### Principal component regression (PC regression)

PC regression, derived from PCA, has previously been used to quantify batch removal^9^. Briefly, the R^2^ was calculated from a linear regression of the covariate of interest (e.g., the batch variable *B*) onto each PC. The variance contribution of the batch effect per PC was then calculated as the product of the variance explained by the *i*^*th*^ PC and the corresponding *R*^2^(*PC*_*i*_ |*B*). The sum across all variance contributions by the batch effects in all PCs gives the total variance explained by the batch variable as follows:

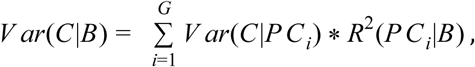

where *V ar*(*C*|*PC*_*i*_) is the variance of the data matrix *C* explained by the *i*^*th*^ PC.

#### Graph connectivity

The graph connectivity metric assesses whether the kNN graph representation, *G*, of the integrated data directly connects all cells with the same cell identity label. For each cell identity label *c*, we created the subset kNN graph *G(N*_*c*_;*E*_*c*_) to contain only cells from a given label. Using these subset kNN graphs, we computed the graph connectivity score using the equation:

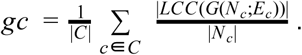

Here, *C* represents the set of cell identity labels, |*LCC()*| is the number of nodes in the largest connected component of the graph, and |*N*_*c*_| is the number of nodes with cell identity *c*. The resultant score has a range of (0;1], where 1 indicates that all cells with the same cell identity are connected in the integrated kNN graph, and the lowest possible score indicates a graph where no cell is connected. As this score is computed on the kNN graph, it can be used to evaluate all integration outputs.

#### K-nearest neighbor batch effect test (kBET)

The kBET algorithm (version 0.99.6, release 4c9dafa) determines whether the label composition of a k-nearest neighborhood of a cell is similar to the expected (global) label composition^9^. The test is repeated for a random subset of cells, and the results are summarized as a rejection rate over all tested neighborhoods. Thus, kBET works on a k-nearest neighbor (kNN) graph.

We computed kNN graphs where *k* = 50 for joint embeddings and corrected feature outputs via the Scanpy preprocessing steps (previously described). To test for technical effects and to account for cell type frequency shifts across datasets, we applied kBET separately on the batch variable for each cell identity label. Using the kBET defaults, a *k* equal to the median of the number of cells per batch within each label was used for this computation. Additionally, we set the minimum and maximum thresholds of *k* to 10 and 100, respectively. As kNN graphs that have been subset by cell identity labels may no longer be connected, we computed kBET per connected component. If >25% of cells were assigned to connected components too small for kBET computation (smaller than *k*\*3), we assigned a kBET score of 1 to denote poor batch removal. Subsequently, kBET scores for each label were averaged and subtracted from 1 to give a final kBET score.

We noted that k-nearest neighborhood sizes can differ between graph-based integration methods (e.g., Conos and BBKNN) and methods in which the kNN graph is computed on an integrated embedding. This difference can affect the test outcome because of differences in statistical power across neighborhoods. Thus, we implemented a diffusion-based correction to obtain the same number of nearest neighbors for each cell irrespective of integration output type (**Supplementary Note 1**). This extension of kBET allowed us to compare integration results on kNN graphs irrespective of integration output format.

#### Graph local inverse Simpson’s Index (graph LISI)

The LISI, a diversity score, was proposed to assess both batch mixing (iLISI) and cell type separation (cLISI)^18^. LISI scores are computed from neighborhood lists per node from integrated kNN graphs. Specifically, the inverse Simpson’s index is used to determine the number of cells that can be drawn from a neighbor list before one batch is observed twice. Thus, LISI scores range from 1 to N, where N is the total number of batches in the dataset.

Typically, neighborhood lists to compute LISI scores are extracted from weighted kNN graphs with *k* = 90 nearest neighbors at a fixed perplexity of 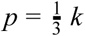. These nearest neighbor graphs are constructed using Euclidean distances on PCA or other embeddings. In contrast, integrated graphs that are output by methods such as Conos or BBKNN typically contain far fewer than *k* = 90 neighbors. Running LISI metrics with differing numbers of nearest neighbors per node showed a bias of LISI scores toward graph-based integration outputs (data not shown). Thus, the original LISI score is not applicable to graph-based outputs.

To extend LISI graph-based integration outputs, we developed *graph LISI*, which uses the integrated graph structure as an embedded space for distance calculation. The calculated graph distances are then used to determine a consistent number of nearest neighbors per node. We used the shortest path lengths computed via Dijkstra’s algorithm^66^ as a graph-based distance metric (see **Supplementary Note 2** for details). Our graph LISI extension produces consistent metric values with the standard LISI implementation for non-graph-based integration outputs (**Supplementary Fig. 24**).

As LISI scores range from 1 to N, indicating perfect separation and perfect mixing respectively, we rescaled them to the range 0 to 1. For iLISI and cLISI this involved a three-step process. First, we used scalings for cLISI and iLISI as follows: *cLISI*: *f* (*x*) = 2 − *x*, where a low value corresponds to low cell type separation; *iLISI* : *g*(*x*) = *x* − 1, where a low value corresponds to low batch integration. Second, we computed the median across neighborhoods per method: *cLISI* = *median f* (*x*), *x* ∈ *X*; *iLISI* = *median g*(*x*), *x* ∈ *X*. Finally, we rescaled the LISI scores by the minimum and maximum observed median scores across tasks.

#### Isolated label scores

We developed two isolated label scores to evaluate how well the data integration methods dealt with cell identity labels shared by few batches. Specifically, we identified isolated cell labels as the labels present in the least number of batches in the integration task. The score evaluates how well these isolated labels separate from other cell identities.

We implemented two versions of the isolated label metric: the isolated label F1 and isolated label ASW. The F1 score metric first determines the cluster with the largest number of an isolated label; the F1-score of the cells of the isolated label is then computed against the cells within the cluster. Specifically, the F1 score is a weighted mean of precision and recall given by the equation:

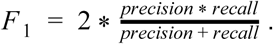

It returns a value between 0 and 1, where 1 shows that all of the isolated label cells and no others are captured in the cluster. The isolated label ASW score computes the ASW on the PCA embedding subset to the isolated labels (see ASW metric above). This score is scaled to between 0 and 1 as described for the ASW score. For both functions, in cases of multiple isolated labels, the mean score of all isolated labels is returned as the final score.

#### HVG conservation

The HVG conservation score is a proxy for the preservation of the biological signal. If the data integration method returned a corrected data matrix, we computed the number of HVGs before and after correction for each batch via Scanpy’s *highly_variable_genes* function (using flavor = “cell ranger”). If available, we computed 500 HVGs per batch. If fewer than 500 genes were present in the integrated object for a batch, the number of HVGs was set to half the total genes in that batch. The overlap coefficient is as follows:

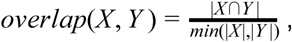

where *X* and *Y* denote the fraction of preserved informative genes. The overall HVG score is the mean of the per-batch HVG overlap coefficients.

#### Cell cycle conservation

The cell cycle conservation score evaluates how well the cell cycle effect can be scored before and after integration. We computed cell cycle scores using Scanpy’s *score_cell_cycle* function with a reference gene set from Tirosh *et al*.^*67*^ for the respective cell cycle phases. We used the same set of cell cycle genes for mouse and human data (using capitalization to convert between the gene symbols). We then computed the variance contribution of the resulting S and G2/M phase scores using PC regression (see Principal Component regression), which was performed for each batch separately. The differences in variance before, *V ar*_*before*_, and after, *V ar*_*after*_, integration were aggregated into a final score between 0 and 1, using the equation:

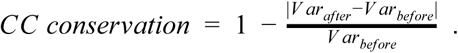

In this equation values close to 0 indicate lower conservation and 1 indicates complete conservation of the variance explained by cell cycle. In other words, the variance remains unchanged within each batch for complete conservation, while any deviation from the pre-integration variance contribution reduces the score.

#### Trajectory conservation

The trajectory conservation score is a proxy for the conservation of the biological signal. We compared trajectories computed after integration for certain clusters that had been manually selected during the data preprocessing step. Trajectories were computed using diffusion pseudotime implemented in Scanpy (*sc*.*tl*.*dpt*). We assumed that trajectories found in the unintegrated data for each batch gave the most accurate biological signal. Therefore, the starting cell of the trajectory, post-integration, was defined by selecting the most extreme cell from the cell type cluster that contained the starting cells of the pre-integration diffusion pseudotime, which was based on the first three diffusion components (see the immune cell task description for more details). Only cells from the largest connected component of the neighborhood graph were considered.

We computed Spearman’s rank correlation coefficient, *s*, between the pseudotime values before and after integration. The final score was scaled to a value between 0 and 1 using the equation *trajectory conservation* = (*s* + 1)/2. Values of 1 or 0 correspond to the same order of cells on the trajectory before and after integration or the reverse order, respectively.

### Benchmarking setup

All integration runs were performed using our Snakemake pipeline. Methods were tested with scaled and unscaled data as input, using the full feature (gene/open chromatin window) set or only HVGs. Where HVGs were used, the top 2000 were selected using a custom method, which selected HVGs in a manner unaffected by batch variance. Specifically, we initially built the *hvg_batch* function on top of the *highly_variable_genes* function from Scanpy. Using the standard function from Scanpy, we obtained the top 2000 HVGs per batch with the *cell_ranger* flavor. The list of HVGs was ranked first by the number of batches in which the genes were highly variable and second by the mean dispersion parameter across batches; the top 2000 were then selected. This *hvg_batch* function is freely available as part of the *scIB* module. Scaled data have zero mean and unit variance per gene; this was performed by calculating z-scores of the expression data using Scanpy’s *scale* function applied separately to each batch (*scale_batch* function in *scIB*). HVG selection and scaling were not applied in the ATAC tasks, as these are not typical steps in an ATAC workflow.

Data integration runs were performed with 12 cores and 24 threads available to each method; 16 GB of memory per core and 131 GB of shared swap memory were available. Thus, up to 323 GB of memory was available for each run. The runtime limit was set to four days (96 hours). Some methods ran out of time or memory and were assigned NA values for the respective integration task. The integration methods were run in separate conda environments for R and Python methods to ensure no clashes in dependencies. Details on how to set up these environments can be found on the *scIB* GitHub repository (www.github.com/theislab/scib). We converted between R and Python data formats using anndata2ri (www.github.com/theislab/anndata2ri) and conversion functions in LIGER and Seurat.

Metrics were run on the integrated and unintegrated AnnData^48^ objects. We selected the metrics for evaluating performance based on the type of output data (**Supplementary Table 2**). For example, metrics based on corrected embeddings (Silhouette scores, PC regression, and cell cycle conservation) were not run where only a corrected graph was output. We calculated an overall score per integration run by taking the weighted mean of the batch removal and bio-conservation scores (weights of 0.4 for batch removal and 0.6 for bio-conservation). In turn, these scores were computed by taking the weighted mean of all metrics that were computed in this category. Weighting was performed by min-max scaling of the score across all integration runs within each task, so that each metric was equally discriminative between all integration runs. Notably, scaling via z-scores (previously used for trajectory benchmarking^29^) instead of min-max gives similar overall rankings (Spearman’s R >0.94 for all tasks). Using this method, we were able to compute comparable overall performance scores even when different numbers of metrics were computed per run.

### Usability assessment

We assessed the usability of integration methods, via an adapted objective scoring system. A set of nine categories were defined (adapted from Saelens *et al*.^*29*^) to comprehensively evaluate the user-friendliness of each method (**Supplementary Fig. 21** and **Supplementary Data 6**). The first five categories (open source, version control, unit testing, tutorial and function documentation) assessed the quality of the code, its availability, the presence of a tutorial to guide users through one or more examples, and (ideally) usage in a non-native language (i.e., from Python to R or vice versa). The other four categories (peer-review, evaluation of accuracy, evaluation of robustness, and benchmarking) assessed whether the method was published in a peer-reviewed journal, how the paper evaluated the accuracy and robustness of the method, and the inclusion of benchmarking with other published algorithms in the paper. The mean value for each category was calculated to obtain a partial score that was then averaged over all categories; this led to one final usability score. Each category was considered to be equally important. **Supplementary Data 6** reports scores and references collected, to the best of our knowledge, for each usability term considered. In particular, we sought information from multiple sources, such as GitHub repositories, Bioconductor vignettes, Readthedocs documentation, original manuscripts, and supplementary material.

### Scalability assessment

The scalability of all data integration tools was assessed according to CPU time and peak memory use. For each run of the Snakemake pipeline, we used the Snakemake benchmarking function to measure time and peak memory use (max RSS). To score time and memory usage, we used a linear regression model to fit time and memory vs. the number of cells on a log-scale separately for each method and each preprocessing combination (completed with *curve_fit* from *scipy*.*optimize*, scipy version 1.3.0). The fit results are shown in **Supplementary Fig. 19**. Each fit had a slope and an intercept calculated as follows:

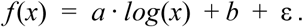

These values were used to compute each area under the curve (AUC) where *A* = 10^4^ and *B* = 10^6^, which corresponded to the approximate range of data task sizes in our study. To derive a scalability score from these areas, we scaled all AUCs by the area of the rectangle that covered all curves. Specifically, we chose the width as the difference of the log-scaled bounds and the height *C* as 10^8^ s (≈ 3 years) and 10^9^ MB (≈1 PB), respectively:

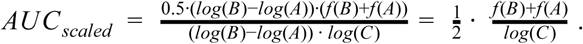

Methods that scale well have a low AUC and, consequently, a low scaled AUC. To obtain a consistent scoring scheme, we inverted the scaled AUCs:

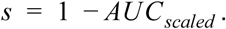

Finally, we reported the scalability scores for CPU time and peak memory use per method and preprocessing combination.

### Visualization

Inspired by the code of Saelens *et al*.^*29*^, we implemented two plotting functions in R. The first visualization displays each integration task separately and shows the complete list of tested integration runs ranked by the overall performance score. Individual and aggregated scores are represented by circles and bars, respectively. The color scheme indicates the overall ranking of each method.

The second visualization provides an overall view of the best performing flavors of each integration method. To obtain this, we first calculated the overall score over metrics for each method in each task (considering only real scRNA-seq data integration tasks). Subsequently, we ranked the methods in each real scRNA-seq task and computed an average rank over scenarios. Importantly, methods that could not be run for a particular task were assigned the same rank as unintegrated data on this task. Finally, we chose the best performing combination of *features (HVG or full features)* and *scaling* flavors for each integration method, and then ranked these from best-to worst-performing. Moreover, we displayed an average usability score, two scalability scores related to time and memory consumption, and the overall scores obtained in the two simulation tasks (although these scores were not used for the ranking). Again bar lengths represented scores and the color scheme indicated the ranking.

## Data and Code availability

The data used in this paper is publicly available and retrievable as described in the data section. Notebooks and R scripts used to preprocess the data, and all of the preprocessing and integration functions, evaluation metrics, and workflows used, are available at www.github.com/theislab/scib.

## Author Contributions

MDL, MB, KC, MI, MCT and FJT wrote the paper. MDL designed the study together with FJT. MDL, AD, MB, and MI prepared the data and LZ prepared the simulations. MI, LZ, DS, MB, MDL, and KC produced the figures. DS, MM, MDL, MB, and KC wrote the code. MB and MDL extended the kBET metric and developed graph LISI, and MM, DS, and MDL developed the novel label-free metrics. KC, MCT, MDL, MB, and MI performed the analysis. MDL, MD, MCT, and FJT supervised the work. All authors reviewed the final paper.

## Acknowledgements

We would like to acknowledge Martijn Nawijn, Herbert Schiller, and Lukas Simon for provision of data and expertise in mapping of lung cell annotations. Philipp Angerer was of great help for timely bug fixes in package dependencies. Further, we would like to thank Rahul Satija and Dana Pe’er for conversations on the evaluation of integration methods, and Mohammad Lotfollahi, Viktor Petukhov, Tim Stuart, Andrew Butler, Romain Lopez, and Adam Gayoso for help on getting their respective methods running in our environment and for discussions on fair evaluation of these methods. Johann Hawe was instrumental in helping us to get our Snakemake pipeline working as we envisioned, and we would like to thank Volker Bergen for help with diffusion computations on connectivity matrices. Special thanks also to Thomas Neumann, who was the crucial driver for being able to greatly speed up our nearest neighbour finding algorithm in C++ to make graph LISI scalable to millions of cells. Finally, we would like to thank all the members of the Theis lab for feedback and discussions.

This work was supported by the ‘ExNet-0041-Phase2-3 („SyNergy-HMGU”)’ and the Incubator grant # ZT-I-0007 sparse2big, both through the Initiative and Network Fund of the Helmholtz Association and by the Chan Zuckerberg Initiative DAF (advised fund of Silicon Valley Community Foundation, 182835), the European Union’s Horizon 2020 research and innovation programme under grant agreement No 874656 and by the Chan Zuckerberg foundation (grant #2019-002438, Human Lung Cell Atlas 1.0).

## Notes

### Competing Interest Statement

F.J.T. reports receiving consulting fees from Roche Diagnostics GmbH and Cellarity Inc., and ownership interest in Cellarity, Inc. and Dermagnostix

### Summary of Updates

Added 1 citation and amended spelling of author names with accents.

