## Supplementary for "Benchmarking atlas-level data integration in single-cell genomics"

### Supplementary Information

#### Supplementary Figures

##### Trajectories and benchmarking results on immune (human)

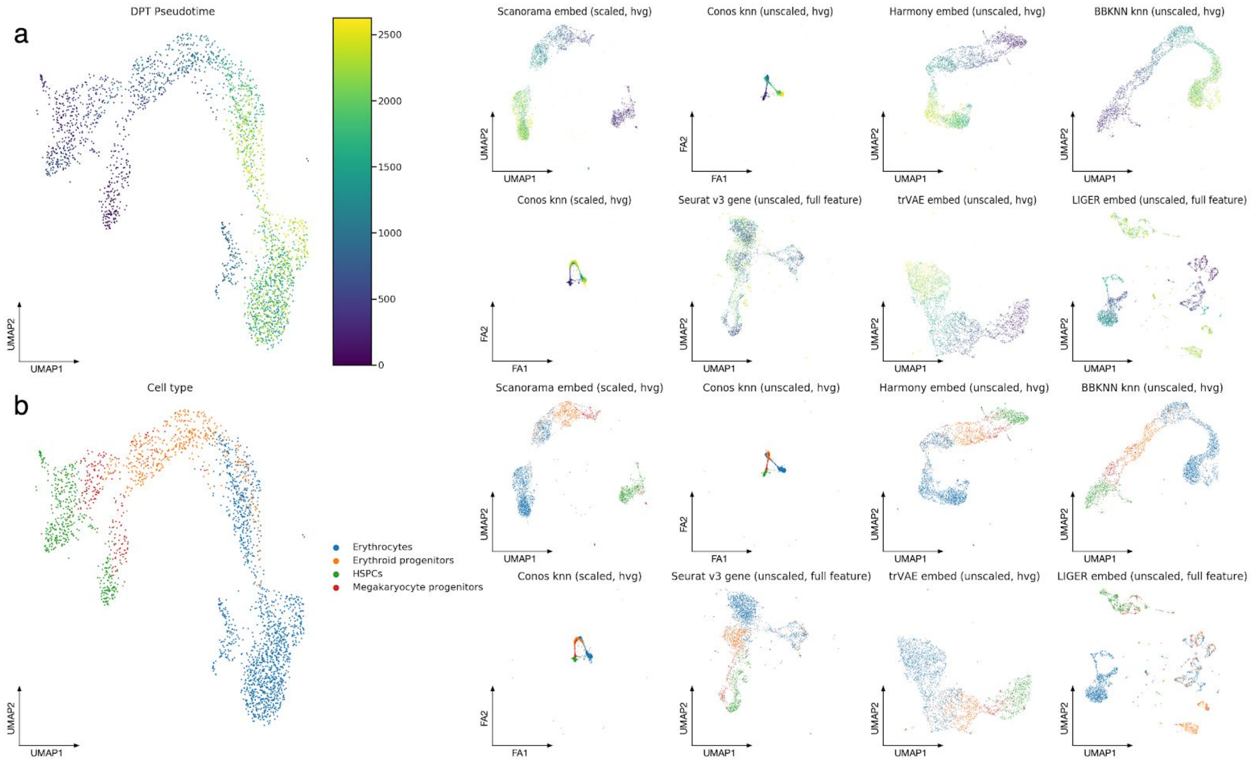


**Supplementary Figure 1: Visualization of the best and worst performers on the immune cell human integration task ordered by overall score on the set of cells belonging to the erythrocyte lineage.** The plots show Force Atlas 2 (Conos) and UMAP (all other methods) layouts for the unintegrated data (left), the top 4 performers (upper rows a and b), and the worst 4 performers (lower rows a and b). (a) shows the order of cells by diffusion pseudotime, while (b) shows cell identity annotations.

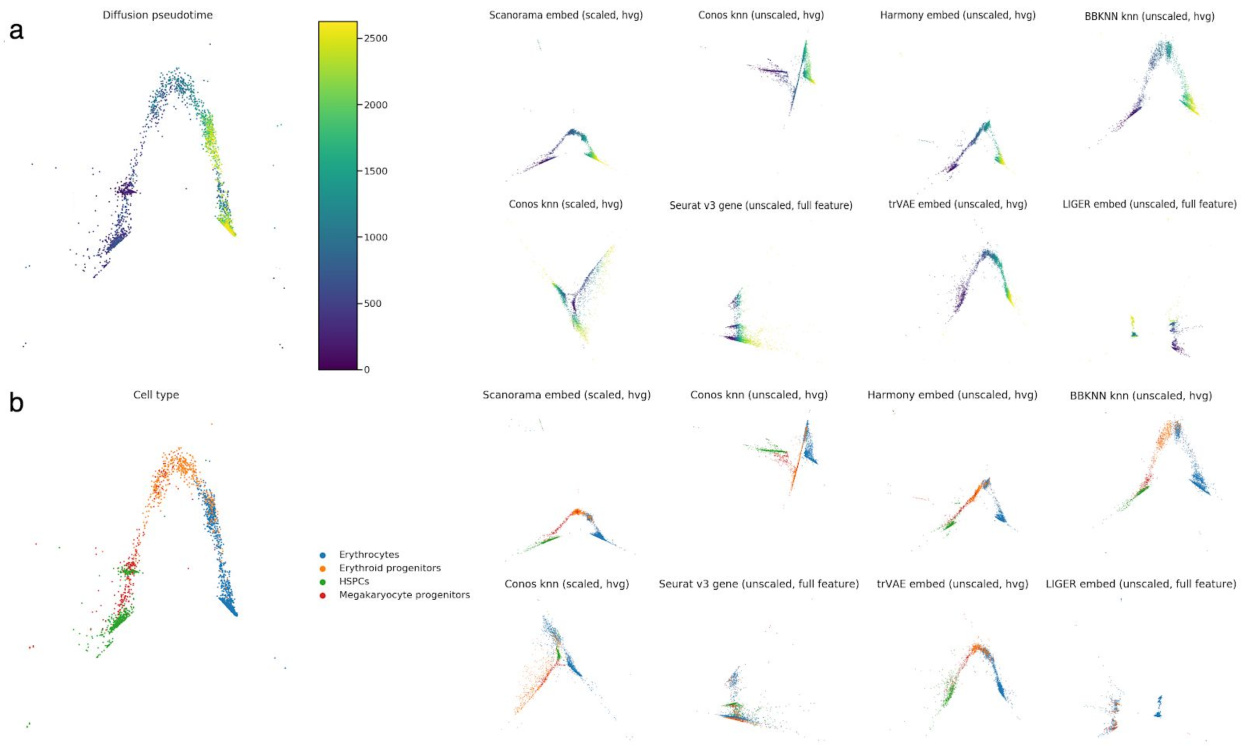


**Supplementary Figure 2: Diffusion maps of diffusion pseudotime (dpt) trajectories on integrated immune cell human data.** Shown are the dpt values of the 4 best (upper row) and 4 worst (lower row) integration methods, ordered by the overall score. (a) shows diffusion maps of the unintegrated data, while the color gradient represents the order of cells by dpt value on the integrated data. (b) shows diffusion maps of the integrated data, where the color gradient represents the dpt value.

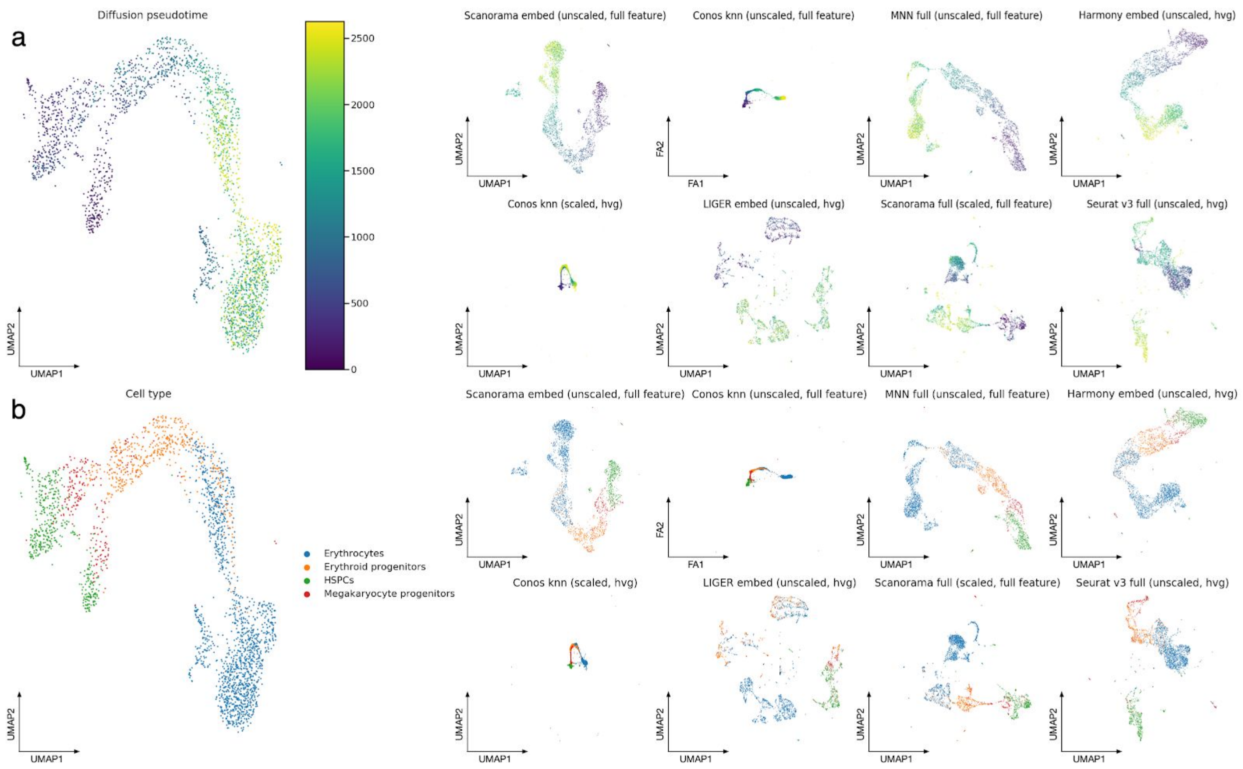


**Supplementary Figure 3: Visualization of the best and worst performers on the immune cell human integration task ordered by trajectory score on the set of cells belonging to the erythrocyte lineage.** The plots show Force Atlas 2 (Conos) and UMAP (all other methods) layouts for the unintegrated data (left), the top 4 performers (upper rows a and b), and the worst 4 performers (lower rows a and b). (a) shows the order of cells by diffusion pseudotime, while (b) shows cell identity annotations.

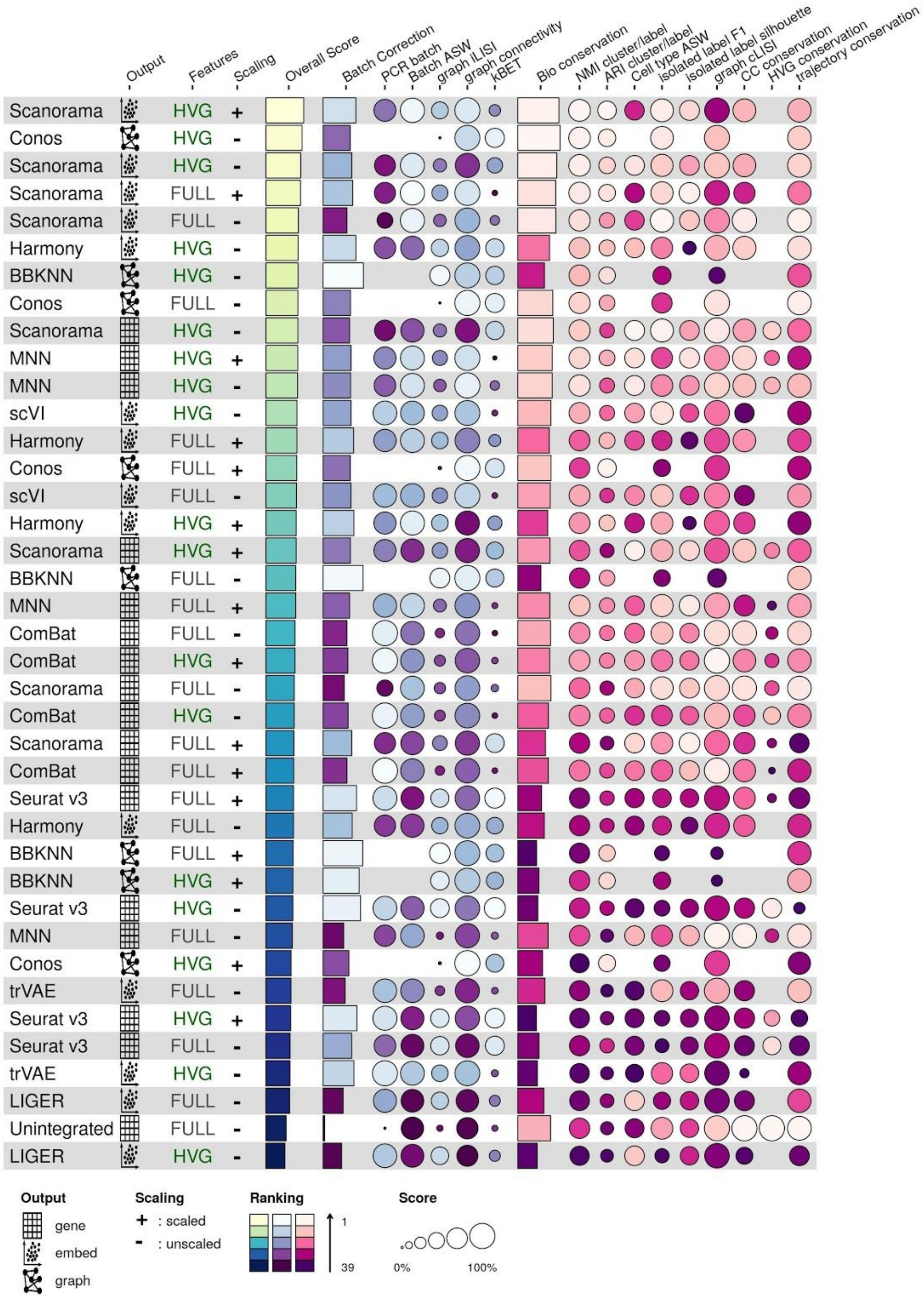


**Supplementary Figure 4: Overview of benchmarking results by overall score for the human immune cell task.** Metrics are divided into batch correction (blue, purple) and bio conservation (pink) categories. Overall scores are computed by a 40:60 weighted mean of these category scores (see **Methods** for further visualization details).

### Trajectories across species

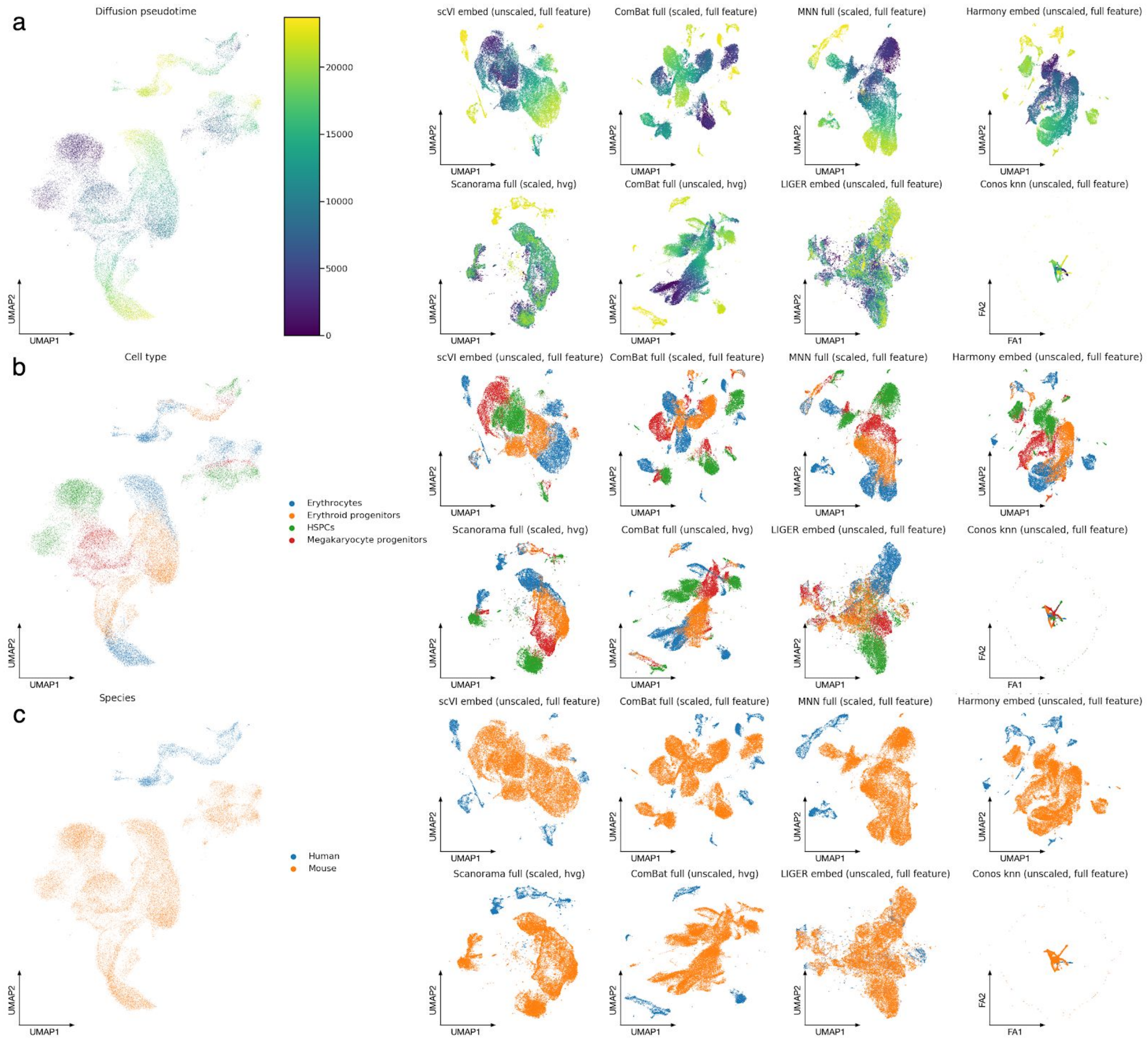


**Supplementary Figure 5: Visualization of the best and worst performers on the immune cell human mouse integration task ordered by trajectory score on the set of cells belonging to the erythrocyte lineage.** The plots show Force Atlas 2 (Conos) and UMAP (all other methods) layouts for the unintegrated data (left), the top 4 performers (upper rows a, b and c), and the worst 4 performers (lower rows a, b and c). Plots are coloured by (a) diffusion pseudotime, (b) cell identity annotations, and (c) species.

### Benchmarking results

#### Immune (human/mouse)

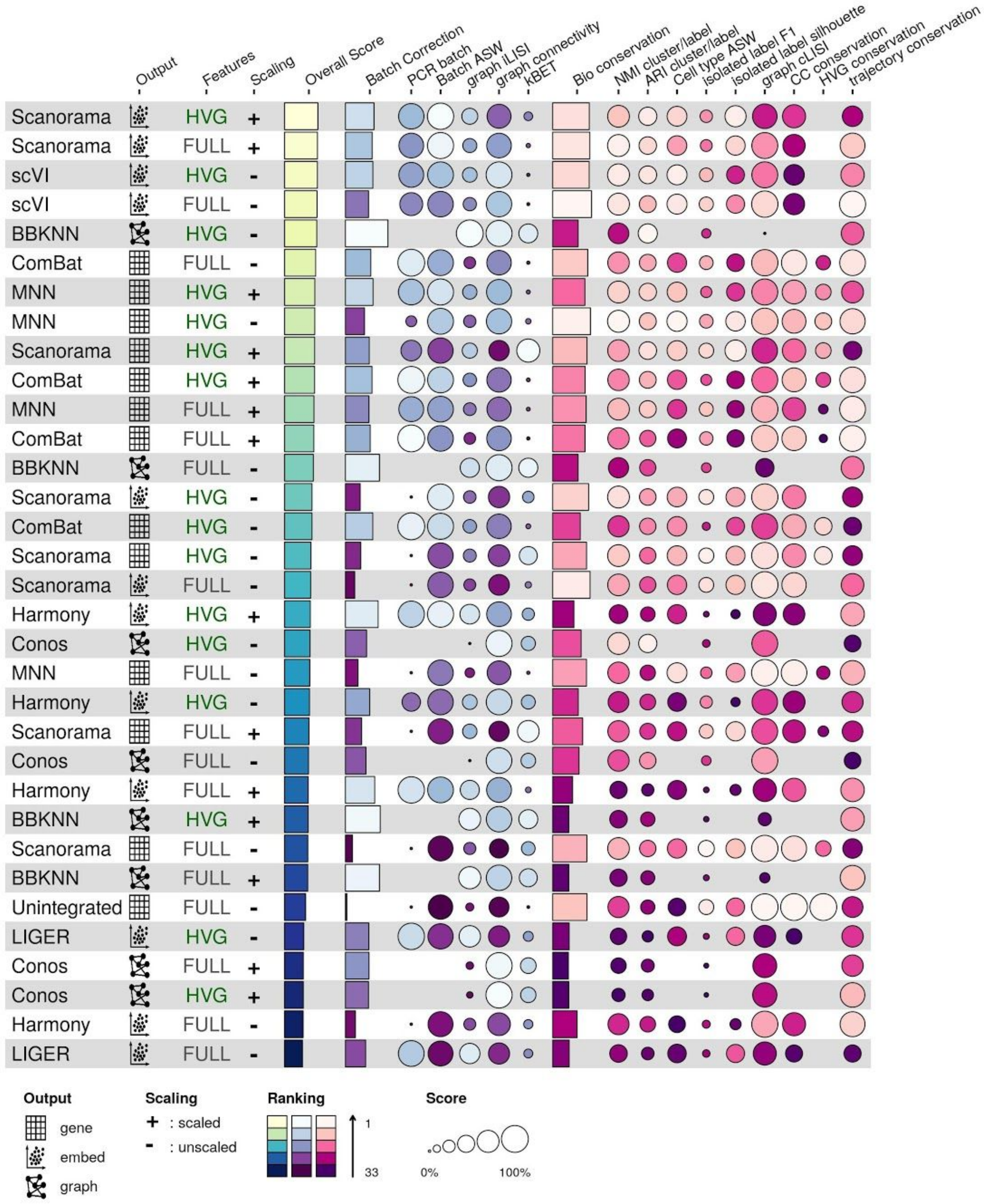


**Supplementary Figure 6: Overview of benchmarking results by overall score for the human/mouse immune cell task.** Metrics are divided into batch correction (blue, purple) and bio conservation (pink) categories. Overall scores are computed by a 40:60 weighted mean of these category scores (see **Methods** for further visualization details). Methods that failed to run are omitted.

#### Simulation 1

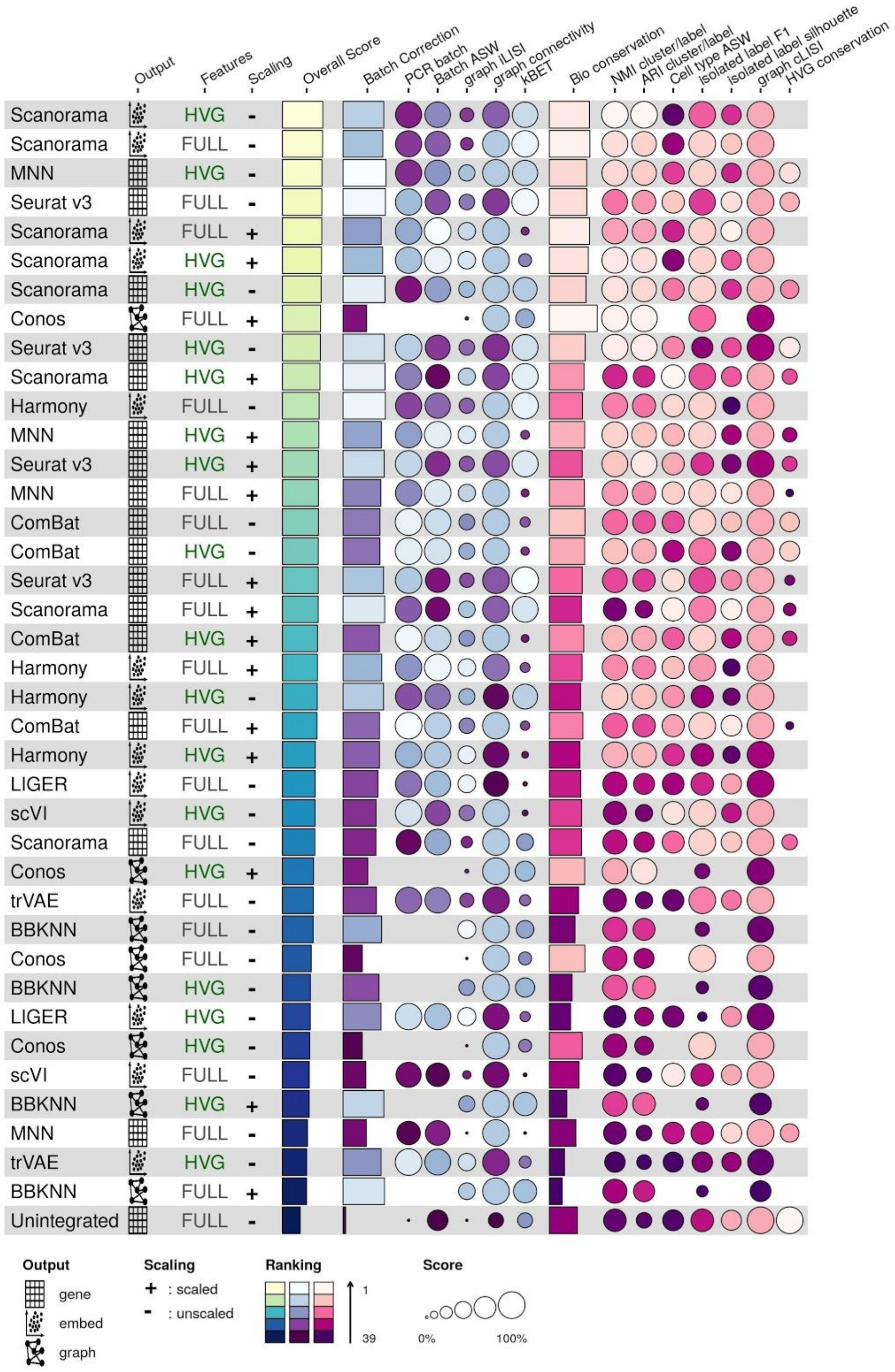

**Supplementary Figure 7: Overview of benchmarking results by overall score for the simulation 1 task.** Metrics are divided into batch correction (blue, purple) and bio conservation (pink) categories. Overall scores are computed by a 40:60 weighted mean of these category scores (see **Methods** for further visualization details).

#### Simulation 2

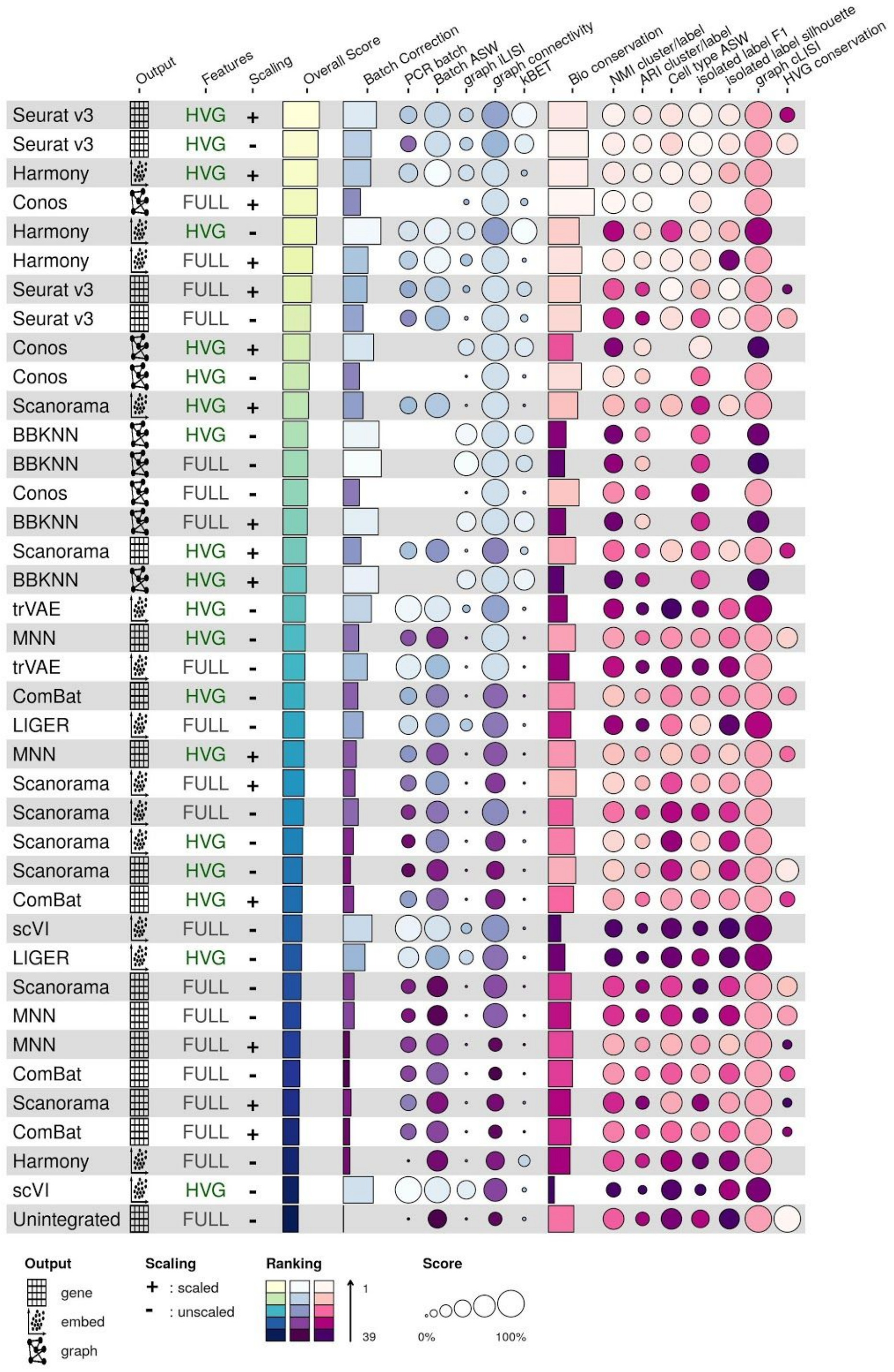

**Supplementary Figure 8: Overview of benchmarking results by overall score for the simulation 2 task.** Metrics are divided into batch correction (blue, purple) and bio conservation (pink) categories. Overall scores are computed by a 40:60 weighted mean of these category scores (see **Methods** for further visualization details).

#### Pancreas

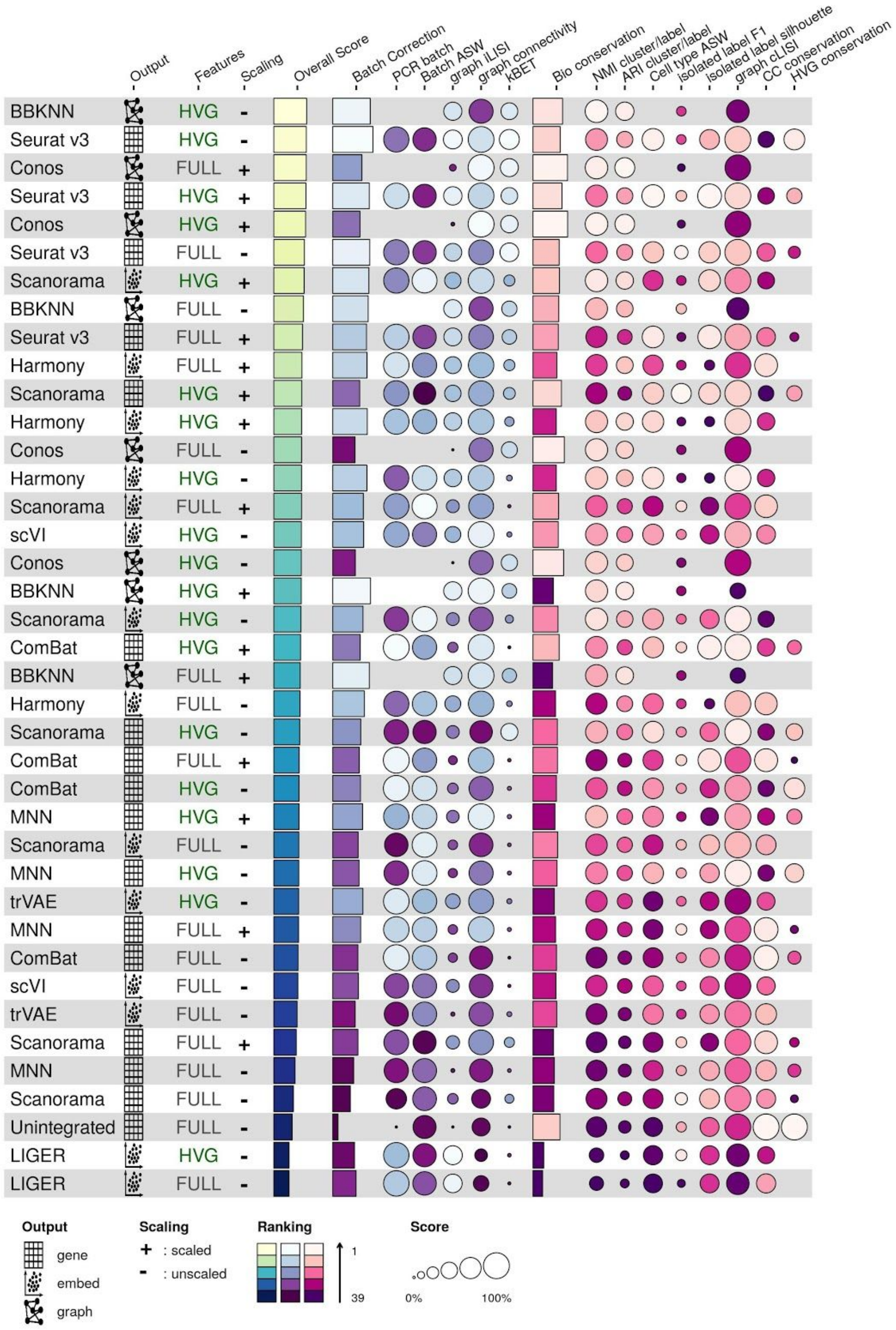

**Supplementary Figure 9: Overview of benchmarking results by overall score for the pancreas task.** Metrics are divided into batch correction (blue, purple) and bio conservation (pink) categories. Overall scores are computed by a 40:60 weighted mean of these category scores (see **Methods** for further visualization details).

#### Lung

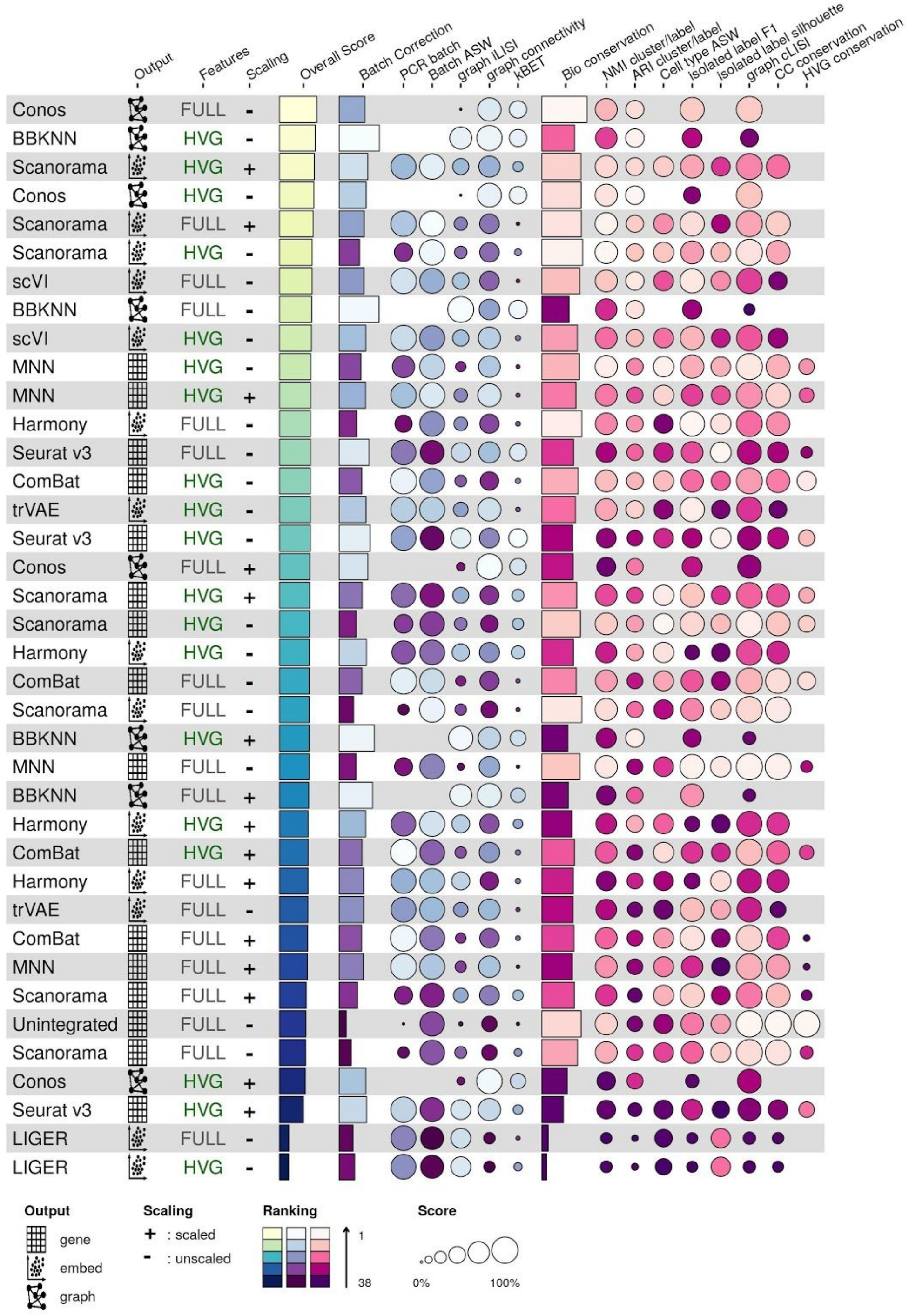

**Supplementary Figure 10: Overview of benchmarking results by overall score for the lung atlas task.** Metrics are divided into batch correction (blue, purple) and bio conservation (pink) categories. Overall scores are computed by a 40:60 weighted mean of these category scores (see **Methods** for further visualization details). Methods that failed to run are omitted.

#### Mouse brain

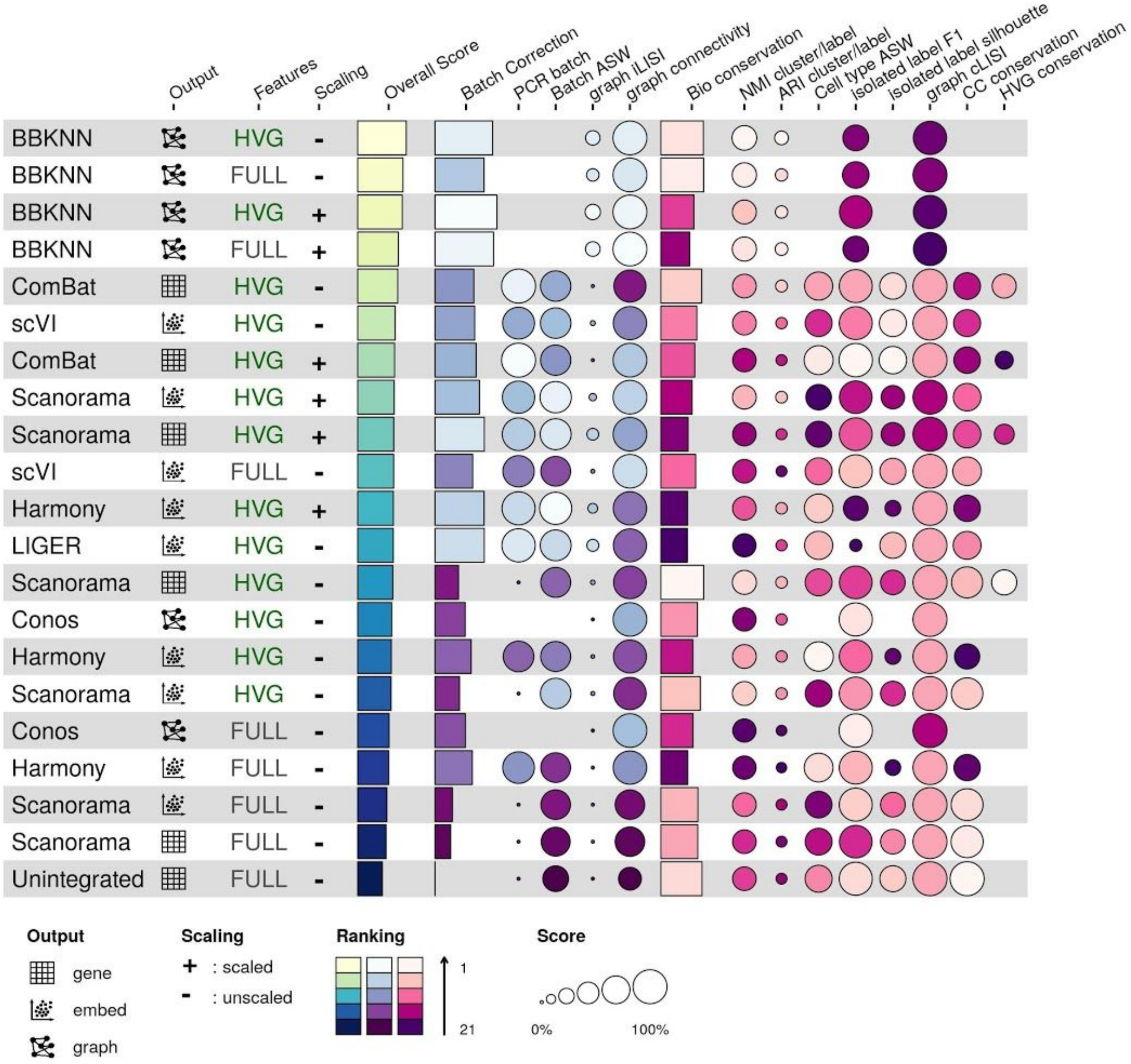


**Supplementary Figure 11: Overview of benchmarking results by overall score for the mouse brain RNA task.** Metrics are divided into batch correction (blue, purple) and bio conservation (pink) categories. Overall scores are computed by a 40:60 weighted mean of these category scores (see **Methods** for further visualization details). Methods that failed to run are omitted. Note that kBET was not run on this task due to computational limitations.

### Embeddings

#### Immune (human)

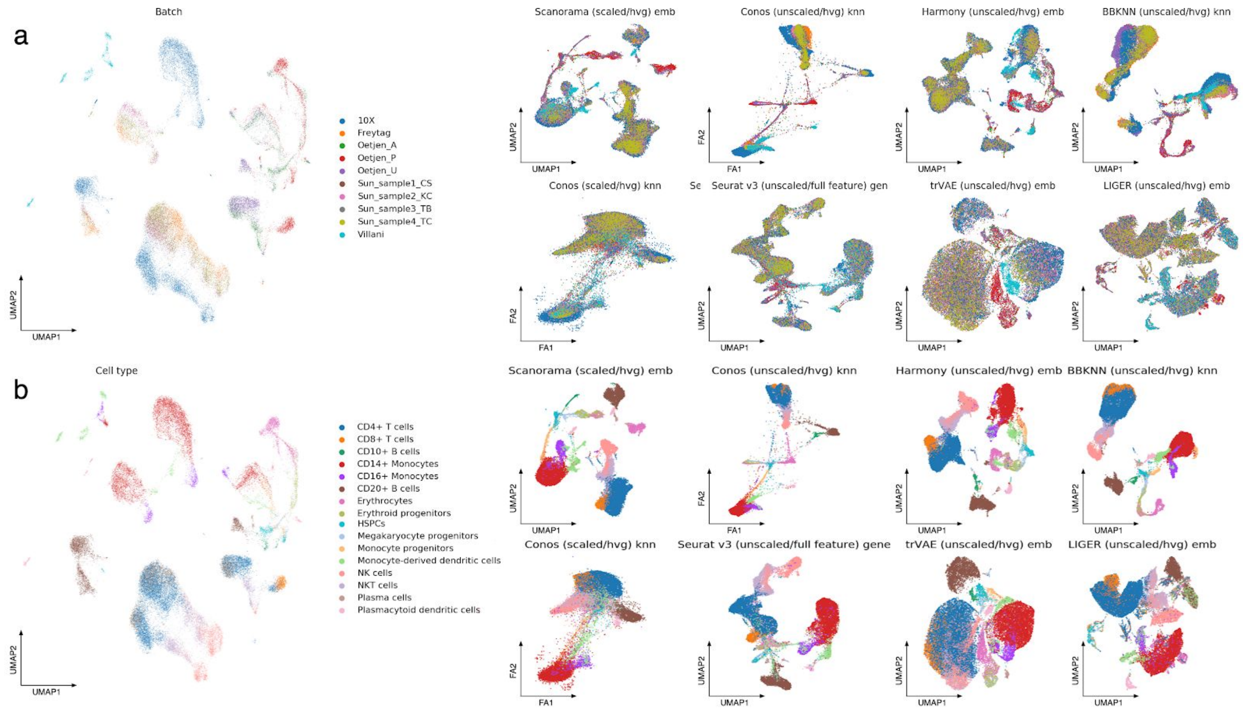


**Supplementary Figure 12: Visualization of the best and worst performers on the immune cell human data.** The plots show Force Atlas 2 (Conos) and UMAP (all other methods) layouts for the unintegrated data (left), the top 4 performers (upper rows a and b), and the worst 4 performers (lower rows a and b). Plots are coloured by (a) batch labels, and (b) cell identity annotations.

#### Immune (human/mouse)

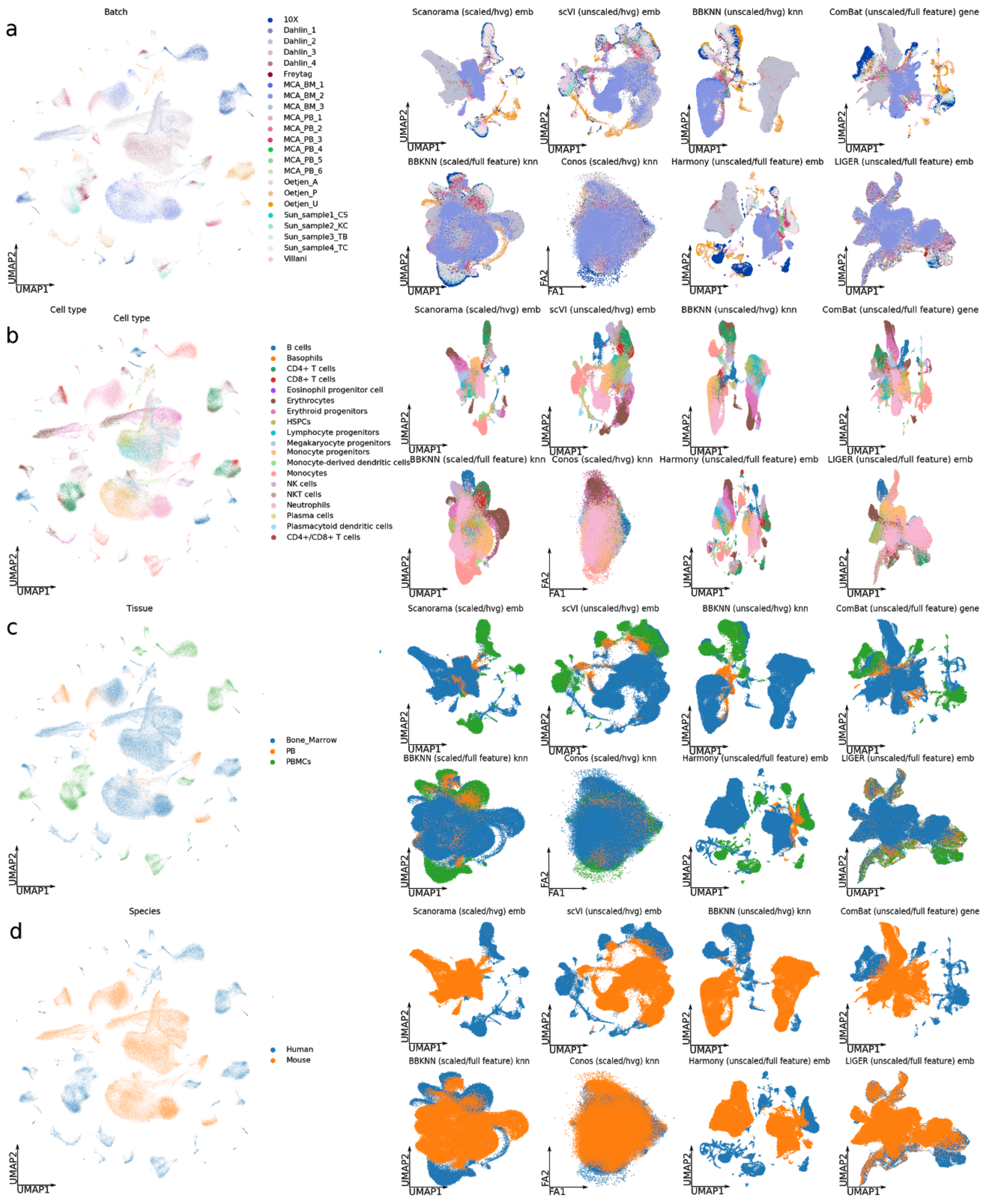


**Supplementary Figure 13**: **Visualization of the best and worst performers on the immune cell human mouse integration task.** The plots show Force Atlas 2 (Conos) and UMAP (all other methods) layouts for the unintegrated data (left), the top 4 performers (upper rows a and b), and the worst 4 performers (lower rows a and b). Plots are coloured by (a) batch labels, (b)

cell identity annotations, (c) tissue, and (d) species. Tissue abbreviations are: PB - peripheral blood, PBMCs - peripheral blood mononuclear cells.

#### Simulation 1

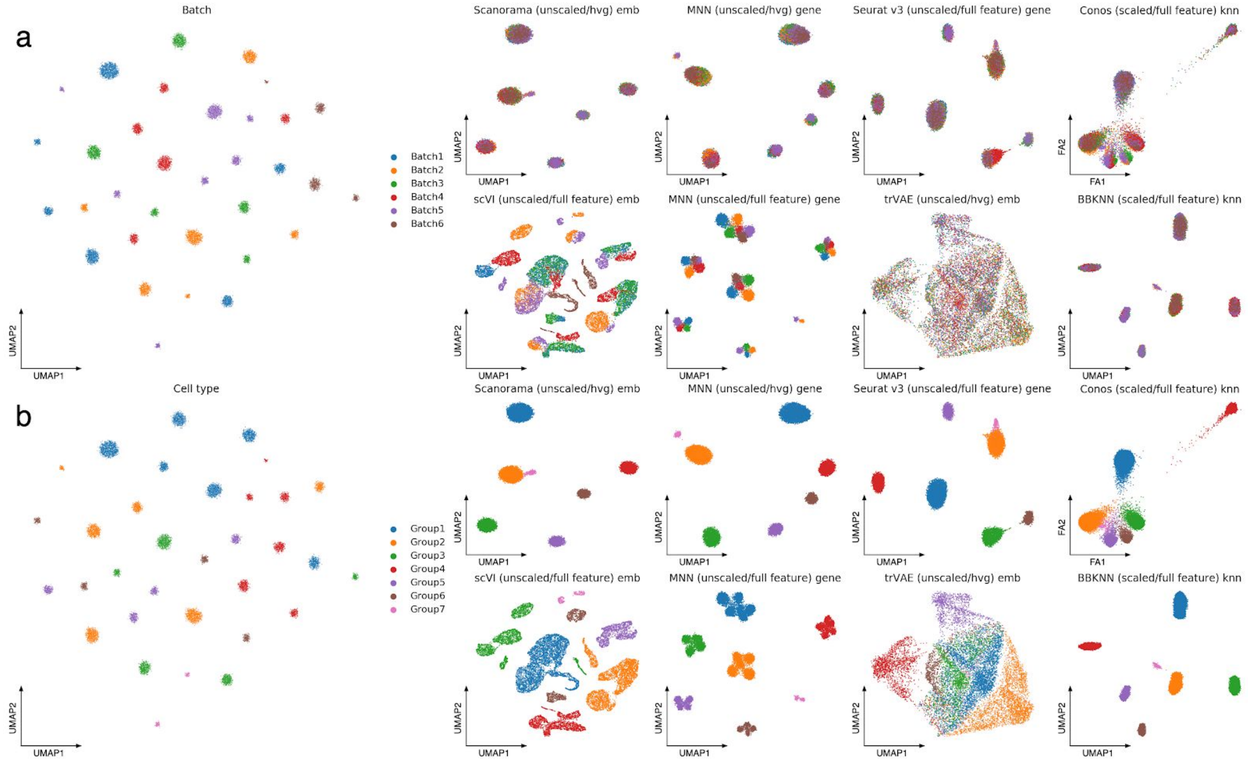


**Supplementary Figure 14: Visualization of the best and worst performers on the simulation 1 task.** The plots show Force Atlas 2 (Conos) and UMAP (all other methods) layouts for the unintegrated data (left), the top 4 performers (upper rows a and b), and the worst 4 performers (lower rows a and b). Plots are coloured by (a) batch labels, and (b) cell identity annotations.

#### Simulation 2

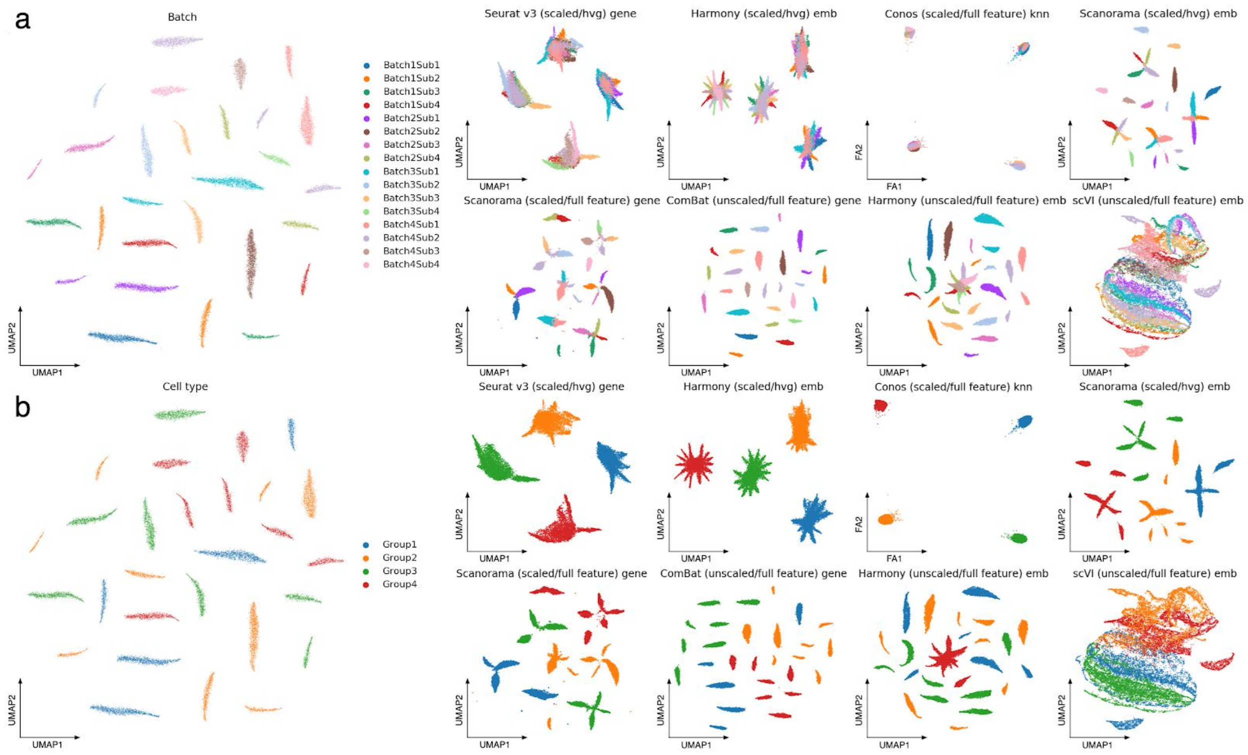


**Supplementary Figure 15: Visualization of the best and worst performers on the simulation 2 task.** The plots show Force Atlas 2 (Conos) and UMAP (all other methods) layouts for the unintegrated data (left), the top four performers (upper rows a and b), and the worst four performers (lower rows a and b). Plots are coloured by (a) batch labels, and (b) cell identity annotations.

#### Pancreas

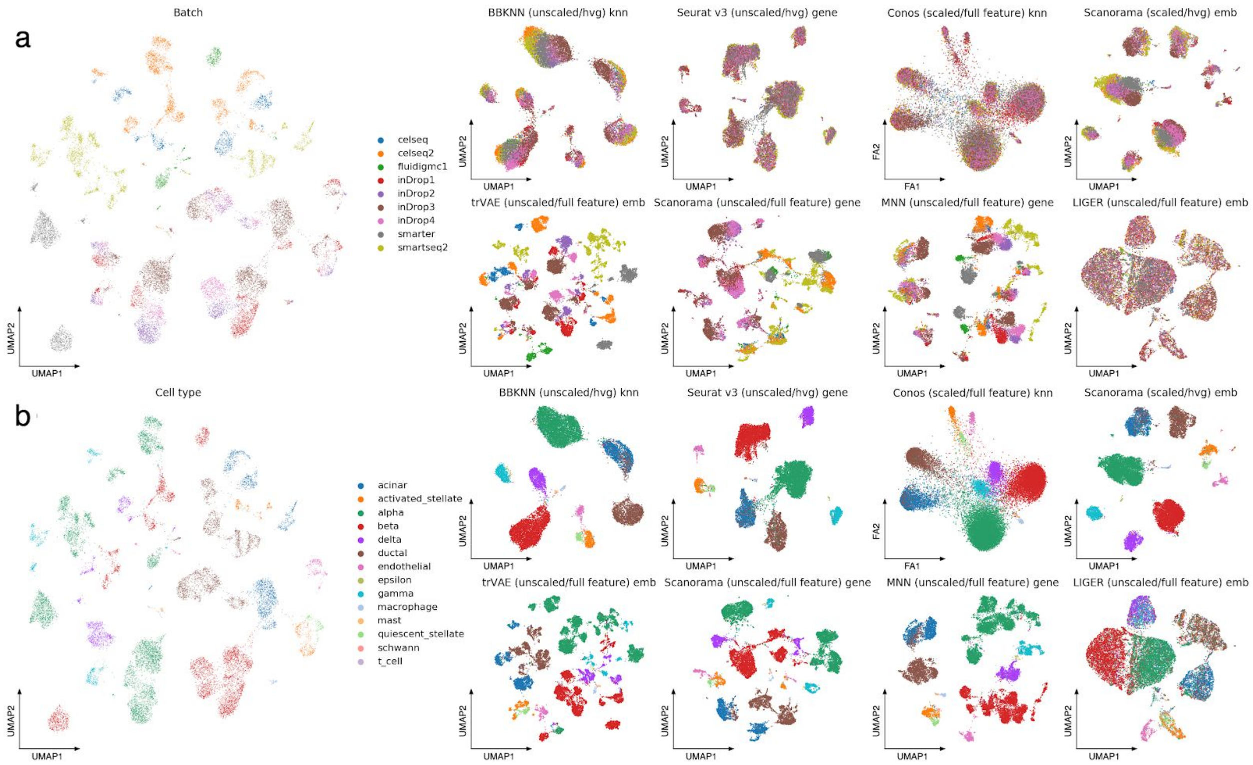


**Supplementary Figure 16: Visualization of the best and worst performers on the pancreas integration task.** The plots show Force Atlas 2 (Conos) and UMAP (all other methods) layouts for the unintegrated data (left), the top 4 performers (upper rows a and b), and the worst 4 performers (lower rows a and b). Plots are coloured by (a) batch labels, and (b) cell identity annotations.

#### Lung

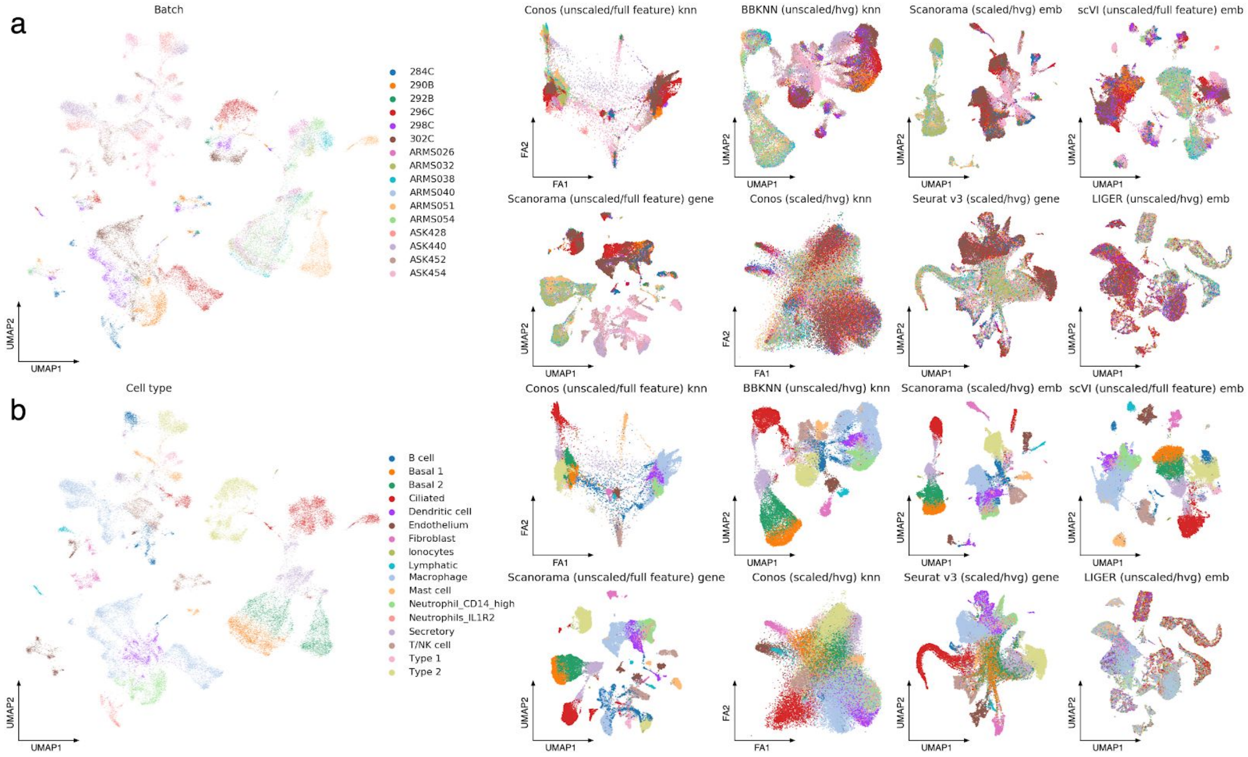


**Supplementary Figure 17: Visualization of the best and worst performers on the lung atlas integration task.** The plots show Force Atlas 2 (Conos) and UMAP (all other methods) layouts for the unintegrated data (left), the top 4 performers (upper rows a and b), and the worst 4 performers (lower rows a and b). Plots are coloured by (a) batch labels and (b) cell identity annotations.

#### Mouse Brain

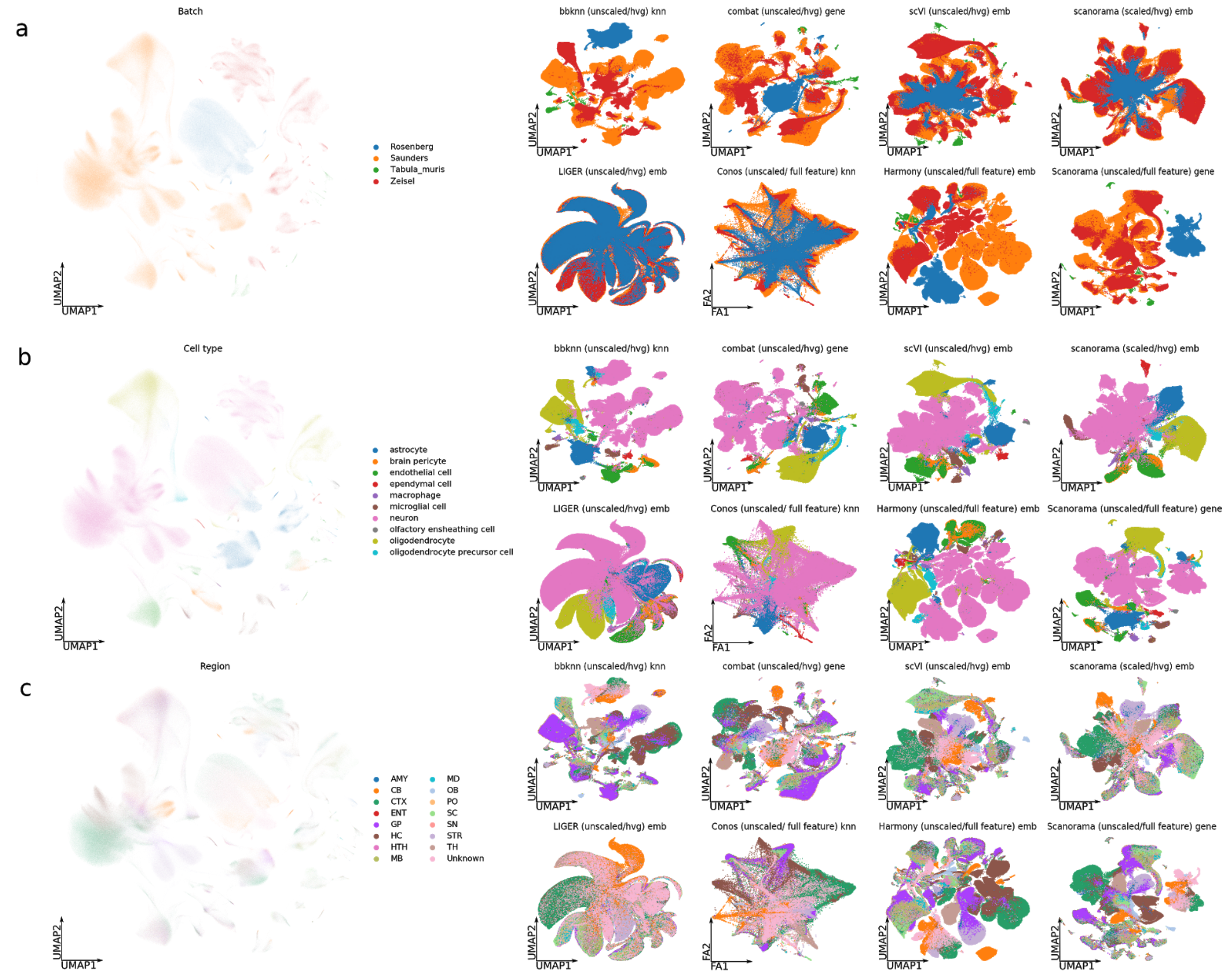


**Supplementary Figure 18: Visualization of the best and worst performers on the mouse brain RNA integration task.** The plots show Force Atlas 2 (Conos) and UMAP (all other methods) layouts for the unintegrated data (left), the top 4 performers (upper rows a, b, and c), and the worst 4 performers (lower rows a, b, and c). Plots are coloured by (a) batch labels, (b) cell identity annotations, and (c) brain regions. Brain region abbreviations are: AMY - amygdala, HC - hippocampus, TH - thalamus, HTH - hypothalamus, CTX - cortex, OB - olfactory bulb, STR - striatum, CB - cerebellum, MB - midbrain, MD - medulla, SN - substantia nigra, ENT - entopeduncular nucleus, GP - globus pallidus and nucleus basalis, PO - pons, and SC - spinal cord (unknown regions could not be inferred in the original publication of Rosenberg *et al.*[1]).

### Performance summary

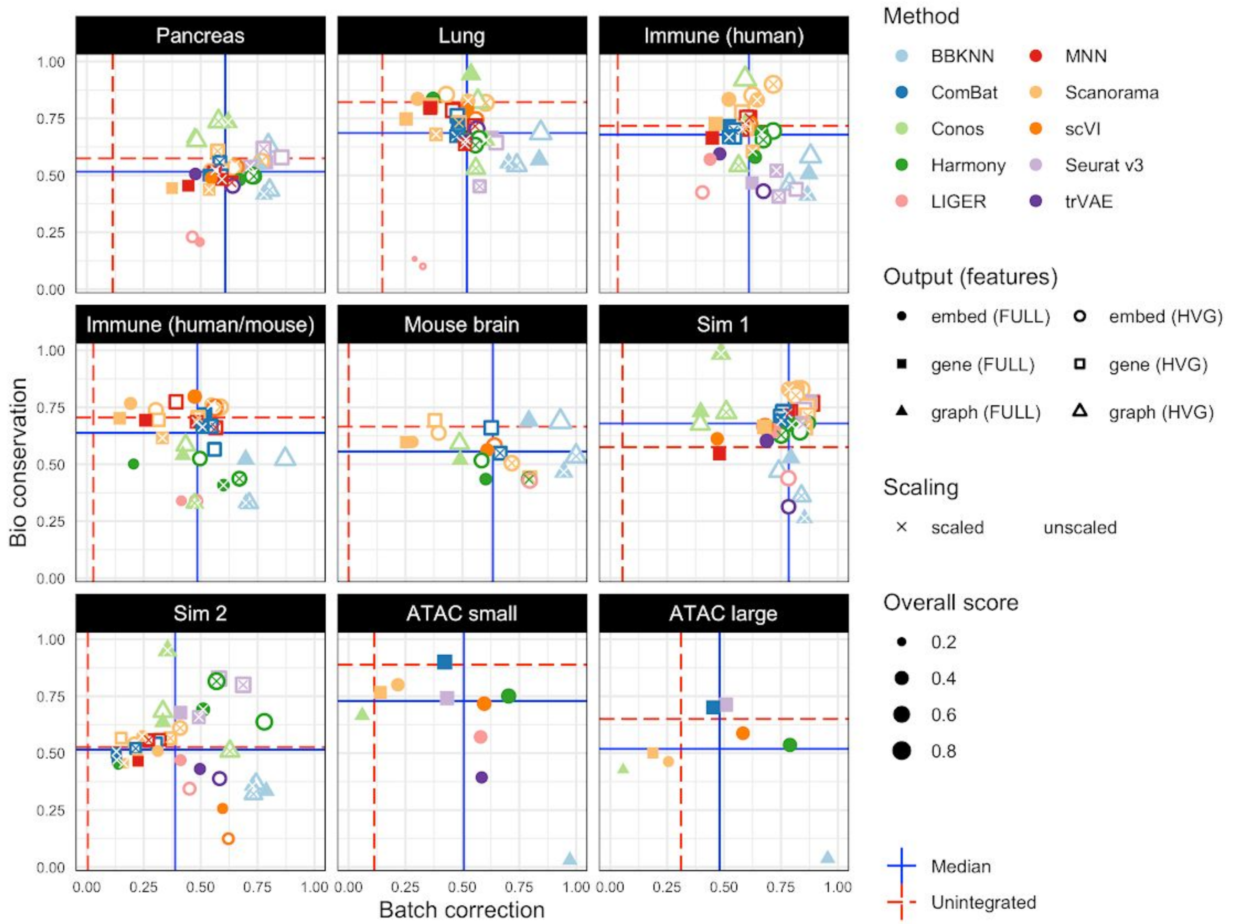


**Supplementary Figure 19: Scatter plots summarizing integration performance on all tasks.** The x-axis shows the overall batch correction score and the y-axis shows the overall bio conservation score. Each point is an individual integration run. Point colour indicates method, size the overall score and shape the output type (embed, gene, graph). Filled points use the full feature set while unfilled points use selected highly variable genes. Points marked with a cross use scaled features. Horizontal lines indicate reference points. Red dashed lines show performance calculated on the unintegrated dataset and solid blue lines the median performance across methods for each dataset.

### Scalability

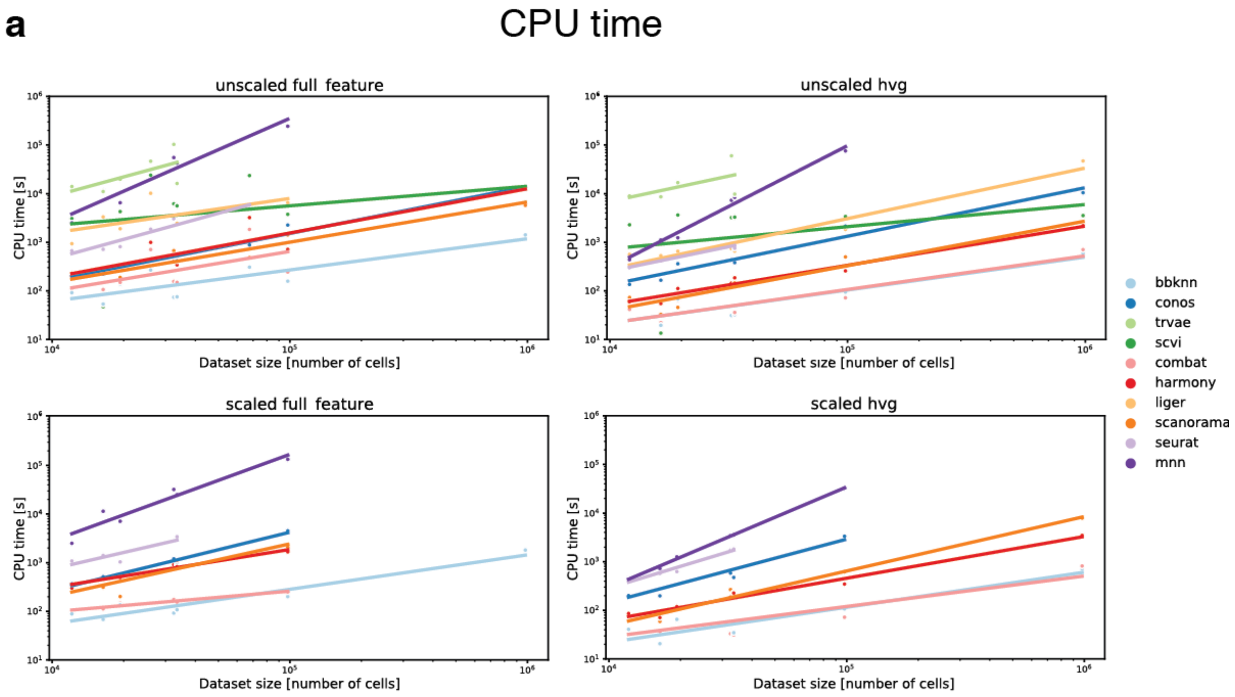


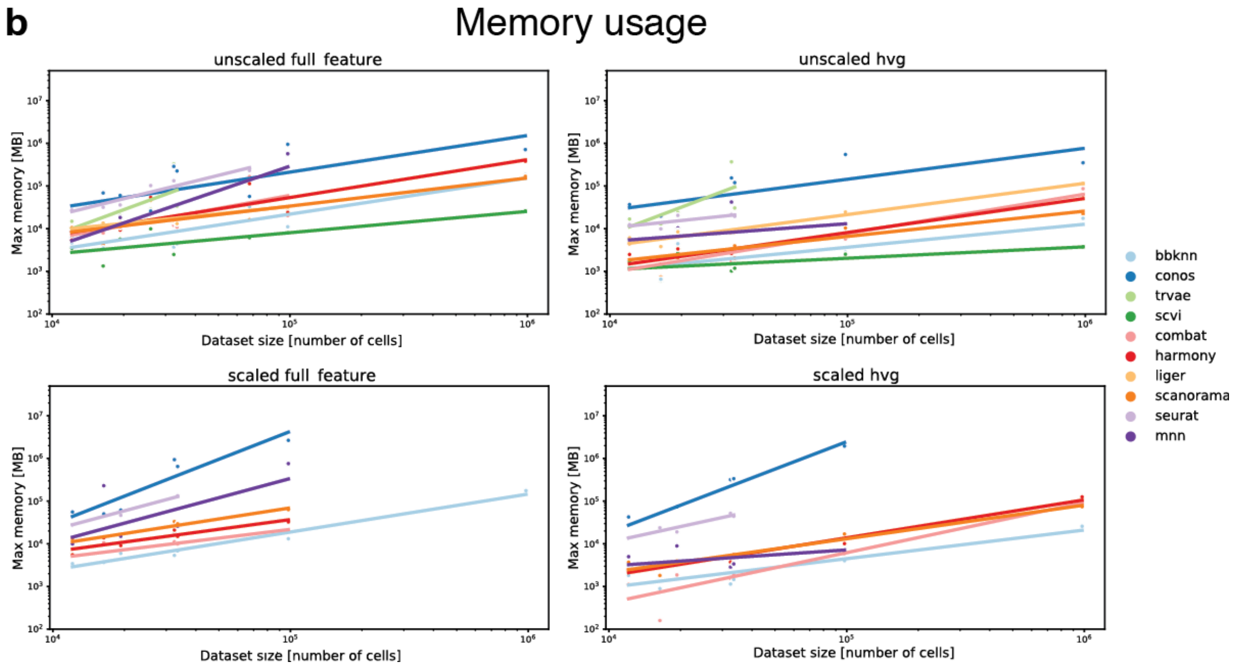


**Supplementary Figure 20: Scalability of each data integration method, separated by preprocessing variant.** a) CPU time in seconds for each method (colored dots) and data integration task. b) Maximum memory usage for each method and scenario. Colored lines denote linear fit of log-scaled time/memory vs log-scaled dataset size for each data integration method and pre-processing combination.

#### Usability

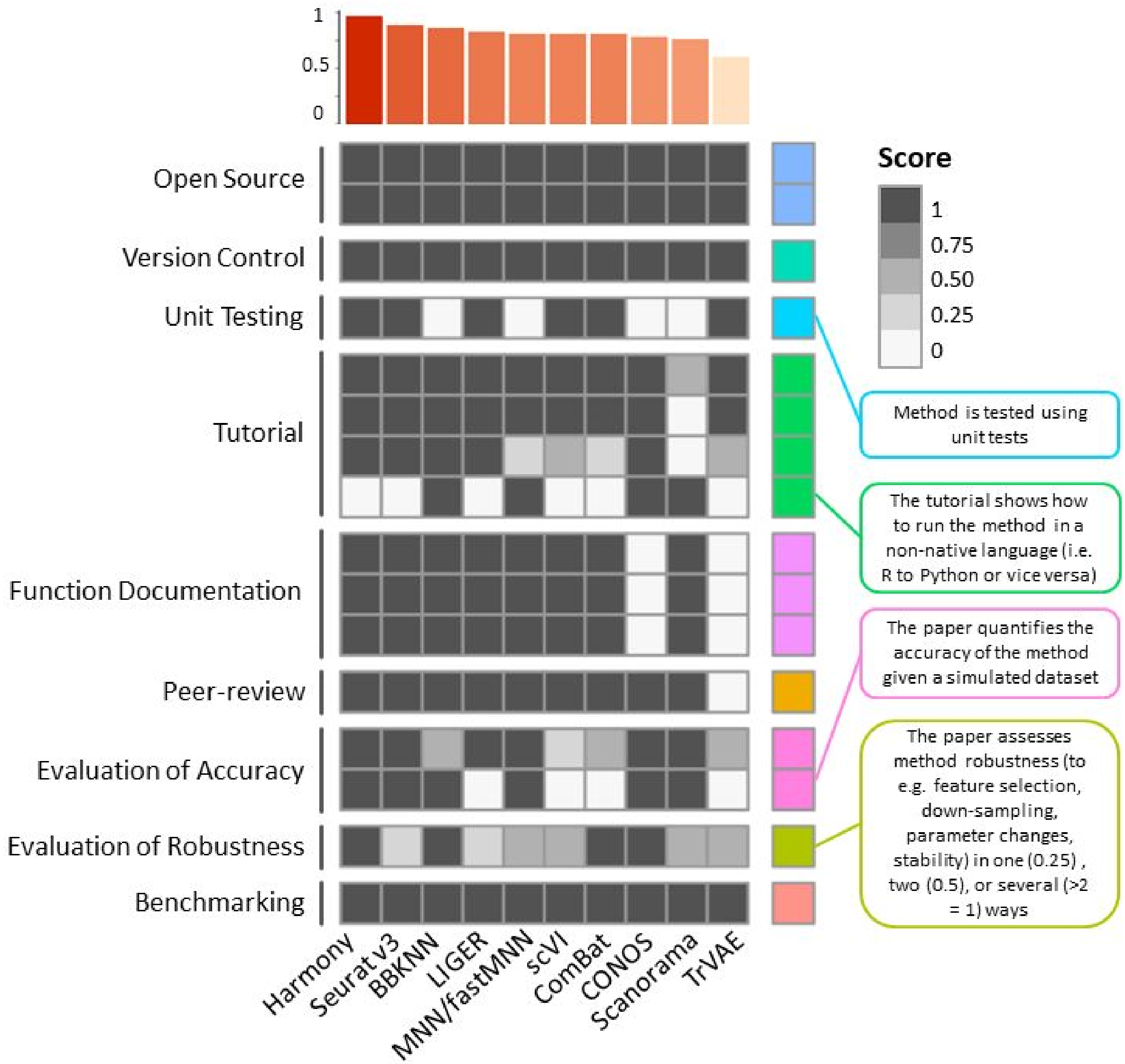


**Supplementary Figure 21: Usability assessment of data integration methods.** The usability of each data integration method was assessed via nine categories (labels on the left), plotted as a heatmap, and ordered by overall usability score. On the right-hand side criteria with poor scores across methods are highlighted for each category. The overall usability score was computed as the mean of all category scores and plotted on top in a barplot.

### ATAC results

#### ATAC small benchmarking results

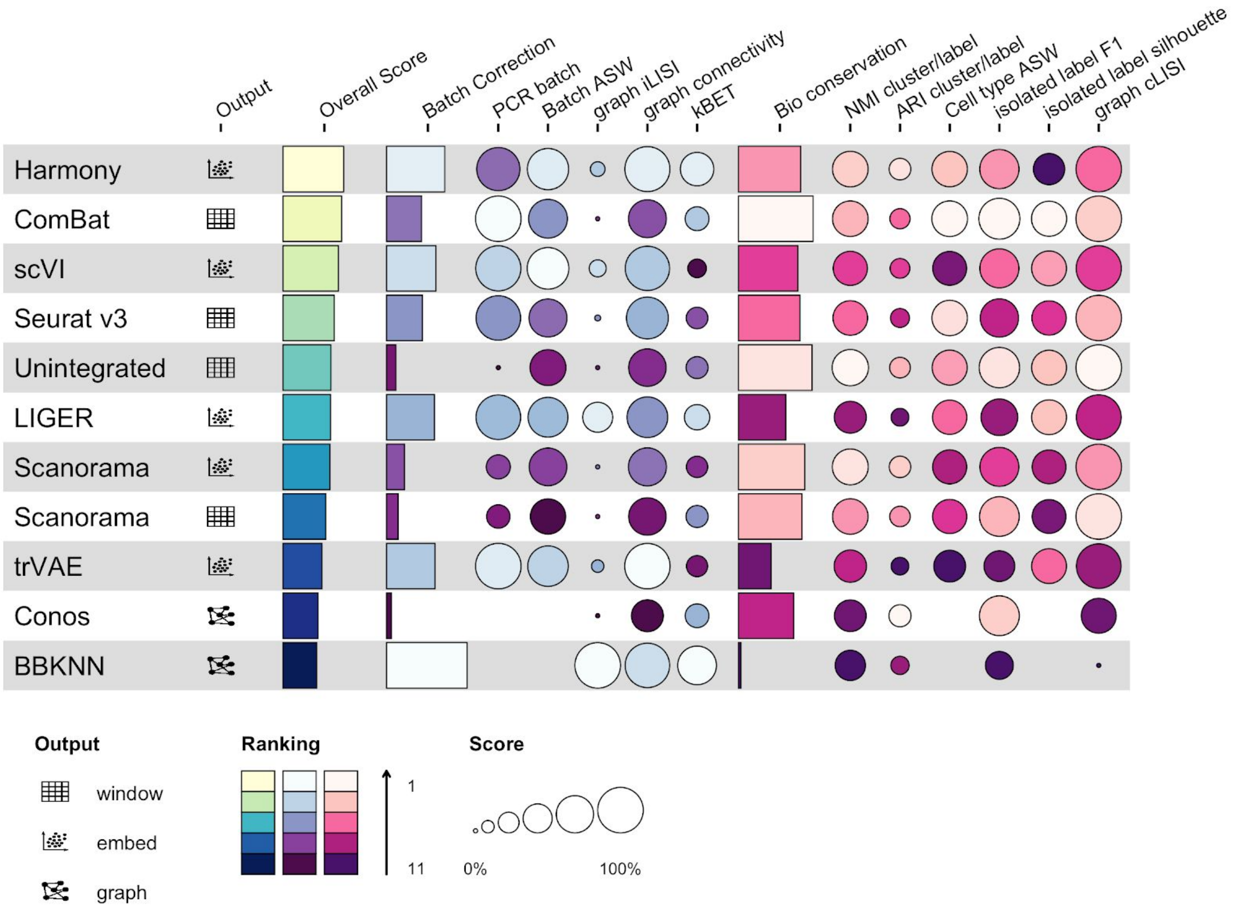


**Supplementary Figure 22: Benchmarking results for the small mouse brain task based on scATAC-seq.** Metrics are divided into batch correction (blue, purple) and bio conservation (pink) categories (see **Methods** for further visualization details). Overall scores are computed by a 40:60 weighted mean of these category scores. Methods that failed to run are omitted.

### ATAC small embeddings

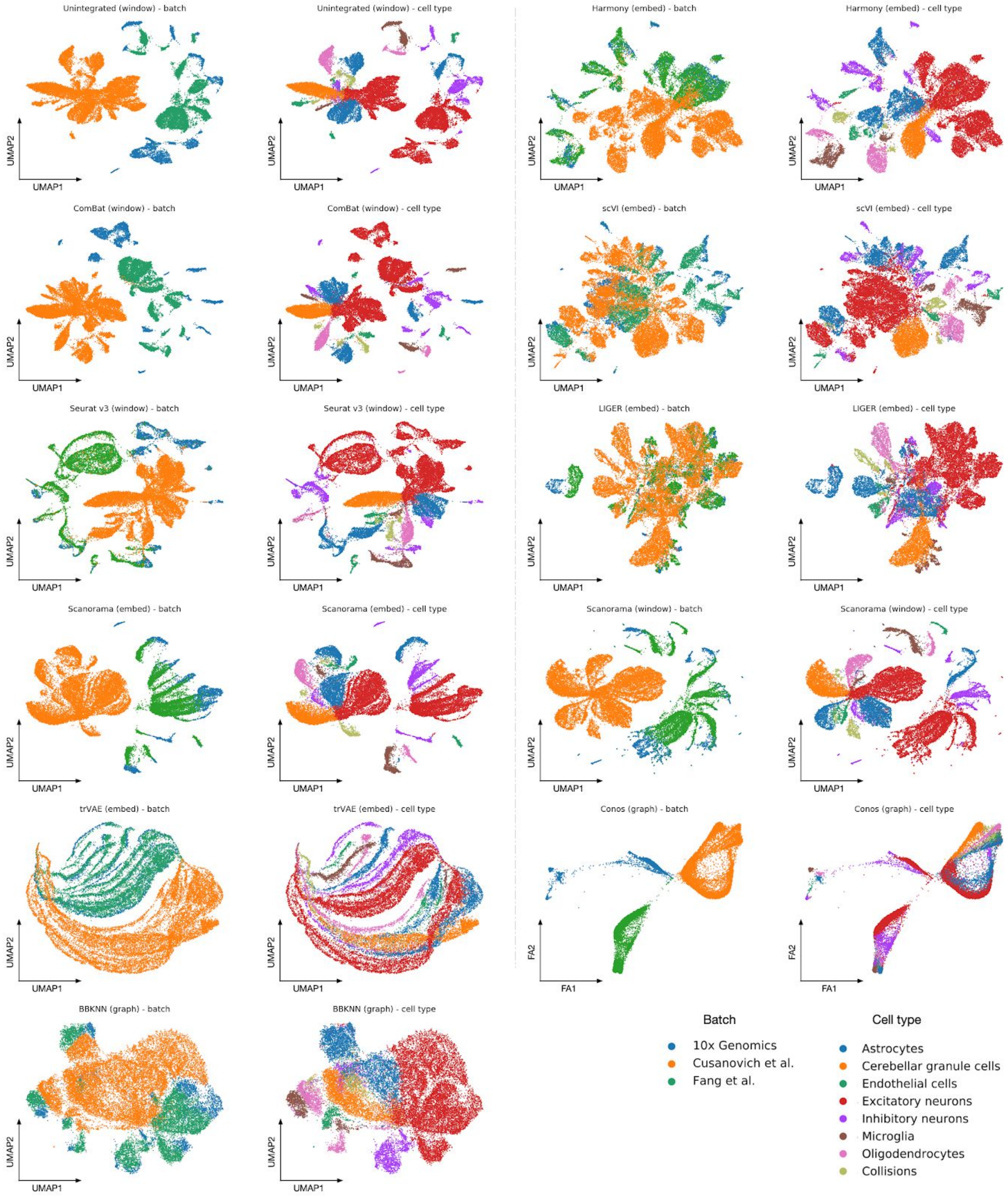


**Supplementary Figure 23: Visualisation of all small ATAC tasks.** The plots show Force Atlas 2 (Conos) and UMAP (all other methods) layouts for the unintegrated scenario and the best method to the worst method from left to right then top to the bottom. For each method, there are two plots colored by batch labels and cell identity annotations.

#### ATAC large embeddings

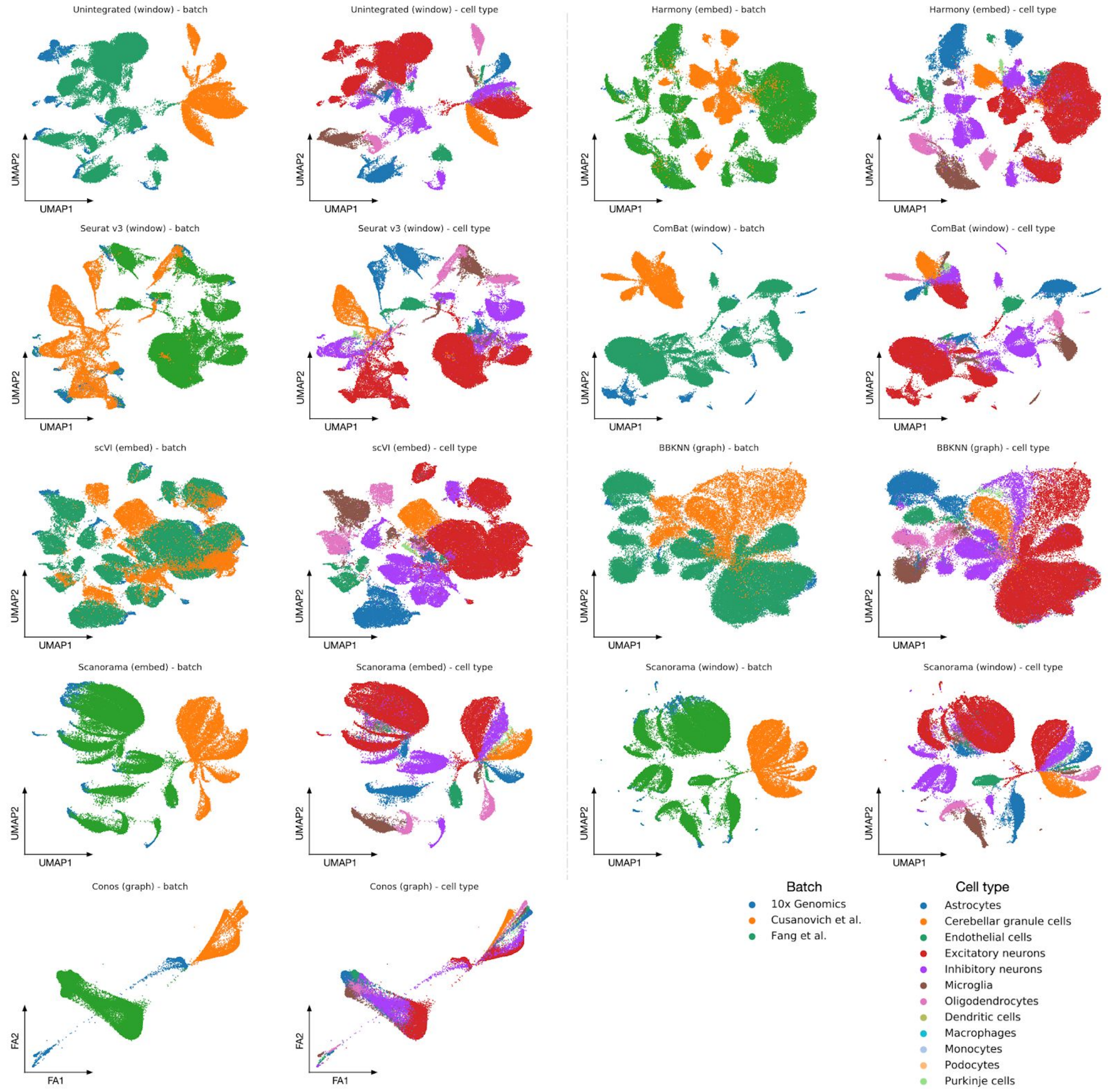


**Supplementary Figure 24: Visualisation of all large ATAC tasks.** The plots show Force Atlas 2 (Conos) and UMAP (all other methods) layouts for the unintegrated scenario and the best method to the worst method from left to right then top to the bottom. For each method, there are two plots colored by batch labels and cell identity annotations.

### iLISI comparison

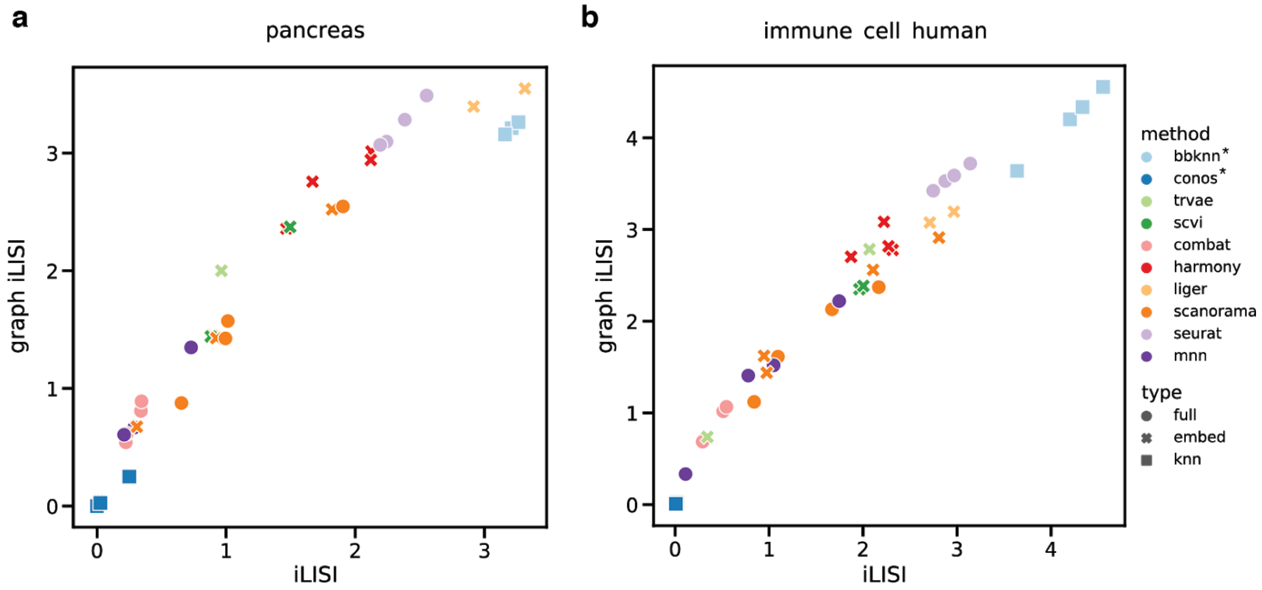


**Supplementary Figure 25: Comparison of graph iLISI and iLISI scores** - All scores are unscaled, but shifted by -1. For graph-based output (marked with a star in the legend and a filled square in the plot), iLISI does not work because these methods do not provide a Euclidean distance measure. For visualisation, the results for graph-based methods are the same on both x- and y-axis, but were computed with graph iLISI. We compared the two metrics in the pancreas (a) and immune cell human (b) data scenarios. Both scores correlate well on full- and embedding-based data integration methods (circles and crosses), *i.e.* Pearson correlation coefficient is 0.978 for the pancreas task and 0.984 for the immune cell human task.

### Trajectories - addendum

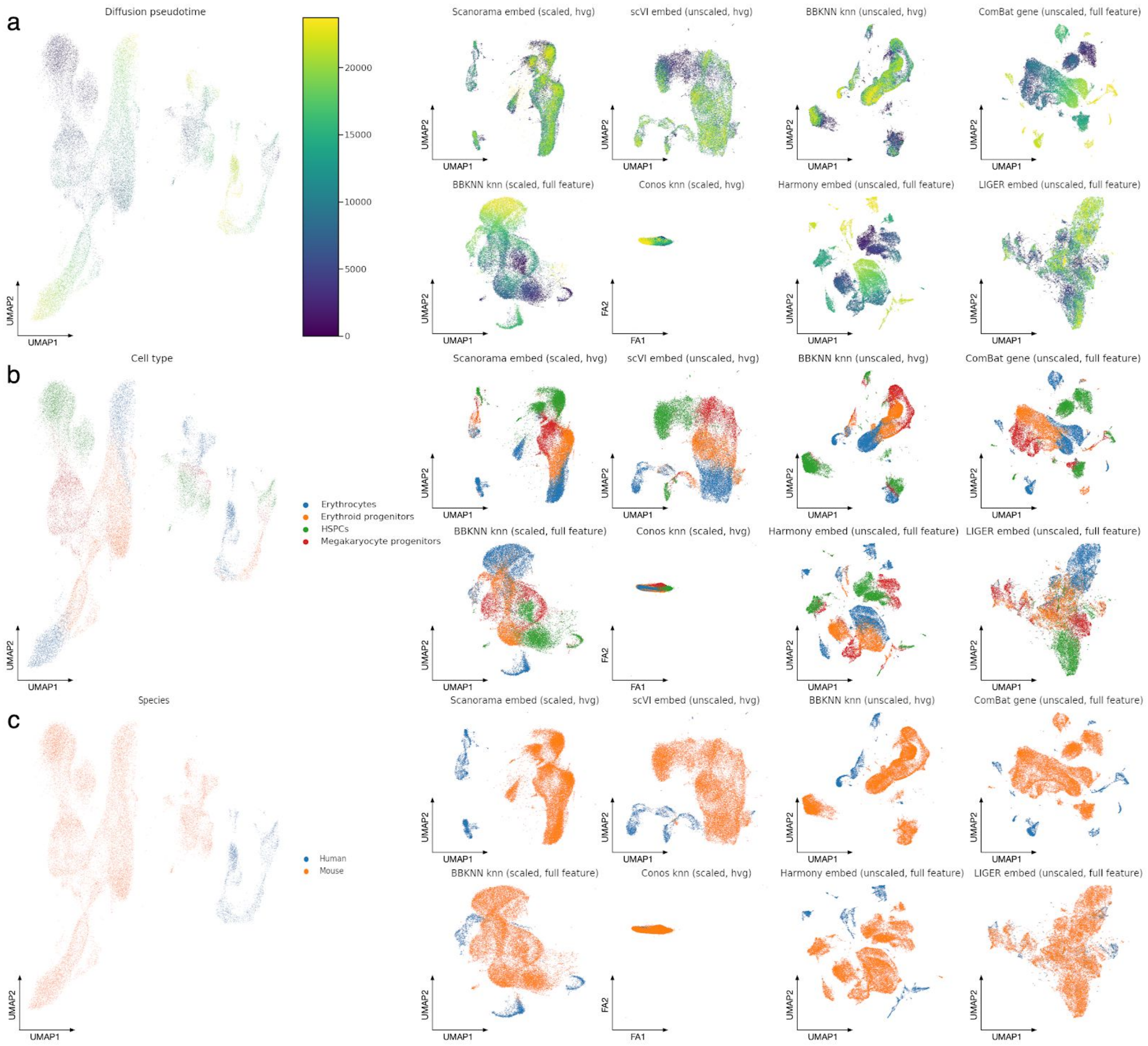


**Supplementary Figure 26: Visualization of the best and worst performers on the immune cell human mouse integration task ordered by overall score.** The plots show Force Atlas 2 (Conos) and UMAP (all other methods) layouts for the unintegrated data (left), the top 4 performers (upper rows a, b and c), and the worst 4 performers (lower rows a, b and c). Plots are coloured by (a) diffusion pseudotime, (b) cell identity annotations, and (c) species.

### Supplementary Table 1: Data integration methods

**Supplementary Table 1**: **Data integration methods available in order of first preprint publication.** Methods that perform only time-series data integration are omitted. The collection is (based on manual literature review and scrna-tools.org[2]; (last retrieved: Feb 2020)

| Method | Method principle | DOI/arXiv ID/url | Github | First preprint Date |
|---|---|---|---|---|
| MNN | Mutual nearest neighbours | 10.1038/nbt.4091 | https://github.com/chriscainx/mnnpy<br><br>https://github.com/MarioniLab/scran | July 18, 2017 |
| Seurat v2 | Canonical correlation analysis (CCA) | 10.1038/nbt.4096 | https://github.com/satijalab/seurat | July 18, 2017 |
| SAUCIE | Sparse, regularized autoencoder | 10.1038/s41592-019-0576-7 | https://github.com/KrishnaswamyLab/SAUCIE/ | December 19, 2017 |
| scVI | Conditional variational autoencoder | 10.1038/s41592-018-0229-2 | https://github.com/YosefLab/scVI | March 30, 2018 |
| Scanorama | SVD + Mutual nearest neighbours | 10.1038/s41587-019-0113-3 | https://github.com/brianhie/scanorama | July 17, 2018 |
| BBKNN | KNN graph integration | 10.1093/bioinformatics/btz625 | https://github.com/Teichlab/bbknn | August 22, 2018 |
| scMerge | Factor analysis model on stably expressed genes | 10.1073/pnas.1820006116 | https://github.com/SydneyBioX/scMerge | September 12, 2018 |
| CONOS | PCA + KNN integration | 10.1038/s41592-019-0466-z | https://github.com/hms-dbmi/conos | November 2, 2018 |

| | | | | |
|---|---|---|---|---|
| LIGER | Integrative non-negative matrix factorization | 10.1101/459891 | https://github.com/MacoskoLab/liger | November 2, 2018 |
| Seurat v3 | CCA + Mutual nearest neighbours | 10.1016/j.cell.2019.05.031 | https://github.com/satijalab/seurat | November 02, 2018 |
| Harmony | PCA + clustering-based correction | 10.1038/s41592-019-0619-0 | https://github.com/immunogenomics/harmony | November 04, 2018 |
| scGen | Conditional variational autoencoder (cell identity labels required) | 10.1038/s41592-019-0494-8 | https://github.com/theislab/scgen | November 29, 2018 |
| RISC | Principal component regression | 10.1101/483297 | N/A | November 29, 2018 |
| scAlign | Bidirectional mapping through deep learning | 10.1186/s13059-019-1766-4 | https://github.com/quon-titative-biology/scAlign | December 22, 2018 |
| scPopCorn | Simultaneous optimisation of subpopulations across samples | 10.1016/j.cels.2019.05.007 | https://github.com/ncbi/scPopCorn | December 28, 2018 |
| scANVI | Semi-supervised variational inference with deep generative models | 10.1101/532895 | https://github.com/chenlingantelope/HarmonizationSCANVI | January 29, 2019 |
| BUSseq | Fits a Bayesian hierarchical model | 10.1101/533372 | https://github.com/songfd2018/BUSseq-0.99.0 | January 29, 2019 |
| FastMNN | PCA + Mutual nearest neighbours | https://marionilab.github.io/FurtherMNN2018/theory/description.html | https://github.com/MarioniLab/scran | June 3, 2019 |

| | | | | |
|---|---|---|---|---|
| scBatch | Sample distance matrix adjustment | 10.1093/bioinformatics/btaa097 | https://github.com/tengfei-emory/scBatch | June 13, 2019 |
| Bermuda | Autoencoder with transfer learning | 10.1186/s13059-019-1764-6 | https://github.com/txWang/BERMUDA | July 2, 2019 |
| SMNN | Supervised mutual nearest neighbors | 10.1101/672261 | https://github.com/yycunc/SMNN | September 20, 2019 |
| BEER | Removal of PCs with batch effects | 10.1038/s41421-019-0114-x | https://github.com/jumphone/BEER | September 24, 2019 |
| trVAE | Conditional variational autoencoder | arXiv:1910.01791 | https://github.com/theislab/trvae | October 4, 2019 |
| MOFA2 | Multi factor analysis model | 10.1101/837104 | https://github.com/bioFAM/MOFA2 | November 9, 2019 |
| scadKNN | Autoencoder (+ KNN classification) | 10.1109/BIBM47256.2019.8982969 | N/A | November 18, 2019 |
| scPhere | Variational autoencoder | 10.1101/853457 | https://github.com/klarman-cell-observatory/scPhere | November 25, 2019 |
| Dmatch | Kernel density matching with external reference | 10.1101/2020.01.05.895136 | https://github.com/qzhan321/Dmatch | January 6, 2020 |
| scDGN | Adversarial networks (cell identity labels required) | 10.1101/2020.01.06.896621 | https://github.com/SongweiGe/scDGN | January 7, 2020 |
| sstGPLVM | Gaussian process latent variable model with t-distributed noise | 10.1101/2020.01.14.906313 | https://github.com/architverma1/sc-manifold-alignment | January 14, 2020 |

| | | | | |
|---|---|---|---|---|
| BATMAN | Minimum weight matching on bipartite graph | 10.1101/2020.01.22.915629 | https://github.com/mandricigor/batman | January 23, 2020 |
| CSS | Represent cells by similarity to clusters in individual samples | 10.1101/2020.02.27.968560 | https://github.com/quadbiolab/simspec | February 28, 2020 |

### Supplementary Table 2: Metrics runs

**Supplementary Table 2: Applicability of metrics to data integration outputs**. Specifically metrics for beyond-label conservation cannot be run on all outputs such as corrected graph outputs and ATAC tasks. The asterisk (*) denotes that no relevant trajectories were found in the ATAC tasks and none were input into the simulation tasks.

| **Metric** | **Graph** | **Embedding** | **Feature** | **RNA** | **ATAC** | **Simulation** |
|---|---|---|---|---|---|---|
| PCR batch | | ✗ | ✗ | ✗ | ✗ | ✗ |
| Batch ASW | | ✗ | ✗ | ✗ | ✗ | ✗ |
| Graph connectivity | ✗ | ✗ | ✗ | ✗ | ✗ | ✗ |
| Graph iLISI | ✗ | ✗ | ✗ | ✗ | ✗ | ✗ |
| kBET | ✗ | ✗ | ✗ | ✗ | ✗ | ✗ |
| Normalized Mutual Information | ✗ | ✗ | ✗ | ✗ | ✗ | ✗ |
| Average Rand Index | ✗ | ✗ | ✗ | ✗ | ✗ | ✗ |
| Cell type ASW | | ✗ | ✗ | ✗ | ✗ | ✗ |
| Graph cLISI | ✗ | ✗ | ✗ | ✗ | ✗ | ✗ |
| Isolated label F1 | ✗ | ✗ | ✗ | ✗ | ✗ | ✗ |
| Isolated label ASW | | ✗ | ✗ | ✗ | ✗ | ✗ |

| | | | | | | |
|---|---|---|---|---|---|---|
| Cell cycle conservation | | ✗ | ✗ | ✗ | | |
| HVG conservation | | | ✗ | ✗ | | ✗ |
| Trajectory conservation | ✗ | ✗ | ✗ | ✗ | * | * |

### Supplementary Note 1: Extending kBET for fair assessment of graph-based integration results

Evaluating how well batch effects are removed in an integration task is complicated by different output formats. Any evaluation metric that can compare graph-based outputs and joint embeddings or corrected feature matrices, must work on the integrated graph (a connectivity matrix). For joint embeddings or corrected feature matrices, such a graph is computed by finding k nearest neighbors based on pairwise distances between cells in the embedding. This process results in a graph where each node has the same out-degree (edges leading outwards). In contrast, a graph-based integration method can output an integration graph with varying *k* per neighborhood. This neighborhood size variance is particularly noticeable in the outputs generated by Conos[3].

We use the kBET[4] metric to assess batch removal for kNN-based outputs. Here, the choice of *k* determines the statistical power of the test per neighborhood. Thus, having a variable *k* means that the rejection of the null hypothesis is less likely in certain regions. As a result, it is important to have a consistent minimal *k* across all tested neighborhoods in all integration outputs. While we can adapt the parameter *k* in our data processing pipeline for methods that output embeddings or corrected feature spaces, it is inherent to the method for graph-based integration methods. In order to benchmark data integration in a consistent manner, we chose to use only the recommended defaults for each method. Thus, we must adapt the input for kBET rather than changing the parameters of the methods that we run to fairly evaluate batch removal across integration output formats.

The output of a graph-based integration method is a graph that encodes the biological signal that is shared across batches. Here, the graph structure, rather than the individual edge, is the important signal. Thus, to increase the number of nearest neighbours we can obtain per cell, we use the local structure in the network to increase the density of the connectivity matrix. Motivated by previous work on diffusion along kNN-graphs in scRNA-seq analysis[5,6], we achieve this by running a diffusion process on the graph. Specifically, we simulate an *N*-step diffusion process where *N* is selected to obtain a minimum of *k* non-zero connectivity per cell. This process is described by the equation:

$$M = \sum_{i=1}^{N} T^{i},$$

where *M* is the diffusion-extended connectivity matrix, and *T* is the row-normalized connectivity matrix.

The above diffusion process is performed at two points in our extended kBET metric. Firstly, we perform graph diffusion on the initial connectivity matrix of graph-based outputs before running kBET. This diffusion run ensures that we have a minimum of *k* nearest neighbors per node. Here, *k* is chosen to match the number of nearest neighbors calculated for other outputs (*k*=50). Secondly, we perform graph diffusion after the connectivity matrix is subsetted to a particular cell identity label. After subsetting, we may obtain multiple connected components in the subsetted graph, especially in poorly integrated datasets. In this setting we first assess which connected components are sufficiently large to evaluate via kBET. A sufficiently large connected component is one with at least 3**k* nodes, where *k* is chosen by the kBET default of the median number of cells per batch within the subsetted data. Note that we enforce minimum and maximum *k* thresholds of 10 and 100. Graph diffusion is performed in all sufficiently large connected components (for all integration outputs) to give a consistent number of nearest neighbors per cell. Cells in connected components that are not sufficiently large are given scores of 1, indicating poor batch integration. Furthermore, cell identity labels where fewer than 75% of cells are in sufficiently large components are given a kBET score of 1 to denote poor batch mixing.

### Supplementary Note 2: Graph LISI extends LISI to graph-based integration results

In order to evaluate batch removal in data integration in a consistent manner, we need metrics that can be applied to all output formats. As corrected expression or accessibility matrices and joint embeddings can both be processed to produce integrated graphs, we specifically require metrics that work on graph structures. The only previously published metric for batch removal that works on graphs is kBET. However, to ensure a robust evaluation of batch removal, it is important to base this assessment on multiple metrics.

Local inverse simpson index (LISI)[7] scores are typically computed on nearest neighbour lists. These neighbour lists are obtained from a kNN graph algorithm computed with *k=90* neighbours. Integrated graph outputs, such as those produced by BBKNN[8] and Conos[3], return integrated graphs often with far fewer neighbours. As these methods do not also output joint embeddings, we cannot simply generate new kNN graphs to produce longer neighborhood lists. Thus, the classical LISI metric cannot be applied to integrated graph outputs.

Here, we extended the classical LISI metric to work on integrated graphs in our *graph LISI* metric. In graph LISI, we replace the distance measurement on joint embeddings with a graph distance to compute large nearest neighbour lists also when nodes only have few nearest neighbours. Specifically, we used Dijkstra's algorithm[9] on the connectivity matrix to compute shortest paths from one cell to all other cells. Thereby, the shortest path length serves as an approximation for the distance on an embedding that is typically used in kNN graph algorithms. As integrated data often form a single, connected graph, such that every cell is connected to all other cells. Using Dijkstra's algorithm, we obtain sufficiently large neighbourhood sizes to compute the LISI for every cell in the largest connected component. In case there are smaller connected components for which we cannot measure graph distances to other cells, these cells belong to an outlier group, which has not been integrated well. Thus, we assign a LISI of 1 to these cells, which reflects the worst possible score. In accordance with the original LISI, we compute the median over all cells to obtain the LISI score. Finally, the LISI score is scaled in two steps to the unit interval (see **Methods**).

On corrected feature matrices (expression or accessibility) and joint embeddings, we construct a kNN graph connectivity matrix via Euclidean distances on the embedded space or on a PCA representation using the compute nearest neighbours function in Scanpy[10] (*sc.pp.neighbors*). Here, we deliberately choose `n_neighbors=15` as a basis for several reasons. Firstly, graph iLISI with `n_neighbors = 15` compared favourably to the original iLISI implementation in contrast with graph LISI using `n_neighbors = 90` on the unintegrated pancreas scenario (data not shown). Secondly, Dijkstra's algorithm runs faster the fewer neighbours are used to create the kNN graph for corrected feature matrices and joint embeddings as the algorithm scales linearly with the number of edges[11]. Thirdly, we want to ensure a fair comparison of all output types. As mentioned above, graph-based outputs tend to have smaller neighbourhood sizes. Thus, we can create similar initial conditions for graph LISI using comparable neighbourhood sizes across output types.

We compared graph iLISI results to the original iLISI on two integration tasks (pancreas and human immune cells, see **Supplementary Fig. 25**). We computed graph iLISI scores on a connectivity matrix with 15 nearest neighbours for corrected feature matrix and joint embedding integration outputs, and on the integrated graph for graph-based outputs. For visualisation of the graph LISI results for graph-based outputs, we used the same scores for both x- and y-axis (as the original iLISI does not apply for integrated graphs). It must be noted that we display iLISI scores after step 1 of the scaling, i.e. the shift by -1, such that the worst possible score is 0. Both scores strongly correlate for non-graph integration outputs (Pearson correlation coefficient for the pancreas task is 0.978 and 0.984 for the immune cell human task). Thus, we conclude that graph LISI is a reasonable metric to assess batch removal (as graph iLISI) and cell type preservation (as graph cLISI), respectively, on graph structures.

### Supplementary Note 3: Detailed analysis of Integration tasks

#### 3.1 Immune cells

For the immune cell atlas, we investigated two separate integration tasks: the first, considering only human samples (n=10); the second, merging human and mouse samples (n=23). In both cases, two tissues were considered: peripheral blood and bone marrow.

##### 3.1.1 Human

In the human immune cell integration task, six challenges can be identified: (1) inter-sample variability arising from the different donors; (2) integration across single-cell protocols (10X and smart-seq2 in Villani's sample); (3) capturing consistent cell populations across tissues of origin; (4) separation of cell subtypes that are transcriptomically similar; (5) preservation of tissue-specific cell annotations as separate clusters; and (6) conservation of the trajectory of erythrocyte development across batches. Challenges (1) and (2) can be solved by removing batch effects across samples and across platforms, respectively, while preserving biological variation. Successfully solving challenge (3) can be achieved by correctly grouping cell types that are found across tissues (e.g., CD8+ and CD4+ T cells, CD20+ B cells, CD14+ and CD16+ monocytes). In challenge (4), we are interested in evaluating whether cell types that share a similar transcriptome (e.g., CD8+ and CD4+ T cells; CD14+ and CD16+ monocytes) can be recapitulated in separate subclusters. Challenge (5) concerns in particular cell annotations that are bone marrow specific, such as monocyte progenitors, erythroid progenitors, erythrocytes and CD10+ B cells. Finally, challenge (6) can be addressed by conserving the trajectory from hematopoietic stem and progenitor cells (HSPCs) via megakaryocyte progenitors and erythroid progenitors, to erythrocytes. It should be noted that we are evaluating the preservation of a global trajectory structure from two points of view: (i) by considering the whole dataset and focusing on the presence/absence of a trajectory which can be visually recognized (**Supplementary Fig. 12**); and (ii) by considering only cell types belonging to the trajectory and assessing whether the cells are placed in a continuum that is consistent with the calculated pseudotime score (**Supplementary Fig. 1-3**). Furthermore, our trajectory metric calculates local conservation of the order of cells in the trajectory per batch compared to unintegrated data.

In the low dimensional embedding plots of the top 4 performing methods (Scanorama (embedding), Conos (unscaled, HVG), Harmony and BBKNN; **Fig. 2 b,c**), all methods appear to have resolved inter-sample and inter-platform batch effects. Moreover, most methods succeeded in capturing consistent cell populations across tissues. Some batch structure remained for Conos, which tended to conserve batch-dependent substructures of CD8+ and CD4+ T cells. Scanorama incorrectly separated plasmacytoid dendritic cells (from Smart-seq2 data from Villani) into two clusters, placing one cluster near plasma cells and the other with dendritic cells. In contrast, BBKNN placed plasma cells, plasmacytoid dendritic cells, and monocyte-derived dendritic cells into a continuum, rather than maintaining a clear separation between the clusters. Nevertheless, the separation of cell subtypes is successfully overcome by the four methods. Particularly, Scanorama and Harmony performed well on this challenge, keeping a clear distinction between CD8+ and CD4+ T cells, and NKT and NK cells. Furthermore, a clear separation between tissue-specific cell types is achieved in all top methods.

The top-performing methods can be distinguished based on trajectory results: while BBKNN and Scanorama conserved the order of cell identity clusters within the trajectory (**Figs. 2 b,c** and **Supplementary Fig. 1,2**), Harmony shows diversity in erythrocyte endpoints but orders progenitor cells in a correct continuum, and Conos (unscale, HVG) does not correctly order the progenitor cell populations. Interestingly, optimal trajectory conservation results are typically obtained when integrating using full gene sets and unscaled data (**Supplementary Fig. 3**). This preprocessing scheme generates the best trajectory conservation in Scanorama (embedding), Conos, and MNN outputs.

We also analysed the poorest preprocessing combinations for the poorest performing methods (Conos (scaled, HVG), Seurat v3, trVAE, and LIGER; **Supplementary Fig. 12**) to evaluate the result of poor data integration. Here, trVAE failed to integrate data between 10X and Smart-seq2 data, whereas the other three methods succeeded. Furthermore, trVAE tended to overcorrect the data, losing the separation between different T cells and NK/NKT cells. In contrast, Seurat v3 successfully removed inter-individual and inter-platform variability and preserved the phenotypic transition between similar cell types. The shortcomings of the worst-performing Seurat v3 approach lie in the conservation of fine-grained biological variation. For example, challenge (3) was only partially solved, since monocyte-derived dendritic cells are overlapping with other cell types. In addition, Seurat v3 failed to preserve tissue-specific cell types such as CD10+ B cells. LIGER, which successfully integrated cross-platform batch effects, instead failed to preserve cell types shared between tissues such as CD20+ B cells and CD4+/CD8+ T cells (challenges (3) and (4)). Likewise, LIGER output contained overlapping erythrocytes, CD14+ monocytes, and CD4+ T cells, completely losing their identity. Similarly, Conos (scaled, HVG) produced an integrated dataset, which, despite integrating batch effects, failed to preserve cell type identity for CD10+ B cells, erythroid progenitors and erythrocytes. Furthermore, none of the poorest performing methods, but trVAE, conserved the trajectory of erythrocyte development (**Supplementary Fig. 1**).

##### 3.1.2 Human and mouse

Integrating mouse and human samples adds a higher level task to the challenges that characterize human samples alone: cross-species integration. In particular, this translates into two separate challenges: (1) cross-species integration inside the same tissue of origin; and (2) cross-species integration between tissues. Furthermore, we are interested in assessing if the methods are able to identify, in the low dimensional embedding plots (**Supplementary Fig. 13, 26**), a cross-species trajectory of erythrocyte differentiation. As in the case of human samples

alone, we can also evaluate integration success by removal of sample and protocol batch effects while preserving tissue-specific cell identities and cell subtypes.
Considering the embedding plots of the top four performing methods (Scanorama, scVI, BBKNN (unscaled, HVG) and ComBat; **Supplementary Fig. 13**), it is striking that none of the methods successfully overcame the batch effect derived from the two species. However, we observed a partially successful integration occurring in specific cases: (i) Scanorama resolved cross-species and cross-tissues batch effects for NK cells; (ii) scVI correctly placed B cells, NK cells, CD8+ T cells, monocytes and erythrocytes in adjacent, partially overlapping clusters, which are shared across tissues and species; (iii) BBKNN placed a human cluster of erythroid progenitors and erythrocytes in close proximity to the mouse counterpart; and (iv) BBKNN and ComBat both placed human B cells adjacent to their mouse counterparts. This indicates that, despite the strong batch effect, these methods were able to capture similar gene expression profiles for cell populations that are shared across species. On the other hand, all top performing methods showed successful batch effect removal for human samples. On the mouse data, all methods, apart from scVI, failed to integrate the two bone marrow studies (Dahlin and MCA), with a clear separation still visible for shared cell types such as neutrophils. This separation of mouse bone marrow studies also affected our trajectory analysis: although scVI, Scanorama, BBKNN, and ComBat preserved the mouse differentiation trajectory in terms of cell placement, a cluster of MCA erythrocytes was separated from the main trajectory in the latter three integration outputs. Moreover, when considering only the cell subset of erythroid differentiation (**Supplementary Fig. 26**), none of the methods were able to reconstruct pseudotime across species. Three methods (Scanorama, scVI, and BBKNN) could however partially maintain a separate ordering of cell labels for human and mouse cells (with BBKNN being successful only for human cells). Interestingly, also the best trajectory conservation results only contained a separated reconstruction of the pseudotime trajectory by species (**Supplementary Fig. 5**). Specifically, all methods were able to preserve pseudotime and cell type placement for the mouse cells, while the human trajectory, with fewer cells, proved more challenging.
A different scenario is depicted by the four poorest-performing methods: BBKNN (scaled, full features), Conos, Harmony, and LIGER (**Supplementary Fig. 13**). While Conos overcorrected the data, removing batch variation due to species but also biological cell type variation, Harmony, BBKNN, and LIGER represent interesting cases. Harmony exhibited a similar behavior to the previously described best performing methods: despite failing the cross-species integration, it placed human CD8+ T cells and B cells in the vicinity of their mouse counterparts in a subset of batches; however, cell types such as CD8+ T cells, CD4+ T cells, NKT and NK cells were separated into multiple clusters, even when belonging to the same donor (e.g., 10X sample). This represents an inconsistent integration output, even within the same tissue (e.g., mouse bone marrow, where cells were separated by study and protocol), which is particularly difficult to handle on unseen data. Nevertheless, mouse peripheral blood and mouse bone marrow are partially integrated, when belonging to the same study (MCA). BBKNN generated a highly connected integration output, which correctly integrated some cell types shared between tissues and species (e.g., B cells, CD4+ and CD8+ T cells, monocytes and NK/NKT cells), but merged others even inside the same tissue (e.g., basophils and megakaryocyte progenitors in mouse bone marrow). LIGER represents a particularly interesting case, being the only method

which consistently integrated across species and other batch effects while conserving broad cell type structure (e.g., erythrocytes, B cells, NKT and NK cells, monocytes). However, LIGER also removed biological variation between cell types, especially transcriptomically similar cell populations (e.g., CD4+ and CD8+ T cells). Indeed, even the conserved broad cell type clusters were heterogeneous, merging distinct cell types (e.g., neutrophils and monocytes). Moreover, smaller clusters, such as plasmacytoid dendritic cells and basophils, can no longer be detected. Finally, trajectory structure is generally poorly conserved across the bottom performing methods. BBKNN and Harmony partially succeeded at reconstructing the ordering of cells (**Supplementary Fig. 26**), yet especially for Harmony this is not well represented in the global placement of clusters in a UMAP.

#### 3.2 Simulation 1

Simulation 1 consists of six batches designed to replicate an experiment consisting of multiple samples from a single tissue (with seven cell types), produced using different technologies. This simulation presents several challenges for integration. The simulated batches differ in number of cells (1000 - 3000), cell type proportions (0 - 35%) and counts per cell (30 - 100% of baseline). Integration methods must attempt to remove the technical differences between batches while maintaining differences between cell types and retaining cell types that are only present in some batches.

Most of the methods performed well on this task, resulting in embeddings that showed distinct clusters by cell type but little evidence of separation between batches (**Supplementary Fig. 14**). This extended down to the worst performing method BBKNN (scaled/full feature). In general methods were able to improve batch correction without a significant loss of bio-conservation (and in several cases an improvement) (**Supplementary Fig. 19**). Group 7 represented a rare cell type that is only present in two of the six batches at low proportion. While Seurat v3 and Scanorama placed this group close to another cell type they were still able to be separated. The worst performing methods failed in different ways. While the trVAE embedding didn't clearly separate cell types they could still be distinguished with the exception of Group 7 which is mixed with other cell types. In contrast, MNN (unscaled/full feature) undercorrected on this task. The embedding showed clear separation between cell types but within those groups the different batches could still clearly be distinguished. The scVI (unscaled/full feature) embedding also showed undercorrection with multiple clusters for some cell types and separation between batches within multiple clusters. Interestingly MNN (unscaled/HVG) was one of the best performers, suggesting that the undercorrection may be a result of including additional features in the integration.

#### 3.3 Simulation 2

Simulation 2 is designed to replicate a more complex experiment with a nested design and includes four batches, each of which has three subbatches. This design is analogous to a multi-center experiment where each center processes multiple batches (possibly using different technologies). In this scenario the between center batch effect (batch) can be expected to be larger than the batch effect between samples from the same center (subbatch). The extra level of variation presents a challenge for integration methods which must remove the batch and subbatch effects while retaining differences between cell types. The number of groups in Simulation 2 has been reduced to four.

As would be expected given the more complex scenario we observed a greater spread of performance on Simulation 2 compared to Simulation 1 (**Supplementary Fig. 19**). The top performing methods (Seurat v3, Harmony) were able to improve both batch correction and bio-conservation compared to the unintegrated dataset. The scaled/full feature version of Conos received the best bio-conservation score at the cost of a slightly worse batch correction score. All three of these methods produced embeddings with clearly separated cell types, however some separation of subbatches within these groups is still visible (**Supplementary Fig. 15**). In this scenario the worst performing methods tended towards undercorrection and were unable to remove either the batch or subbatch effects. These methods received low scores for both batch correction and bio-conservation. The embeddings show partial integration where cell type groups are nearby each other but not fully merged resulting in regions of the cell type but with distinct clusters by subbatch. Encouragingly, even the worst-performing methods did not overcorrect the data such that different cell types were merged together. The scVI result (unscaled/full feature) presents an exception to this rule: it overcorrected, removing much of the separation between cell types.

#### 3.4 Pancreas

The human pancreas task consists of nine batches from six datasets. We have several challenges in the dataset: Firstly, we integrated different experimental protocols with varying sequencing depth. Data from the CEL-seq and CEL-seq2 are UMI counts, which were converted to transcript numbers through binomial statistics. Thus, the resulting values can be considered as UMI-count-like. inDrop is a UMI-based 3' biased protocol, SMARTer is a full-length protocol and was already RPKM-normalised, while SMART-Seq2 and Fluidigm C1 are full-length and highly sensitive protocols, which do not contain UMIs. Secondly, we have a nested batch effect as we consider four different donors from the inDrop dataset as separate batches, while all other datasets are treated as single batches. Thirdly, the datasets differ in data complexity, ranging from four major endocrine cell types in the SMARTer dataset to 14 different cell types in the inDrop dataset. In addition, T cells were only found in the inDrop dataset. We distinguished two subtypes of stellate cells (activated and quiescent), which should ideally be placed in close proximity to one another, but should not overlap. Likewise, the immune cell types (mast cells, macrophages, and T cells) should be placed in close proximity in the embedding plots. This collection of datasets was used in several data integration method publications to benchmark methods, which helps the reader to compare our results to the respective original literature[3,7,12–15].

Overall, the top performing methods integrated all batches correctly, accounting for both nested batch effects and different scales of the protocols, while separating cell types (see

**Supplementary Fig. 9,16**). Seurat v3 and Conos were not affected by the nested batch structure, as all cells were evenly distributed within each cell type. In contrast, BBKNN and Scanorama (embedding, scaled/HVG) showed patches or bands of cells from the same batch, indicating an incomplete removal of the batch effect. For example, alpha cells in BBKNN and Scanorama (embedding) integration results contained batch substructure separating SMARTER cells from the remaining alpha cells in Scanorama (embedding), while the inDrop batches separated slightly from the other batches in BBKNN. We observed a similar grouping of the batches in beta cells. Thus, Conos and Seurat v3 visually corrected the nested batch effect better than BBKNN and Scanorama (embedding).

Examining the distribution of rare cell types (e.g., epsilon cells, T cells, macrophages, mast cells and Schwann cells), we observed several differences across the top performing methods. For instance, epsilon cells were partially merged with quiescent stellate cells in Seurat v3 and have submerged in the Conos FA plot. In contrast, BBKNN and scanorama (embedding) separated the cells from other cell types. The subtypes of stellate cells partially overlapped in Seurat v3 (also with Schwann cells), while they are placed in close proximity but separately in all other top performing methods.

Interestingly, Seurat v3, BBKNN, and Conos indicated transition states (*e.g.*, between alpha and ductal cells in Seurat v3 and delta and beta cells in BBKNN) that were not present in the unintegrated datasets, nor have these transitions been reported in the literature. Therefore, the indicated transitions are spurious and a result of mild overcorrection. Only the Scanorama embedding data integration showed clearly distinct cell types in the UMAP plot, which matches its high bio-conservation scores (**Supplementary Fig. 16** and **Supplementary Data 3**). In general, Seurat v3 and Conos removed the nested batch effects, while conserving most of the strong biological signal in the pancreas datasets. BBKNN and Scanorama (embedding) accounted less well for the nested batch effect structure but better conserved rare cell types.

Interestingly, the corrected feature matrix of Scanorama (unscaled/full feature) was among the worst-performing methods, being only slightly better than unintegrated data. Specifically, inDrop donors, fluidigmC1 and SMARTer, CEL-seq and CEL-seq2 were integrated, respectively, in separate clusters for each cell type. In the LIGER UMAP plots we observed a good batch effect removal and a broad conservation of major cell types (e.g., alpha, beta, gamma and delta cells), however these cell types mix in the embedding. Furthermore, rare cell types were merged and no longer detectable in the LIGER UMAP plot. trVAE removed the batch effect in the inDrop donors, but failed to do so in other datasets (fluidigmC1, SMARTer, SMARTSeq2); although the CEL-seq and CELseq2 datasets were placed in close proximity for each cell type, they form separate clusters nonetheless. MNN distinctly undercorrected the batch effect in the data: while cell types are proximal across batches, neither donors from inDrop datasets nor different protocols were integrated fully.

#### 3.5 Lung atlas

The lung atlas integration task consists of three datasets taken from a single publication[16]. These datasets consist of 10X and Drop-seq data from lung transplants and biopsies. There are five particular challenges of the lung atlas integration task. These challenges encompass: (1)

Inter-individual variation between human donors, (2) Integration of drop-seq (*ASKXXX* donor IDs) and 10X data (*ARMSXXX* and *XXXB/C* donor IDs), (3) separation of neutrophil and basal cell subtypes with specific annotation, (4) detection of rare cell types shared by few donors (ionocytes), and (5) integration across sampling types and locations. Where challenges (1), (2), (3), and (4) are solved by removing all batch effects pertaining to donor and protocol variation while retaining detailed biological variation, it is more difficult to determine success for challenge (5). Specifically, donors with IDs *XXXB/C* and *ASKXXX* (where *X* denotes a digit) were obtained from lung transplants and tissue resections, whereas *ARMSXXX* donors were sampled via biopsies. While transplant samples typically probe the lung parenchyma, biopsies probe the airways. These sampling protocols result in cell type composition differences. For example, while biopsy samples contain basal 1, basal 2, ciliated and secretory cells, these are either absent (basal) or only present as minor cell populations in transplant donor data. Furthermore, while there are several cell type annotations that are present across samples (ciliated, secretory, endothelium, dendritic cells, and macrophages), these cell types can differ between sampling locations. Especially in secretory and endothelium cells it is expected that spatial location affects the transcriptome to make these cell types distinct between biopsy and transplant samples. Secretory cells from biopsy samples were originally labeled specifically as club cells, where transplant secretory cells contained no higher resolution annotation. Moreover, endothelial cells from lung parenchyma (transplant donors) will be predominantly respiratory endothelial cells that are involved in air exchange, while endothelial cells from airway walls have no such function. Thus, integration of secretory or endothelial cells into a single cluster represents a removal of biological signal. Yet, removing this signal may be intended if a low resolution overview of the data across batches is preferred. This overview may be preferable for tasks such as cell annotation transfer.

The top 4 performers in this integration task were Conos, BBKNN, Scanorama, and scVI (**Supplementary Fig. 10)**. These methods generally succeeded in integrating Drop-seq and 10X datasets (**Supplementary Fig. 17)**. Scanorama and scVI performed particularly well in this regard, whereas Conos and BBKNN still showed a separation by 10x and Drop-seq datasets within macrophages.

A central aspect that differentiates the top performing methods was the merging and preservation of cell type information from challenges (3), (4), and (5). While all top performing methods preserved basal cell subtypes, neutrophil subtypes were merged by Conos and BBKNN. Scanorama retained some visual substructure between neutrophil subtypes but did not preserve the differences shown in the unintegrated case, whereas scVI separated neutrophil subtypes but merged IL1R2 neutrophils with the dendritic cell cluster. Distinguishing neutrophil subtypes is particularly challenging for data integration methods as these subtypes are predominantly present in exclusive donors. A similar separation of top 4 method performance is found considering ionocytes: while Scanorama and scVI retained a separate ionocyte cluster, this visual separation was not found in BBKNN embeddings, and only vaguely detectable in the Conos output. Considering cell identities that are shared between sampling locations, we found that no high performing integration method merged secretory cells from airway samples and tissue resections. While this may reflect negatively in our metrics, it also suggests a sensitivity to secretory cell subtypes. Overall, Scanorama tended to maintain the sampling location signal also in ciliated, endothelium, and dendritic cells, while the latter 3 cell types were merged by Conos, BBKNN, and scVI. Especially in endothelium cells, this can be regarded as removal of biological variation. In contrast, macrophages were best integrated by Scanorama. These cells are predominantly found in transplant samples and thus spatial location played a lesser role here. Conos and BBKNN also integrated these cells across donors, however the dataset substructure is still visible here (10X vs Drop-seq separation). Interestingly, scVI integrated macrophages well across platforms, but exhibited substructure in the macrophage population that did not exist in the unintegrated data; specifically separating a subset of macrophages mostly from a single donor (298C).

Poor integration performance varied between methods. While LIGER strongly overcorrected the data by mixing cell types and batches (removing most of the biological variation in the data), the poorest performing Conos, Seurat v3 and Scanorama gene results still showed distinguishable cell types. Conos and Seurat v3 both over-integrated the data: Conos created a strongly connected embedding in which most cell types were overlapping and neutrophil subtypes and ionocytes are indistinguishable; and Seurat v3 merged basal 1 and basal 2 cells, and even alveolar type 1 and type 2 cells (otherwise noticeably distinct populations). Moreover, Seurat v3 merged secretory and endothelium cells, ignoring differences in spatial location. Interestingly, B cells, which were mainly from a single donor (ASK454), exhibited substructure within this batch in the unintegrated data, but were merged by Seurat v3. This suggests that Seurat v3 performed a stronger dimensionality reduction than other methods and thus also merged signals that may be variable within a single batch. Indeed, the original annotations were also generated via a Seurat analysis pipeline[16]. The Scanorama gene run on full features performed poorly as it failed to integrate any of the three datasets (the dominant nested batch effect). It also separated small clusters of data points from the rest, likely due to fitting of spurious signals in the full gene set data.

#### 3.6 Mouse brain (RNA)

The mouse brain RNA integration task consists of 4 datasets produced using different protocols. The particular challenge of this task is its size, since we have almost 1 mio. cells to integrate. Due to its size, we omitted the kBET metric for this task, as it did not scale to datasets of this size. Furthermore, mouse brain data is captured across spatial locations, and consists of both single-cell and single-nucleus RNA-seq data. While we evaluated biological label conservation only on the broad cell type labels with our metrics (**Supplementary Fig. 11**; note label-free conservation was also measured), we also investigated the spatial arrangement of cells, especially as specific subtypes of neurons, for instance, are restricted to certain regions in the brain (**Supplementary Data 3**). Furthermore, the dataset from Zeisel *et al.*[17] profiled, as the only study, cells from the Pons (PO) and the hypothalamus (HTH), while Saunders *et al.*[18], as the only study, profiled the brain regions Entopeduncular Nucleus (ENT), Globus pallidus and nucleus basalis (GP), and distinguished frontal and posterior cortex. The other two studies only provided the label ‘cortex’ (CTX). It must be noted that the spatial information in the Rosenberg *et al.*[1] dataset was inferred based on marker gene expression, which we opted to label as unknown instead (66,648 cells). Nonetheless, an ideal data integration method would remove the batch effect and integrate the cell types resolved by their location in the brain. Here, we combined three pieces of information (dataset, cell type, and location) to visually assess the quality of integration (**Supplementary Fig. 18**). We focused on the following aspects: first, we examined how all datasets integrate and if we observe an integration of single-nucleus RNA-seq (Rosenberg dataset) and scRNA-seq datasets; next, we examined how well cell types integrated and whether rare and abundant cell types could still be distinguished; likewise, we examined unexpected spurious connections, for example, transitions from neurons to endothelial cells; finally, we considered the spatial substructure within a cell type to check if cell types match spatially (e.g., cerebellar astrocytes, neurons from the hippocampus, and neurons from the cortex).

The best performing methods BBKNN and Combat integrated the Saunders and Zeisel datasets well. However, BBKNN failed to integrate the Rosenberg single-nucleus RNA-seq dataset, while Combat integrated Rosenberg only partially. We observed differences in the integration for different cell types. ComBat integrated endothelial cells and brain pericytes from all datasets, while other cell types of the Tabula Muris[19] and the Rosenberg dataset (e.g., oligodendrocytes, oligodendrocyte precursor cells and astrocytes) remained separated. Thus, we focused on the integration of Saunders and Zeisel datasets in the following. Both BBKNN and ComBat integrated rare cell types (e.g., ependymal cells, microglial cells and macrophages) well and oligodendrocytes and oligodendrocyte precursors partially well (placed in close proximity and overlap slightly). We observed different clusters of oligodendrocytes originating from different locations in the brain. In BBKNN, we observed two main clusters, one consisting of cells from the thalamus, hippocampus and the GP, the other consisting of cells from pons (PO), spinal cord (SC), and midbrain (MB). The latter regions have only been profiled in Rosenberg and Zeisel *et al.*, which in turn did not integrate well in this setting. However, the distinction of two major oligodendrocyte clusters may indicate regional differences of the cells. Such a substructure was not shown in the ComBat corrected data: while the complex neuronal structure was preserved in general, there was little overlap with respect to regions. For instance, neither cortical nor hippocampal neurons from Zeisel and Saunders overlapped in the UMAP. For astrocytes, the overlap for Zeisel and Saunders was better, however certain subtypes (e.g., cerebellar astrocytes) were incorrectly separated. Brain pericytes and endothelial cells showed several subclusters based on their location. Here, we observed that endothelial cells and brain pericytes clustered together across locations and datasets. Based on the function and location close to blood vessels, these two cell types have highly similar transcriptional profiles, and the influence of the spatial location within the brain is less pronounced. Overall, although BBKNN did not integrate snRNA-seq (Rosenberg) and scRNAseq data well, two datasets were integrated while the spatial information of the cell types was preserved. Combat integrated the snRNA-seq partially, but the spatial information is less preserved compared to BBKNN.

While scVI and Scanorama (using highly variable genes, scaled data, and embedding) successfully integrated three out of four datasets, some the cell types partially overlapped. Both scVI and Scanorama placed rare cell types as separate clusters, in which all datasets are well mixed (except for microglial cells from Tabula Muris, which cluster separately from the other datasets). In scVI, all cell types from Tabula Muris were placed in close proximity to the corresponding cell types from the other datasets. Both scVI and Scanorama showed transitions between otherwise unrelated cell types: microglia and neurons or endothelial cells and neurons were connected. Cell types where transitions are expected (e.g., oligodendrocytes and oligodendrocyte precursor cells) were conversely only connected as expected by scVI. Concerning spatial information, all regions mixed equally in both methods, and neuronal brain regions such as cortex (CTX) and striatum (STR) were correctly integrated. Overall, scVI preserved more of the rare cell type information than Scanorama. We conclude that the batch effect across datasets, especially from the single-nucleus RNA-seq protocol, is stronger than the spatial signal.

Among the worst performing integration runs, LIGER and Conos integrated all datasets, but neither rare cell types nor rare regions (e.g., olfactory bulb and olfactory ensheathing cells, macrophages and microglial cells) could be discerned. Furthermore, all populations overlapped in the center of the plot. Here, the plots show transitions between otherwise unrelated cell types. For instance, LIGER connected astrocytes and neurons or endothelial cells and neurons. Concerning spatial information, all regions except the cerebellum (CB) were mixed. Thus, LIGER corrected the bias of snRNA-seq and scRNA-seq protocols, but largely removed cell type and spatial information. While the poorest version of the Conos integration results integrated all datasets and coarsely accounted for the cell type separation, all cells were connected into a single, large component. Indeed, several subclusters connected unrelated cell populations (e.g., neurons and macrophages, astrocytes and endothelial cells, and microglial cells, neurons and astrocytes). Furthermore, oligodendrocyte precursor cells were split into two separate clusters (the cells from Rosenberg did not integrate with the datasets from Zeisel and Saunders for this population). Thus, Conos tended to mismatch cell populations. Scanorama (full feature/corrected feature matrix) failed to integrate the Rosenberg dataset, while integrating Tabula Muris partially and the two datasets Saunders and Zeisel well. In contrast to Scanorama, Harmony integrated the Rosenberg dataset partially (e.g., astrocytes, oligodendrocytes, ependymal cells, and endothelial cells), but failed to consistently integrate Zeisel and Saunders datasets. Furthermore, Scanorama (full feature) separated the rare cell types clearly while Harmony created partial overlaps. Information on brain regions was preserved in Scanorama and Harmony, in which cells from the same region per dataset clustered together. Upon closer inspection, Scanorama preserved the spatial substructure slightly better than Harmony. For example, cortical neurons did not integrate in a single cluster in Harmony, while they partially overlapped in the Scanorama plot. Overall, the poorest performing methods integrated snRNA-seq data fully (LIGER, Conos), partially (Harmony) or not at all (Scanorama gene), similar to the top performers. However, the cell type variation was strongly sacrificed with increasing batch correction.

Overall, none of the methods created an ideal data integration and the distance from top 4 to bottom 4 methods was less obvious than in other tasks. In particular, when spatial information was preserved, snRNA-seq was not integrated well with scRNA-seq. Vice versa, when all datasets were successfully integrated, the spatial distribution of the cells was lost. The batch effect (i.e., protocol differences across datasets) was dominant in neurons, while less apparent in the rare cell types. A possible explanation for this effect may be the experimental handling of the cells. For example, endothelial cells and brain pericytes are small and approximately round and therefore easy to handle once they are isolated. In contrast, neurons are relatively large, fragile and have complex shapes (long, branched dendrites and axons). In the single-nucleus protocol, cell size was not a limiting factor as only nuclei were extracted from the cells. Saunders *et al.*[18] used Drop-Seq, while both Zeisel and Tabula Muris datasets were created with the 10X Genomics protocol. Here, size limitations may play a role in the final data quality. Therefore, partial integration of snRNA-seq and scRNA-seq happened on non-neuronal cell types. Interestingly, scVI and scanorama (hvg, embedding) were the only methods where we observed a successful integration of neurons of the cortex and striatum. Furthermore, BBKNN was ranked highly in particular for batch effect removal, while UMAPs indicated poorer performance than the metrics suggested (in particular for the Rosenberg snRNA-seq dataset). A contributing factor may be the lack of the kBET metric on this task, which only left 2 batch removal metrics that could be calculated for graph-based outputs. Thus, BBKNN batch removal results in particular are likely to be less robust for the mouse brain RNA task than for other tasks. We conclude that the mouse brain integration task was a particularly difficult challenge as the batch effect was inhomogeneous within each dataset and, particularly for neurons, overall stronger than the differences due to location of the cells. Thus, focussing on a particular cell type (such as neurons) potentially helps to obtain a cleaner integration, which matches subtypes and spatial locations more accurately.

#### 3.7 Small and large ATAC tasks

The scATAC-seq data consist of 3 datasets. Each dataset was produced using a different protocol: single nucleus ATAC-seq from Fang *et al.*[20], single cell combinatorial indexing from Cusanovich *et al.*[21] and 10X Chromium for the 10X dataset. We generated two different integration tasks from these three datasets by largely changing the relative proportions of cells between datasets to generate a large ATAC task with strongly imbalanced cell populations between datasets (5%:20%:75% for 10x, Cusanovich *et al.* and Fang *et al.*) and a smaller ATAC task with more balanced cell contributions (13%:57%:30% for 10x, Cusanovich *et al.* and Fang *et al.*).

Integration of scATAC-seq datasets posed a clear challenge to the methods developed for the integration of scRNA-seq dataset. Firstly, scATAC-seq data is binary compared to the gene expression data which contains counts of expressed genes. Secondly, scATAC-seq data can be collected for every position in the genome in comparison to scRNA-seq data which is only collected for genes. This poses a challenge on the selection of the features to be used as a basis for data integration. Peaks of open chromatin typically have different (mainly) non-overlapping coordinates per batch and therefore integration based on peaks would prove difficult. For that reason, we use non-overlapping sliding windows (5000bp) as the canonical, unbiased unit for processing open chromatin data and as a basis for data integration. However, the number of windows in the genome is too large to use as a basis for the integration, as the majority of the integration methods do not scale well with the number of considered features. Therefore we selected the top variable windows per batch. This posed a third challenge for integrating scATAC-seq datasets: the more batches/cells to integrate, the less shared highly variable windows there are between them, hindering the integration task.

The number of 5000bp non-overlapping sliding windows in the mouse genome is >500,000. We reduced this number by selecting, per batch, the 150,000 most highly variable windows for both the large and the small integration tasks. When merging batches (after discarding cells which are not covered by at least 500 windows) the number of cells was reduced to 67,612 and 25,960 for the large and small tasks, respectively, and the number of features per task was reduced to 57,447 and 57,070 for the large and small tasks, respectively.

We observed that all integration methods underperform in the batch correction of the three scATAC-seq batches for both the small and large tasks. From visual inspection of the low dimensional embeddings, we observed a preference for integrating the 10X and the Fang *et al.* datasets over the Cusanovich *et al.* dataset (**Supplementary Fig. 22, 23**); only Seurat v3 preferably integrated the Cusanovich *et al.* dataset with the 10x dataset (**Supplementary Fig. 24**) in the large integration task. The 10X and the Fang *et al.* datasets shared a large number of highly informative windows among them (approximately 85% of windows), while the Cusanovich *et al.* dataset shared a much lower number of highly informative windows with the other two datasets (approximately 45%). Also, a simple correlation between the percentage of the sum of open regions per window for the shared regions showed that the original data from 10X is highly correlated to the original data from Fang *et al.* (R = 0.87), while the Cusanovich *et al.* raw data matrix is lowly correlated with the raw 10X dataset (R = 0.18) and the Fang *et al.* dataset (R = 0.13). This likely explains why most methods showed better integration results between 10x and Fang *et al.* datasets. Seurat v3 instead integrated Cusanovich *et al.* with 10X best in the large integration task, and Fang *et al.* with 10X in the small integration task. Thus, Seurat v3 was the only method for which the relative proportion of cells per batch had an effect, since in the large integration task we have a ratio of 5% (10X) : 20% (Cusanovich *et al.*) : 75% (Fang *et al.*) and in the small integration task the ratios are 13% (10x) : 57% (Cusanovich *et al.*) : 30% (Fang *et al.*). This was likely the case because Seurat v3 internally uses different data integration orders when more than two datasets are considered, depending on the size of the datasets.

From the top performing methods (Harmony, ComBat, Seurat v3 and scVI; **Fig. 4** and **Supplementary Fig. 22-24**), Seurat v3 and ComBat showed the least batch correction, with clearly separated batches on the UMAPs. These methods ranked low for batch correction (6th / 7th and 4th / 5th for the small and big tasks, respectively), but instead had some of the highest biological conservation scores. Both methods placed the same cell types from different datasets in close proximity, but clearly preserved separated cell clusters. BBKNN, Harmony and scVI instead were the top methods at batch effect removal. BBKNN outperformed in batch correction but at the expense of a very low biological conservation score, mainly due to the graph cLISI score. This was most likely due to the internal optimization which enforced connections across batches for each cell. Because of that, isolated cell clusters were lost after the BBKNN integration, as can be observed in the respective UMAP (**Figure 4** and **Supplementary Fig. 22-24**). Harmony ranked second for batch correction, placing the same cell types from all three batches in close proximity on the UMAP; it thus achieved a better biological conservation score due to a higher cell type ASW. Finally, scVI showed a good compromise between batch correction and biological conservation (**Fig. 4** and **Supplementary Fig. 22**).

Across the large and small ATAC tasks, five methods consistently performed worse than the unintegrated datasets in the overall score: BBKNN, trVAE, LIGER, Scanorama, and Conos. The low overall performance of BBKNN, discussed above, was due to a very low cLISI score. For the large integration task, both Scanorama and Conos underperformed the unintegrated datasets in both batch correction and biological conservation. In the small task, LIGER and trVAE achieved some moderate batch integration with poor biological conservation. LIGER interestingly positioned microglia from the 10x and the Fang *et al.* datasets with Cerebellar granule cells from the Cusanovich *et al.* dataset, a cell type that was only present in the Cusanovich *et al.* dataset and remained mostly correctly unmerged in other integration methods.

Finally, trVAE and LIGER could not scale up to integrate the large ATAC-seq dataset, and MNN could not integrate the small dataset either.
